# Determining the specificity of Cascade binding, interference, and primed adaptation *in vivo* in the *Escherichia coli* type I-E CRISPR-Cas system

**DOI:** 10.1101/169011

**Authors:** Lauren A. Cooper, Anne M. Stringer, Joseph T. Wade

## Abstract

In CRISPR-Cas immunity systems, short CRISPR RNAs (crRNAs) are bound by CRISPR-associated (Cas) proteins, and these complexes target invading nucleic acid molecules for degradation in a process known as interference. In type I CRISPR-Cas systems, the Cas protein complex that binds DNA is known as Cascade. Association of Cascade with target DNA can also lead to acquisition of new immunity elements in a process known as primed adaptation. Here, we assess the specificity determinants for Cascade-DNA interaction, interference, and primed adaptation *in vivo*, for the type I-E system of *Escherichia coli*. Remarkably, as few as 5 bp of crRNA-DNA are sufficient for association of Cascade with a DNA target. Consequently, a single crRNA promotes Cascade association with numerous off-target sites, and the endogenous *E. coli* crRNAs direct Cascade binding to >100 chromosomal sites. In contrast to the low specificity of Cascade-DNA interactions, >18 bp are required for both interference and primed adaptation. Hence, Cascade binding to sub-optimal, off-target sites is inert. Our data support a model in which initial Cascade association with DNA targets requires only limited sequence complementarity at the crRNA 5□ end, whereas recruitment and/or activation of the Cas3 nuclease, a prerequisite for interference and primed adaptation, requires extensive base-pairing.

**IMPORTANCE:** Many bacterial and archaeal species encode CRISPR-Cas immunity systems that protect against invasion by foreign DNA. In the *Escherichia coli* CRISPR-Cas system, a protein complex, Cascade, binds 61 nt CRISPR RNAs (crRNAs). The Cascade complex is directed to invading DNA molecules through base-pairing between the crRNA and target DNA. This leads to recruitment of the Cas3 nuclease that destroys the invading DNA molecule and promotes acquisition of new immunity elements. We make the first *in vivo* measurements of Cascade binding to DNA targets. Thus, we show that Cascade binding to DNA is highly promiscuous; endogenous *E. coli* crRNAs can direct Cascade binding to >100 chromosomal locations. By contrast, we show that target degradation and acquisition of new immunity elements requires highly specific association of Cascade with DNA, limiting CRISPR-Cas function to the appropriate targets.

## INTRODUCTION

Clustered Regularly Interspaced Short Palindromic Repeats (CRISPR)-Cas (CRISPR-associated) systems are adaptive immune systems found in approximately 40% of bacteria and 90% of archaea [1]. CRISPR-Cas systems are characterized by the presence of CRISPR arrays and Cas proteins. CRISPR arrays are genomic loci that consist of short repetitive sequences (“repeats”), interspaced with short sequences of viral or plasmid origin (“spacers”) [2–5]. Spacers are acquired during a process known as “adaptation”, in which a complex of Cas1 and Cas2 integrates invading DNA into a CRISPR array, effectively immunizing the organism from future assault by the invader [6]. In the archetypal type I-E CRISPR system of *Escherichia coli*, immunity occurs via two processes known as “biogenesis” and “interference”. During biogenesis, a CRISPR array is transcribed, and Cas6e processes the transcript into individual 61 nt CRISPR RNAs (crRNAs) that each include a single 32 nt spacer sequence flanked by partial repeat sequences [7, 8]. Individual crRNAs are then incorporated into Cascade, a protein complex composed of five different Cas proteins (Cse1 (Cas8e)-Cse2_2_-Cas7_6_-Cas5-Cas6e) [7, 9]. During interference, Cascade complexes bind to target DNA sequences known as “protospacers” that are complementary to the crRNA spacer, and are immediately adjacent to a short DNA sequence known as a “Protospacer-Associated Motif” (PAM) [10] that is bound by Cse1 [11, 12]. The crRNA bound by Cascade forms an R-loop with the target DNA, which in turn leads to recruitment of the Cas3 nuclease, DNA cleavage, and elimination of the invader [13–19].

For type I CRISPR-Cas systems, adaptation can occur by two mechanisms: “naïve” and “primed”. Typically, naïve adaptation requires only Cas1 and Cas2 [6, 20]. Primed adaptation, by contrast, requires all Cas proteins and an existing crRNA [21]. The molecular details of primed adaptation are poorly understood. Spacers acquired by primed adaptation correspond to locations on the same DNA molecule as the protospacer [21–24]. Some type I CRISPR-Cas systems acquire spacers preferentially from one strand [21,23,25], whereas others acquire spacers from both strands [24,26,27]. Primed adaptation has been proposed to involve translocation of Cas3 away from the Cascade-bound protospacer [21, 28].

There are conflicting reports on the relationship between interference and primed adaptation. Initially, it was proposed that primed adaptation occurs only when Cascade-protospacer interactions are sub-optimal and cannot lead to interference, e.g. with a sub-optimal PAM, or with mismatches in the PAM-proximal region of the protospacer known as the “seed” [21,23,29– 31]. However, more recent studies have shown that at least some protospacers can lead to both interference and primed adaptation, indicating that the requirements for interference and primed adaptation overlap [24,32,33].

Prior to interference or primed adaptation, Cascade must bind to the target protospacer. This requires an interaction between Cse1 and the PAM, and base-pairing between the crRNA and protospacer DNA [11,12,14]. PAM recognition is required for both Cascade binding and later recruitment and activation of Cas3 [11,13,16,19]. Changes to the optimal PAM weaken Cascade binding to a protospacer [34]. Nonetheless, some sub-optimal PAMs are sufficient for interference, albeit with lower efficiency than the optimal PAM [31]. Sequences within the crRNA spacer are also required for initial binding of Cascade to a protospacer; mutations in positions 1-5 and 7-8 adjacent to the PAM of the protospacer (the seed sequence) reduce the affinity of Cascade for the protospacer [29].

Reports of the sequence determinants for Cascade binding, interference and primed adaptation are conflicting [16,18,23,31,35]. In particular, the impact of extensive mismatches in the crRNA:DNA hybrid on Cascade binding and primed adaptation is unclear [16, 35]. Importantly, association of Cascade with protospacer DNA has not previously been studied in an *in vivo* context. Here, we use ChIP-seq to perform the first *in vivo* assessment of Cascade binding to its DNA targets. Our data show that base-pairing between the crRNA and protospacer with as few as 5 nt in the seed region, coupled with an optimal PAM, is often sufficient for Cascade binding. Hence, crRNAs, including those transcribed from the native *E. coli* CRISPR loci, drive off-target binding at over a hundred chromosomal sites. If Cascade binding to DNA were sufficient for interference or primed adaptation, these off-target binding events would likely be catastrophic for the bacterium [36, 37]. However, we show that extensive base-pairing between the crRNA and protospacer from the PAM-proximal end is required for efficient interference and primed adaptation. Thus, under native conditions, Cascade samples potential DNA target sites, but limits nuclease activity to protospacers that meet a higher specificity threshold that would only be expected of on-target sites.

## RESULTS

### An AAG PAM and seed matches are sufficient for Cascade binding to DNA target sites in vivo

Previous studies of Cascade association with protospacer DNA have been performed *in vitro* using purified Cascade and crRNA. To determine the *in vivo* target specificity of *E. coli* Cascade, we used ChIP-seq to map the association of Cse1-FLAG_3_ and FLAG_3_-Cas5 (FLAG-tagged strains retain CRISPR function; Supplementary Figure 1) across the *E. coli* chromosome in Δ*cas3* (interference-deficient) cells constitutively expressing all other *cas* genes and each of two crRNAs that target either the *lacZ* promoter or the *araB* promoter (both targets are chromosomal; Figure S2A-B). ChIP-seq data for Cse1 and Cas5 were highly correlated (R^2^ values of 0.93-0.99 for *lacZ*-targeting cells, and 0.99 for *araB*-targeting cells), consistent with Cse1 and Cas5 binding DNA together in the context of Cascade. We detected association of Cascade with many genomic loci for each of the two spacers tested (Figure 1A-B; Table S1). In all cases, the genomic region with strongest Cascade association was the on-target site at *lacZ* or *araB*. Off-target binding events occurred with <20% of the ChIP signal of on-target binding. To determine the sequence requirements for off-target Cascade binding with each of the two crRNAs used, we searched for enriched sequence motifs in the Cascade-bound regions, excluding the on-target site (Table S2). For both the *lacZ* and *araB* spacers, the most enriched sequence motif we identified was a close match to the canonical PAM, AAG, on the non-target strand, followed by 5 nt of sequence complementarity at the start of the crRNA seed region (Figure 1C-D; c.f. Figure S2A-B). In some cases, we observed Cascade binding events associated with non-AAG PAMs; however, these sites were more weakly bound, and/or had matches in seed region beyond position 5. For example, Cascade binding events targeted by the *araB* spacer were significantly more likely to have matches at positions 7-9 in cases where the PAM was not AAG (an average of 2.2 out of 3 matches per target sequence if the PAM was not AAG [n = 19], and 1.2 out of 3 matches if the PAM was AAG [n=41]; Fisher’s Exact Test *p* = 0.00005). We conclude that as few as 5 bp in the seed region, together with an AAG PAM, are sufficient for Cascade binding, with additional base-pairing in/near the seed region increasing binding and/or overcoming the need for an AAG PAM.

**Figure 1.**
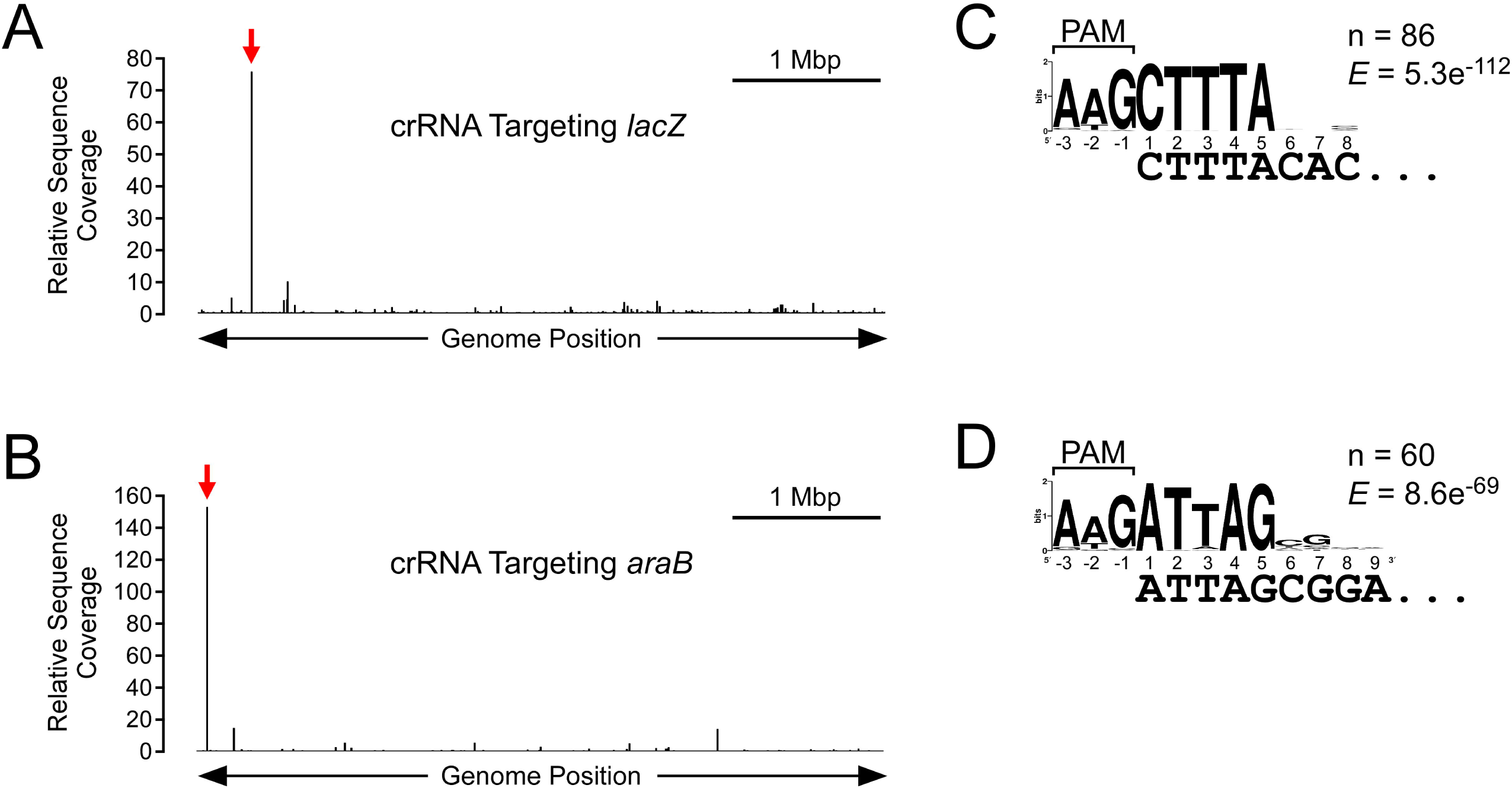
Extensive off-target Cascade binding in *E. coli*. **(A)** Binding profile of Cse1 across the *E. coli* genome, as determined by ChIP-seq, for Cse1-FLAG_3_ cells (AMD543) expressing a plasmid-encoded crRNA targeting the *lacZ* promoter region (pCB380). The graph indicates the relative sequence read coverage (see Methods for details) across the genome in a Cse1 ChIP-enriched sample. A scale-bar is shown for 1 Mbp. The location of the on-target binding site at the *lacZ* promoter is shown by the red arrow. **(B)** Binding profile of Cas5 across the *E. coli* genome, as determined by ChIP-seq, for FLAG_3_-Cas5 cells (AMD554) expressing a plasmid-encoded crRNA targeting the *araB* promoter region (pCB381). The location of the on-target binding site at the *araC* promoter is shown by the red arrow. **(C)** Enriched sequence motif associated with off-target Cascade binding sites when targeting the *lacZ* promoter, as determined by MEME. The likely PAM sequence is indicated. The number of identified motifs and the MEME E-value are shown. **(D)** Enriched sequence motif associated with off-target Cascade binding sites when targeting the *araB* promoter, as determined by MEME. The likely PAM sequence is indicated.

### Extensive off-target Cascade binding driven by endogenous spacers

We identified several sites of Cascade binding that were shared between cells targeting *lacZ* and cells targeting *araB*. These bound regions were not associated with sequences matching the seed regions of either crRNA. We reasoned that these off-target binding events may be due to Cascade association with the endogenous *E. coli* crRNAs. To test this hypothesis, we performed ChIP-seq of Cse1-FLAG_3_, as described above, for cells expressing only the endogenous CRISPR RNAs from their native loci. Thus, we identified 188 binding sites for Cascade (Figure 2A; Table S1). These sites were associated with four enriched sequence motifs, with each motif corresponding to a canonical AAG PAM and 5-10 nt matching the seed region of a crRNA from the CRISPR-I array (spacers #1, #3, #4, and #8; Figure 2B; Figure S2C; Table S2). The strongest binding events were associated with spacer #8 of CRISPR-I (“sp1.8”; Figure 2B; Figure S2C). To confirm that Cascade binding events were due to association with endogenous crRNAs, we repeated the ChIP-seq experiment in cells lacking the CRISPR-I array and in cells lacking the CRISPR-II array. Deletion of CRISPR-II had little effect on the profile of Cascade binding (Figure 2C; Table S1). In contrast, deletion of CRISPR-I resulted in loss of Cascade binding to almost all sites bound in wild-type cells (Figure 2D; Table S1). Instead, low-level binding of Cascade was observed at a small number of sites that were associated with a weakly enriched sequence motif corresponding to a perfect PAM and 8 nt matching the seed region of spacer #2 of CRISPR-II (Figure S2D and Figure S3; Table S2).

**Figure 2.**
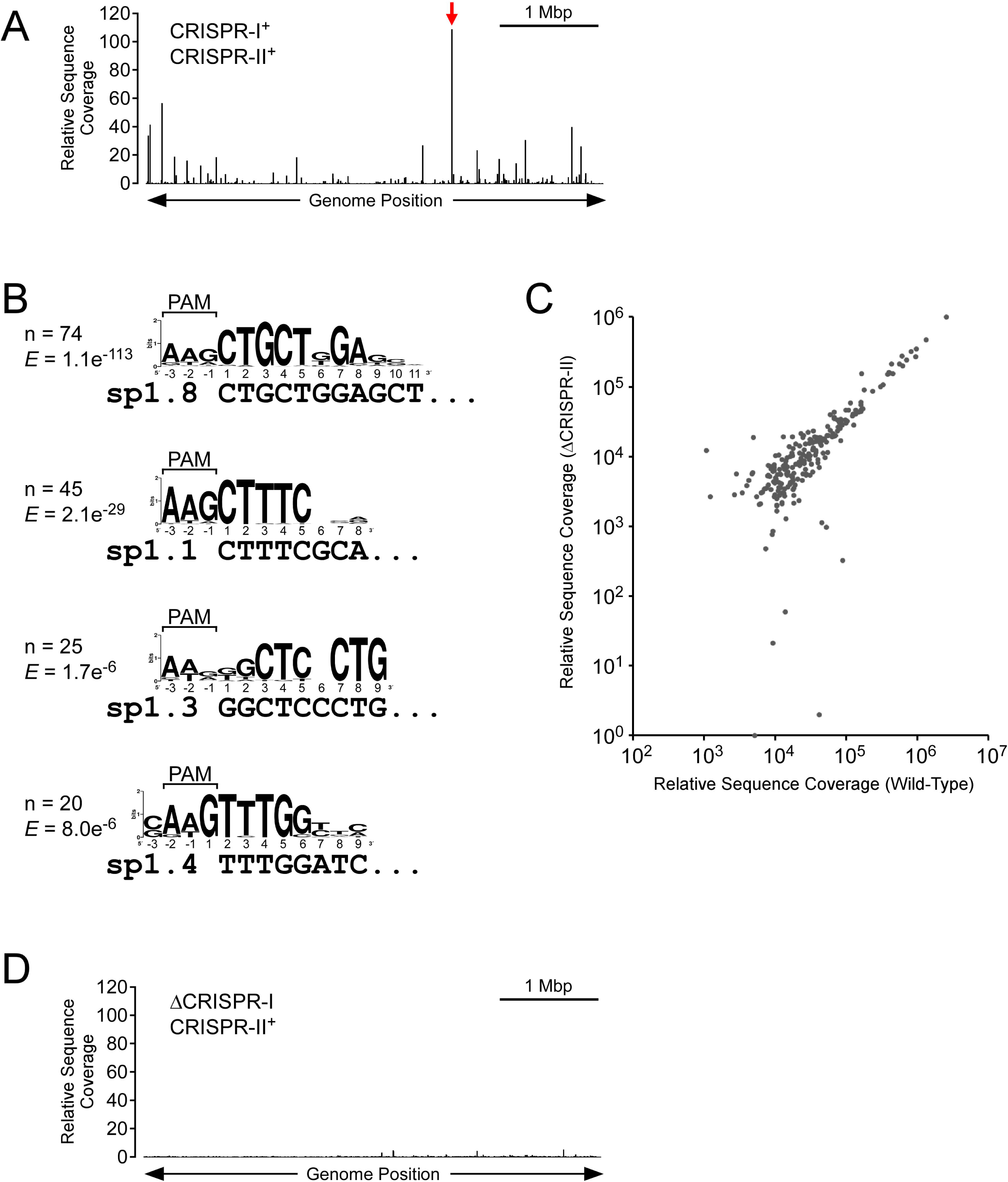
Endogenous crRNAs drive Cascade association with over a hundred chromosomal sites. **(A)** Binding profile of Cse1 across the *E. coli* genome, as determined by ChIP-seq, for Cse1-FLAG_3_ cells (AMD543) expressing only endogenous crRNAs. The location of the binding site within *yggX* is shown by the red arrow. **(B)** Enriched sequence motifs associated with Cascade binding sites in cells expressing only endogenous crRNAs. The four motifs are associated with four of the CRISPR-I spacers, as indicated. The likely PAM sequence is also indicated. The number of identified motifs and the MEME E-value are shown. **(C)** Comparison of Cascade binding events in Cse1-FLAG_3_ cells with both CRISPR arrays intact (AMD543) and or CRISPR-II deleted (LC060). Sequence read coverage is shown for CRISPR-1^+^ CRISPR-2^+^ (AMD543) and CRISPR-I^+^ ΔCRISPR-II cells (LC060), for all ChIP-seq peaks identified for either strain. **(D)** Binding profile of Cse1 across the *E. coli* genome, as determined by ChIP-seq, for Cse1-FLAG_3_ cells expressing only endogenous crRNAs, but with CRISPR-I deleted (LC077).

### CRISPR-I spacer #8 is the major determinant of off-target Cascade binding in cells expressing endogenous crRNAs

Our data suggested that the majority of Cascade binding associated with endogenous crRNAs is due to CRISPR-I, and that the dominant spacer from CRISPR-I is sp1.8. To confirm this, we measured Cascade binding by ChIP-seq in cells lacking CRISPR-I but expressing a plasmid-encoded sp1.8 crRNA. Note that the plasmid-expressed sp1.8 crRNA differs from sp1.8 at the last two nucleotides of the spacer. However, these mismatches are not expected to affect Cascade binding [23, 38]. Most of the Cascade binding sites we observed were identical to those seen in cells expressing both CRISPR arrays, or cells expressing only CRISPR-I (Figure 3A; Table S1), and corresponded to regions containing strong matches to sp1.8 (orange dots in Figure 3A correspond to regions containing a match to the sp1.8 motif shown in Figure 2B). As expected, and unlike for cells expressing CRISPR-I, we detected only a single strongly enriched sequence motif (Figure S4A; Table S2). This motif, as expected, corresponds to an AAG PAM and 9 nt matching the seed region of sp1.8 (Figure S2C). We also detected a weakly enriched sequence motif (Figure S4B; Table S2) that corresponds to an AAG PAM and the 11 nt immediately downstream of the second repeat on the plasmid encoding the sp1.8 crRNA. This is likely due to formation of a non-canonical crRNA that consists of the sequence between the second repeat and the transcription terminator (Figure S2E). A transcription terminator hairpin has previously been shown to function analogously to repeat sequence in the *E. coli* crRNAs [39].

**Figure 3.**
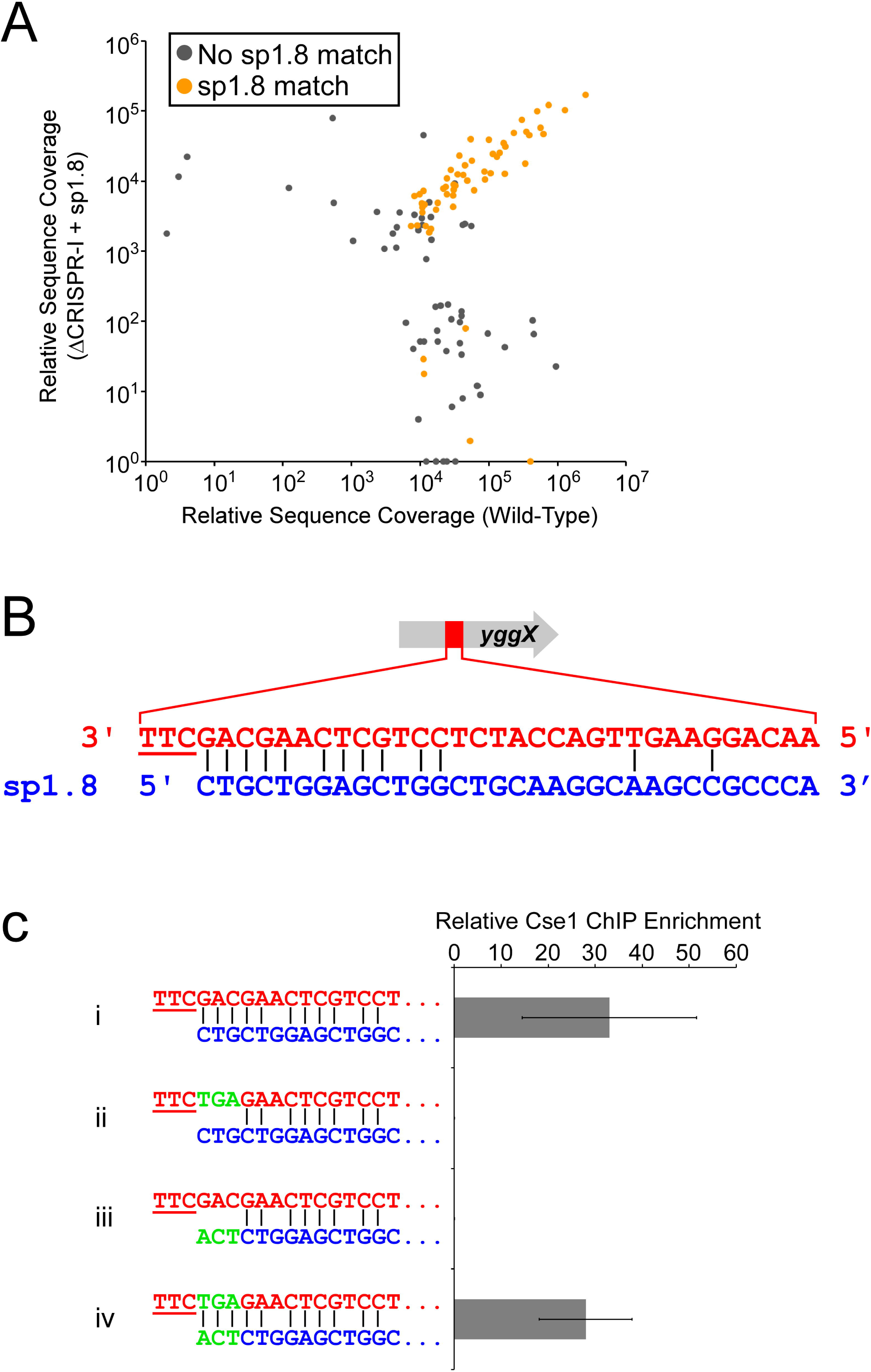
CRISPR-I Spacer #8 is responsible for the majority of Cascade binding in cells expressing only endogenous crRNAs. **(A)** Comparison of Cse1-FLAG_3_ binding events in cells with both CRISPR arrays intact (AMD543), and cells with CRISPR-I deleted (LC077) that express CRISPR-I spacer #8 from a plasmid (pLC008). Sequence read coverage is shown for all ChIP-seq peaks identified for either strain. ChIP-seq peaks associated with the CRISPR-I spacer #8 motif (first motif listed in Figure 2B) are shown in orange. **(B)** Predicted base-pairing interaction between CRISPR-I spacer #8 and a protospacer within *yggX*. The PAM is underlined. **(C)** ChIP-qPCR measurement of Cse1 binding at wild-type (i and iii; AMD566) and mutant (ii and iv; LC099) protospacers in *yggX* for cells expressing wild-type (i and ii; pLC008) or mutant (iii and iv; pLC010) CRISPR-I spacer #8 from a plasmid. The mutations in spacer #8 restore base-pairing potential with the mutant protospacer, as indicated. Values represent the average of three independent replicate experiments. Error bars show one standard deviation from the mean.

The most enriched Cascade target region in cells with CRISPR-I, and cells expressing sp1.8 crRNA, was inside the *yggX* gene. We identified a sequence in this region with an AAG PAM and matches to positions 1-5 and 7-10 of sp1.8 (Figure 3B). We used targeted ChIP-qPCR to measure Cascade binding to this site in cells lacking CRISPR-I but expressing a plasmid-encoded sp1.8 crRNA (mismatches to sp1.8 at the last two nucleotide positions, as described above). We compared binding of Cascade to *yggX* in wild-type cells, and cells where the putative protospacer was mutated in the region predicted to bind the sp1.8 crRNA seed. As expected, we observed greatly reduced Cascade binding at the mutated site relative to the wild-type site. Similarly, we observed greatly reduced Cascade binding at the wild-type site when we expressed a mutant sp1.8 with changes in the seed region (Figure 3C). However, when we combined the mutant spacer with the mutant protospacer, base-pairing potential was restored, and we observed wild-type levels of Cascade binding (Figure 3C). We conclude that sp1.8 is the major determinant for off-target Cascade binding in cells expressing endogenous crRNAs.

### Off-target Cascade binding events do not affect local gene expression

Cascade binding events can lead to transcription repression by preventing initiating RNA polymerase binding to a promoter, or acting as a roadblock to elongating RNA polymerase within a transcription unit [38, 40]. To determine if off-target events driven by endogenous spacers affect local gene expression, we measured global RNA levels using RNA-seq in Δ*cas3* cells with other *cas* genes constitutively expressed, with either intact CRISPR arrays or a ΔCRISPR-I deletion. We detected few differences in RNA levels between the two strains (Table S3), and none of the differences correspond to genes within 1 kb of a Cascade binding site identified by ChIP-seq. We conclude that off-target binding by a Cas3-deficient complex does not impact local gene expression.

### No evidence for RNA-targeting by *E. coli* Cascade

A recent report suggested that Cascade binding to RNAs in *Pseudomonas aeruginosa*, which has a type I-F system, leads to Cas3-mediated degradation of the target RNA [41]. Moreover, this study suggested that only 8 nt of sequence complementarity between the crRNA and target RNA, and a flanking 5□-GGA-3□ sequence, are required to recruit Cas3. This is similar to the sequence requirement for off-target binding to DNA sites (Figures 1 and 2), suggesting that Cascade could target many endogenous RNAs [42]. To determine whether the *E. coli* type I-E CRISPR-Cas system targets RNA in a similar way, we measured global RNA levels using RNA-seq in cells expressing *cas3* from a plasmid and all other *cas* genes from their chromosomal loci, with either intact CRISPR arrays or a ΔCRISPR-I deletion. We compared these data to the data described above for Δ*cas3* cells with either intact CRISPR arrays or a ΔCRISPR-I deletion. We reasoned that targeted RNAs would be less abundant in cells expressing both Cas3 and CRISPR-I. However, we detected only two genes, *ykgE* and *glpD*, for which RNA levels were significantly lower in the *cas3*^+^ CRISPR-I^+^ strain than strains lacking either or both of *cas3* and CRISPR-I (Table S3). Only one of these genes (*glpD*) contains an 8 nt sequence complementary to the 3□ end of a spacer in CRISPR-I (spacer #4; we included the predicted 5’ UTR in the search for both RNAs). Given the length of the two genes, finding an 8 nt match by chance is not unlikely. Moreover, three other genes contain the same 8 nt match to spacer #4, with the same 3 nt flanking sequence, but these genes did not have the RNA profile expected for a Cascade target. Thus, our data strongly suggest that the type I-E CRISPR-Cas system in *E. coli* does not target RNA using a mechanism similar to that described for the type I-F system in *P. aeruginosa*.

### Off-target Cascade binding is not associated with interference

Previous studies have suggested that extensive mismatches at the PAM-proximal end of the spacer/protospacer prevent interference [16, 35]. To determine whether off-target Cascade binding events lead to interference, we constructed a Δ*yggX* Δ*cas3* strain expressing all other *cas* genes, with both CRISPR arrays intact. We introduced a plasmid expressing *cas3*, or an equivalent empty vector. We then transformed these strains with a plasmid containing the off-target protospacer from *yggX* that is an imperfect match to sp1.8, an equivalent plasmid with a protospacer that is a perfect match to sp1.8, a plasmid with a protospacer that is a perfect match to CRISPR-I spacer 2 (“sp1.2”), or an empty vector. We reasoned that the number of viable transformants for plasmids with interference-proficient protospacers would be low for cells expressing Cas3, since interference would cause loss of the protospacer-containing plasmid, leading to killing by the antibiotic selection. In contrast, the number of viable transformants for plasmids with interference-deficient protospacers, or cells not expressing Cas3, should be high. We measured the transformation efficiency for plasmids containing each of the two protospacers in cells with a Cas3-expressing plasmid, or an equivalent empty vector. The efficiency of interference was calculated using the ratio of transformation efficiency for cells with/without Cas3. As expected, the protospacer that perfectly matches sp1.8 resulted in highly efficient interference. Similarly, the protospacer that perfectly matches sp1.2 resulted in highly efficient interference. We conclude that sp1.2 is efficiently assembled into Cascade, despite the lack of chromosomal off-target binding events detected by ChIP-seq. By contrast, the protospacer with the native *yggX* sequence (i.e. imperfect match to sp1.8) resulted in no detectable interference (Figure 4A). We conclude that off-target Cascade binding events do not cause interference.

**Figure 4.**
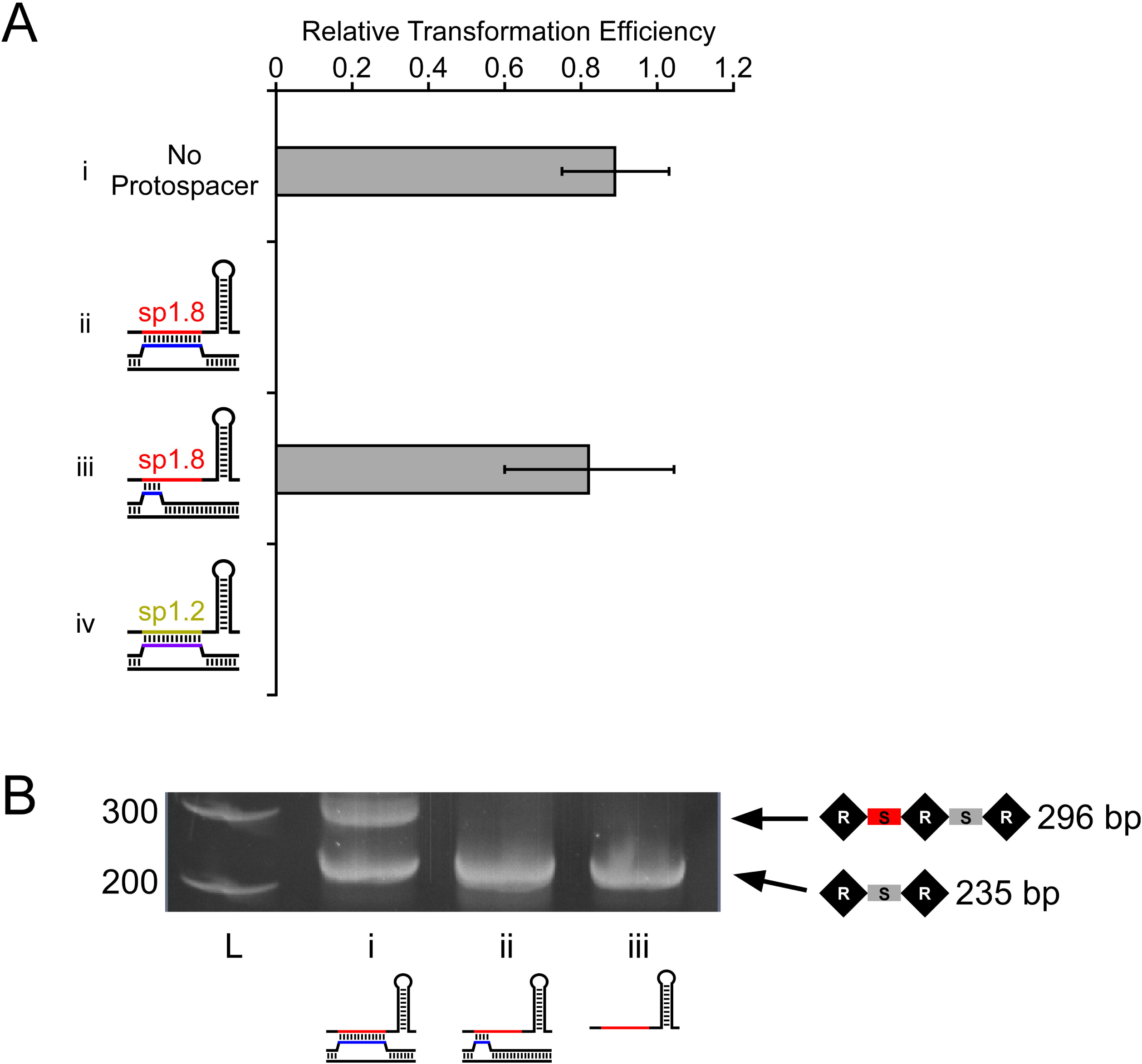
Off-target Cascade binding events are not associated with interference or primed adaptation. **(A)** Relative efficiency of transformation of a *cas3*-expressing plasmid (pAMD191) into LC103 cells expressing spacer #8 from a native CRISPR-I array, and containing (i) empty pBAD24 (“No Protospacer”), (ii) a plasmid with a protospacer that base-pairs perfectly with spacer #8 (pLC022), (iii) a plasmid with a protospacer that has only partial base-pairing with CRISPR-I spacer #8 (pLC021; protospacer sequence matches the off-target Cascade binding site in *yggX*), or (iv) a plasmid with a protospacer that base-pairs perfectly with CRISPR-I spacer #2 (pLC057). Note that crRNAs were expressed from the chromosome since both CRISPR arrays are intact in these strains. Transformation efficiency was calculated relative to that of empty pBAD33, as described in the Methods. Values represent the average of three independent replicate experiments. Error bars show one standard deviation from the mean. The calculated transformation efficiency for protospacers (ii) and (iv) was 0, but the limit of detection in this assay was 3e^-5^. **(B)** PCR-amplification of the start of the CRISPR-II array to detect primed adaptation in cells expressing CRISPR-I spacer #8 (AMD536), *cas3* (pAMD191), and with (i) a protospacer that base-pairs perfectly with CRISPR-I spacer #8 (pLC022), (ii) the protospacer from *yggX* that has only partial base-pairing with CRISPR-I spacer #8 (pLC021), or (iii) empty vector (pBAD24). L = molecular weight ladder, with marker sizes (bp) indicated. The expected PCR product sizes are indicated. Note that there is a non-specific PCR product of ∼380 bp for both samples i and ii (indicated by the asterisks).

### Off-target Cascade binding is not associated with primed adaptation

Protospacers with multiple mismatches to a crRNA can still cause primed adaptation [23], and a recent study concluded Cascade can bind to a protospacer with extensive mismatches, including in the seed region or at the PAM-distal end, and that these binding events cause primed adaptation [35]. To test whether off-target Cascade binding is sufficient for primed adaptation, we used the strains described above that contained a plasmid with a protospacer that is either an imperfect or a perfect match to sp1.8. We then introduced a plasmid with an inducible copy of *cas3*, under non-inducing conditions, to avoid interference. Following induction of *cas3* expression, we harvested cells and PCR-amplified the 5□ end of the CRISPR-II array to determine whether new spacers had been acquired because of primed adaptation. We observed robust primed adaptation for the protospacer with a perfect match to sp1.8, but no detectable adaptation for the off-target protospacer with an imperfect match to sp1.8 (Figure 4B). We conclude that off-target Cascade binding events do not lead to primed adaptation.

### Strong Cascade binding to protospacers with extensive mismatches at the crRNA PAM-distal end

To further delineate the protospacer sequence requirements for Cascade binding, interference and primed adaptation, we constructed 13 variants of a protospacer that matches sp1.8. We selected sp1.8 because it elicits robust Cascade binding, interference, and primed adaptation (Figures 3 and 4). Protospacer variants (Figure 5A) included those with (variant i; the “optimal” protospacer) full sequence complementarity, and an optimal, AAG PAM; (variants ii - iii) non-optimal PAMs: CCG, which is expected to completely abolish Cascade binding [34], and ATT, a sub-optimal sequence previously shown to cause primed adaptation but not detectable interference [31]; (variants iv - viii) two or three mismatches in the first three positions of the seed; and (variants ix – xiii) stretches of ≥6 nt mismatches at various positions within the protospacer.

We pooled cells containing each of the protospacer variants. We used ChIP of Cse1-FLAG_3_ in Δ*cas3* cells to measure association of Cascade with all protospacers within the pool (see Methods). As expected, the protospacer with a CCG PAM (variant ii) had far less Cascade association than did the optimal protospacer (variant i) (Figure 5A). We presume that the level of ChIP signal for the protospacer with the CCG PAM (variant ii) represents the background of this experiment. The protospacer with a sub-optimal, ATT PAM (iii), showed reduced Cascade binding relative to the optimal protospacer (variant i), but was well above the experimental background (Figure 5A). Similarly, mismatches in the seed region (variants iv - viii) resulted in a reduction in Cascade association (Figure 5A). Our data for PAM and seed mutants are consistent with earlier studies showing that these sequences are important for Cascade binding [17,29,30,34].

**Figure 5.**
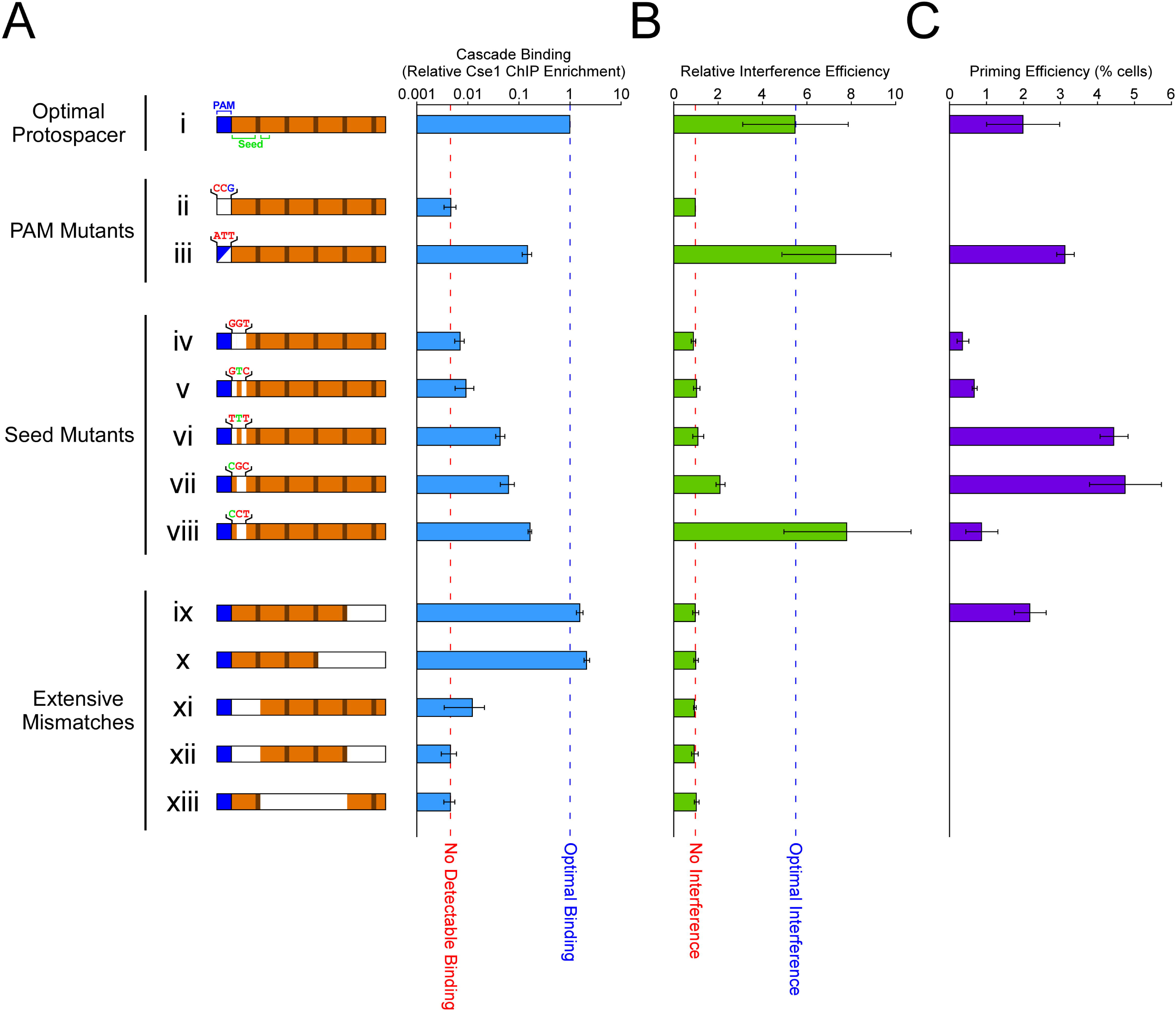
Assessment of Cascade binding, interference and primed adaptation for a panel of protospacer variants. **(A)** Relative Cse1 association (in strain LC099, which expresses CRISPR-I spacer #8 crRNA from the native CRISPR-I array) for each of 13 protospacer variants. The protospacer variants are: (i) optimal protospacer that has a perfect match to CRISPR-I spacer #8 and an AAG PAM (pLC023); (ii) CCG PAM (pLC027); (iii) ATT PAM (pLC029); (iv) mismatches at positions 1-3 (pLC031; wild-type is CTG); (v) mismatches at positions 1 and 3 (pLC033); (vi) mismatches at positions 1 and 3 (pLC035); (vii) mismatches at positions 2 and 3 (pLC032); (viii) mismatches at positions 2 and 3 (pLC034); (ix) mismatches across positions 25-32 (pLC024); (x) mismatches across positions 19-32 (pLC025); (xi) mismatches across positions 1-6 (pLC026); (xii) mismatches across positions 1-6 and 25-32 (pLC028); (xiii) mismatches across positions 7-24 (pLC030). Values represent the average of five independent replicate experiments. Error bars show one standard deviation from the mean. **(B)** Relative efficiency of interference (see Methods for details) for each of the indicated protospacer variants (pLC023-pLC035). Values represent the average of three independent replicate experiments. Error bars show one standard deviation from the mean. **(C)** Levels of primed adaptation in AMD688 cells expressing CRISPR-I spacer #8 from a native CRISPR-I array, *cas3* (pAMD191), and with each of the 13 indicated protospacer variants (pLC023-pLC035). Adaptation was measured by monitoring the conversion of a *yfp* reporter construct into the actively expressed state (see methods for details). Values shown are the percentage of the population that converted to YFP^+^. Background levels of adaptation were detected for variants (ii), (x), (xi), (xii) and (xiii).

Mismatches in the protospacer from positions 1-6 (variants xi and xii) or 7-20 (variant xiii) abolished Cascade binding (Figure 5A). This is consistent with the observation from our ChIP-seq data that sequence matches in positions 1-8 appear to be required for Cascade binding to off-target sites using sp1.8 (Figure 2B; Figure S4A). Strikingly, mismatches across positions 25-32 (variant ix) or positions 19-32 (variant x) did not reduce Cascade association relative to the optimal protospacer (variant i) (Figure 5A). Thus, our data confirm that PAM-proximal sequence is necessary for Cascade binding, while PAM-distal sequence is insufficient for Cascade binding.

### Extensive crRNA-protospacer base-pairing is required for interference and primed adaptation

We next determined which of the protospacer variants lead to interference. Using a modification of a previously described assay (see Methods) [23, 31], we measured the level of interference with a plasmid target for each of the 13 protospacers, using Δ*cas1* cells that cannot acquire new spacers; primed adaptation cannot contribute to the level of interference in these cells. As expected, the optimal protospacer (variant i) was associated with robust levels of interference, whereas protospacer variants that do not bind Cascade (variants ii, xi, xii, and xiii; Figure 5A) were not associated with detectable interference (Figure 5B). Protospacers with PAM and seed variants that showed reduced but not abolished Cascade binding (variants iii, vi, vii, and viii; Figure 5A) were associated with a range of interference levels that correlate well with the level of Cascade binding. Seed mutants with stronger defects in binding exhibited on detectable interference. However, the ability of protospacers to cause interference did not always correlate with the level of Cascade association. Specifically, we detected no interference for either of the protospacer variants with mismatches only at the PAM distal end (variants ix and x; Figure 5B), even though these protospacers bind Cascade at least as well as the optimal protospacer (Figure 5A).

Previous studies have proposed that some protospacers with sub-optimal PAMs or mismatches in the seed region are not subject to detectable interference, but do cause primed adaptation [21,23,28,31,35]. We determined whether the 13 protospacer variants caused primed adaptation in a plasmid context. We used a highly sensitive assay for adaptation that relies on expression of a *yfp* reporter gene that is encoded immediately upstream of a CRISPR array. Translation is terminated upstream of *yfp* in cells without newly acquired spacers, whereas acquisition of one spacer/repeat puts *yfp* back in frame [43], causing cells to fluoresce. We introduced an inducible copy of *cas3* into cells with an intact CRISPR-I array, and containing each of the protospacers on a high-copy plasmid, and the *yfp* reporter construct. We then induced expression of *cas3* and measured the level of primed adaptation using flow cytometry. In this experiment we expect the sp1.8 crRNA from the native CRISPR-I array to cause primed adaptation. We detected primed adaptation for all protospacers associated with detectable interference (variants i, iii, vii, and viii; Figure 5C), although the level of adaptation was lower for two of the constructs with the highest levels of interference (variants i and viii). This is likely to be due to high levels of interference reducing the available substrate for adaptation [33]. By contrast, we observed no adaptation for protospacers that do not bind Cascade (variants ii, xi, xii, and xiii; Figure 5C). Strikingly, we observed primed adaptation for four protospacers that were not associated with detectable interference (Figure 5C). Three of these protospacers have seed mismatches, and exhibited the lowest levels of Cascade binding (Figure 5A; variants iv, v and vi). The other protospacer has mismatches across positions 25-32 (variant ix). Thus, for these protospacers, we detected robust Cascade binding and primed adaptation but we were unable to detect interference. For the protospacer with mismatches across positions 19-32 (variant x), we detected no primed adaptation. Thus, for this protospacer, we detected robust Cascade binding, but no primed adaptation or interference. Overall, our data suggest that extensive crRNA-protospacer base pairing from the PAM-proximal end is required for both interference and primed adaptation, and that primed adaptation is a more sensitive assay of CRISPR-Cas function than interference.

## DISCUSSION

### Base-pairing in the seed region together with an AAG PAM is sufficient for Cascade to bind DNA

No previous studies have measured Cascade binding to protospacer DNA *in vivo*. Our ChIP data indicate that an AAG PAM and as little as 5 nucleotides of base-pairing at the start of the seed region are sufficient for *E. coli* Cascade to bind DNA targets. The sequence requirements for protospacer binding in type II systems are similarly relaxed [44–46]. The affinity of Cascade for a protospacer increases as the extent of base-pairing increases, but maximal affinity occurs with no more than an 18 bp match at the PAM-proximal end (Figure 5A). Analysis of Cascade interactions with DNA *in vitro* suggests that Cascade associates for brief periods with PAM-containing sequences, and for longer periods if there is partial base-pairing in the seed region [28]. Our data support these observations, although we did not detect ChIP signal at PAM sequences that lack seed matches, suggesting that seed base-pairing contributes more to Cascade association *in vivo*. Consistent with this suggestion, the difference in ChIP signal for off-target and on-target sites is considerably less than the difference between dwell times *in vitro* [28], although the use of crosslinking in ChIP may also contribute to this difference since crosslinking “locks” Cascade on the DNA.

### AAG is the optimal PAM in *E. coli*

Three previous studies proposed that AAG, GAG, TAG, AGG, and ATG are optimal PAMs in *E. coli* [23,31,47], while another study suggested that AAG, ATG and GAG PAMs were associated with moderately higher affinity Cascade binding than an AGG PAM [34]. Our data clearly indicate that AAG is the optimal PAM for off-target sites, with most off-target Cascade binding events being associated with an AAG PAM. Specifically, 65% of Cascade binding sites associated with a detectable motif have an AAG PAM for the crRNAs targeting *lacZ* and *araB*, and the plasmid-encoded sp1.8 crRNA. Moreover, off-target Cascade binding events with higher enrichment scores, suggestive of higher Cascade affinity, were more likely to be associated with an AAG PAM than Cascade binding events with lower enrichment scores (76% vs 61% for the top 20% and bottom 80% of bound regions, respectively, after sorting by Cse1 enrichment level). We hypothesize that the dependence on the PAM for Cascade binding is increased in situations where base-pairing only occurs in the seed region. According to this model, complete or near-complete base-pairing between the crRNA and protospacer would weaken the requirement for an optimal PAM, obscuring differences in PAM affinity. This would explain why previous studies suggested that there are at least three optimal PAMs [23,31,34,47].

### Defining the crRNA seed

The seed region of a crRNA has been previously defined as positions 1-5 and 7-8, with position 1 being immediately adjacent to the PAM [29]. However, our data suggest that the length of the seed varies between crRNAs, since we observed off-target binding with some crRNAs that requires base-pairing in positions 1-5, whereas off-target binding for other crRNAs requires base-pairing up to position 9 (Figures 1 and 2; Figure S3 and S4). We propose that the crRNA sequence determines the length of the seed, and that this reflects the initial binding mode, prior to extended base-pair formation. Every 6^th^ position of the crRNA is flipped out in the Cascade-protospacer complex, and hence does not contribute to base-pairing [15,48,49]. Consistent with this, the importance of position 6 for off-target binding is substantially less than that of positions 1-5 (Figures 1 and 2; Figure S3 and S4). Nonetheless, off-target protospacers had a sequence match to the crRNA at position 6 far more frequently than expected by chance (45% for the crRNAs targeting *lacZ* and *araB*, and the plasmid-encoded sp1.8 crRNA; Binomial Test *p*-value = 2.4e^-10^). We hypothesize that the initial binding of Cascade to a protospacer includes base-pairing interactions at position 6, but that the complex rapidly transitions to a conformation in which the 6^th^ position is flipped out of the helix. Our data are consistent with an *in vitro* study of another type I-E system, where position 6 was also shown to contribute to off-target Cascade binding [50]. The apparent requirement for a sequence match at position 6 is not consistent across all crRNAs we tested, suggesting that the pathway towards stable seed base-pairing differs in a sequence-dependent manner.

### Interference and primed adaptation require extended R-loop formation

Although binding of Cascade to a DNA target requires relatively little sequence identity, our data indicate that robust interference and primed adaptation require at least 18-25 bp, beginning in the seed region. This is consistent with *in vitro* data showing that near-complete R-loop formation is required to license Cas3 activity [16]. Thus, although Cascade binds DNA promiscuously, functional binding occurs with high specificity. Our data support a previously proposed model in which extended R-loop formation triggers a conformational change in Cascade at the PAM-distal end of the spacer, which is then transmitted, presumably through Cse2, to PAM-associated Cse1 [16, 51]. This change in Cse1 conformation then recruits Cas3, and/or activates the nuclease activity of Cas3, as suggested by a recent structural study [51].

### Evidence that interference and primed adaptation are obligately coupled processes

Primed adaptation was initially proposed to be an alternative pathway to interference, with optimal PAM/seed sequences leading to interference, and sub-optimal sequences leading to primed adaptation [21,23,28,31,35,52]. However, primed adaptation has been observed in situations where interference occurs (Figure 5, variants i, iii, vii, and viii) [22,24,32,33], suggesting that primed adaptation and interference can be coupled processes, and supporting the idea that primed adaptation is a positive feedback loop [22]. While these data show that primed adaptation and interference can occur at the same time at a population level, they do not necessarily indicate that individual primed adaptation and interference events are coupled. Moreover, while it has been proposed that interference and primed adaptation are obligately coupled [53], this has not been tested. There are many examples where primed adaptation has been observed in the absence of detectable interference [21,23,31,32,35,52]. However, this can be explained by the fact that primed adaptation is likely to be a more sensitive assay of CRISPR-Cas function than interference, as there would be detectable primed adaptation but not detectable interference in cells where target DNA replication outpaces interference [53]. Our data are consistent with a model in which primed adaptation and interference are coupled processes: seed mismatches reduce Cascade binding, and we observe a corresponding effect on interference and primed adaptation, with primed adaptation being a more sensitive assay for CRISPR-Cas function (Figure 5). The only exception to this trend is the seed mismatch with the highest level of binding (Figure 5, variant viii), which has relatively low levels of primed adaptation. However, very efficient interference with this variant likely depletes substrate for primed adaptation [33]. Unexpectedly, we observed primed adaptation in the absence of detectable interference for a protospacer with mismatches across positions 25-32 (Figure 5, variant ix). We propose that this degree of mismatch at the 3L end of the crRNA greatly reduces, but does not abolish, the isomerization of Cascade into the “active” state that recruits/activates Cas3.

### Extensive, inert, off-target binding of Cascade

Cascade has many off-target binding sites due to its ability to bind DNA with low sequence-specificity. Consequently, the endogenous crRNAs transcribed from the bacterial genome result in extensive off-target binding, even in the absence of an on-target site. Since off-target binding does not involve extended R-loop formation, it has no deleterious effects on genome integrity. We also observed no impact on transcription associated with any of the off-target binding events, despite that fact that targeted Cascade binding is known to repress transcription by occluding promoters or acting as a roadblock for elongating RNA polymerase [38, 40]. Transcription repression by Cascade is considerably weaker when targeting within a transcribed region (i.e. acting as a roadblock) [38]. Given that the location of off-target Cascade binding sites is essentially random with respect to genome organization, and that genes make up ∼90% of the *E. coli* genome, off-target Cascade binding is expected to be primarily intragenic. This may partly explain the lack of impact on transcription. Moreover, a recent study showed that the level of repression by Cascade occlusion of a promoter is greatly reduced with as few as 6 bases mismatched at the PAM-distal end of the spacer/protospacer [54], suggesting that even intergenic off-target Cascade binding sites would be transcriptionally inert. We propose that incomplete R-loop formation results in an unstable Cascade-DNA complex with a relatively high rate of dissociation, such that it cannot compete effectively with initiating or elongating RNA polymerase. Consistent with this model, stable association of Cascade with DNA *in vitro* has been shown to require near-complete R-loop formation [18]. We conclude that type I CRISPR-Cas systems have evolved to tolerate off-target binding driven by the endogenous crRNAs, and are only functional at on-target sites. Given the length of crRNA spacers in type I systems, there is no expectation of complete or near-complete spacer-protospacer base-pairing by chance. It is important to note that self-targeting by type I CRISPR-Cas systems has been described previously, but these would be considered “on-target” events, likely caused by acquisition of spacers from the chromosome. As expected for spacers with perfect sequence complementarity, these self-targeting crRNAs are typically functional in gene regulation and interference [36,37,55].

### Not all crRNAs are created equal

The *E. coli* genome encodes at least 19 crRNAs, yet our data suggest that only four crRNAs contribute to off-target binding of Cascade. All four of these crRNAs are encoded in the CRISPR-I array, and the majority of off-target binding is driven by just one, sp1.8. The lack of off-target binding driven by CRISPR-II crRNAs is likely due to weak transcription of this array, which is repressed by H-NS [56]. In contrast, the CRISPR-I array is likely co-transcribed with the upstream *cas* genes, which are strongly transcribed in the strain used in this study. The preference for specific spacers within CRISPR-I cannot be explained by differences in expression levels, since the crRNAs are transcribed as a single RNA. Rather, biases in spacer usage are more likely due to differential assembly of specific crRNAs into Cascade. Consistent with this, a previous study surveyed crRNAs associated with Cascade. Spacers #2, #4 and #8 from CRISPR-I represented 68% of the Cascade-associated crRNAs [7]. The cause of this bias is unclear, but may in part be due to differences in RNA secondary structure between spacers, which could impact the efficiency of RNA processing by Cas6e. Consistent with this, RNA secondary structure of repeat sequences, and associated processing by Cas6, has been shown to be impacted by spacer sequences in the type I-D system of *Synechocystis* sp. PCC 6803 [57]. Nonetheless, it is likely that other factors influence the level of off-target binding, since the relative association of crRNAs for spacers #2, #4 and #8 with Cascade is likely to be similar [7], and sp1.2 causes efficient interference (Figure 4A), but sp1.8 drives a disproportionately high level of off-target binding relative to sp1.2. Strikingly, there are many more chromosomal sequence matches to the seed sequence of sp1.8 coupled with an AAG PAM than for any other spacer (Table S4). This is likely due to the fact that the sequence from position −1 (i.e. last base of the PAM) to +8 of sp1.8 differs from the canonical Chi site sequence(5□-GCTGGTGG-3□)[58] by a single nucleotide; Chi sites are strongly enriched in the *E. coli* K-12 genome [59]. Moreover, positions 3-7 of sp1.8 (5□-GCTGG-3□) are a perfect match to a sequence that is strongly enriched in the *E. coli* K-12 genome [59]. We conclude that extensive off-target binding driven by sp1.8 is likely due to a combination of high association with Cascade, and a relatively high abundance of potential binding sites in the genome.

## METHODS

### Strains and plasmids

All strains, plasmids, oligonucleotides and purchased, chemically synthesized dsDNA fragments are listed in Table S5. All strains are derivatives of MG1655 [59]. CB386 has been previously described [38]. CB386 contains a chloramphenicol resistance cassette in place of *cas3*. We removed this cassette using Flp recombinase, expressed from plasmid pCP20 [60], to generate strain AMD536. Epitope-tagged strains AMD543 and AMD554 (Cse1-FLAG_3_ and FLAG_3_-Cas5, respectively) are derivatives of CB386, and were generated using the previously described FRUIT method of recombineering [61]. Cse1 was C-terminally tagged in AMD543 by inserting a FLAG_3_ tag immediately upstream of codon 495 using oligonucleotides JW6364 and JW6365. Tagging of Cse1 resulted in an 8 amino acid C-terminal truncation. We predicted based on phylogenetic comparisons and on structural data [49] that this truncation would not impact the function of Cse1. Cas5 was N-terminally tagged in AMD554 by inserting FLAG_3_ using oligonucleotides JW6272 and JW6273. LC060 is a derivative of AMD536 and was generated using (i) FRUIT [61] with oligonucleotides JW7537-JW7540 to delete the CRISPR-II locus, (ii) P1 transduction of the CB386 (Δ*cas3* P*cse1*)::(cat::P_J23199_) region, (iii) FRUIT [61] to C-terminally tag Cse1 with FLAG_3_ (as described above for AMD543), and (iv) pCP20-expressed Flp recombinase [60] to remove the *cat* cassette. LC074 is a derivative of AMD536 in which the CRISPR-I array was deleted using FRUIT [61] with oligonucleotides JW7529 and JW7530 and a synthesized dsDNA fragment (gBlock 14148263; Integrated DNA technologies). LC077 is a derivative of LC074 in which Cse1 was C-terminally tagged with FLAG_3_ (as described above for AMD543). AMD566 is a derivative of AMD536 in which Cse1 was C-terminally tagged with FLAG_3_ (as described above for AMD543). LC099 is a derivative of AMD566 in which the off-target binding site for Cascade in *yggX* was mutated using FRUIT [61] with oligonucleotides JW7635-8. LC103 is a derivative of AMD536 in which the the *yggX* gene was replaced with a kanamycin resistance cassette using P1 transduction from the Keio Collection Δ*yggX*::*kan*^R^ strain [62]. LC106 is a derivative of LC103 with an unmarked, scar-free deletion of *cas1* made using FRUIT with oligonucleotides JW7898-JW7901. AMD688 is a strain that contains a previously reported *yfp* reporter construct that can be used to measure adaptation levels [43]. AMD688 was constructed by P1-transduction of the Δ*cas3*::*cat* cassette from CB386 into MLS1003 (provided by the Lundgren laboratory). The *cat* gene was removed using Flp recombinase, expressed from plasmid pCP20 [60]. AMD688 has an intact copy of the CRISPR-I array (co-transduced with the Δ*cas3*::*cat* cassette from CB386) but lacks the CRISPR-II array.

Plasmids that express crRNAs targeting the *lacZ* promoter (pCB380) and *araB* promoters (pCB381) have been described previously [38]. All other crRNA-expressing plasmids are derivatives of pAMD179. pAMD179 was constructed by amplifying a DNA fragment from plasmid pAMD172 (Integrated DNA Technologies) using with oligonucleotides JW6421 and JW6513. This DNA fragment was cloned into pBAD24 [63] cut with *Nhe*I and *Hin*dIII (NEB) using the In-Fusion method (Clontech). The inserted fragment contains two repeats from the CRISPR-I array, separated by a stuffer fragment containing *Xho*I and *Sac*II restriction sites, and an intrinsic transcription terminator downstream of the second repeat. To clone individual spacers, pairs of oligonucleotides were annealed, extended, and inserted using In-Fusion (Clontech) into the *Xho*I and *Sac*II sites of pAMD179 to generate pLC008 (with oligonucleotides JW6518 and JW7911), pLC010 (with oligonucleotides JW6518 and JW7912), and pAMD189 (with oligonucleotides JW7598 and JW7693). Note that the derivatives of sp1.8 expressed from pLC008 and pLC010 differ from sp1.8 at the last two nucleotide positions to facilitate cloning. These mismatches are not expected to affect crRNA function [23, 38].

pLC021, pLC022, and pLC057 are derivatives of pBAD24 [63] that contain a protospacer matching the off-target Cascade binding site in *yggX* (pLC021), a protospacer with a perfect match to sp1.8 (pLC022), or a protospacer with a perfect match to sp1.2 (pLC057). These plasmids were constructed by annealing and extending pairs of oligonucleotides (JW7913 and JW7914 for pLC021, JW7924 and JW7925 for pLC022, and JW9131 and JW9132 for pLC057), and cloning the resultant DNA fragments into the *Eco*RV and *Sph*I sites of pBAD24. pAMD191 is a derivative of pBAD33 [63] that expresses *cas3* under arabinose control. To construct pAMD191, *cas3* was amplified by colony PCR using oligonucleotides JW7736 and JW7738. The PCR product was cloned into the *Sac*I and *Hin*dIII sites of pBAD33 using In-Fusion (Clontech). All protospacers described in Figure 5 are cloned into plasmid pLC020, the “pre-protospacer plasmid”, which is a derivative of pBAD24 [63]. pLC020 was generated by cloning the ∼500 bp region upstream of *E. coli thyA* (amplified by colony PCR using oligonucleotides JW8040 and JW8128) and the ∼500 bp region downstream of *E. coli thyA* (amplified by colony PCR using oligonucleotides JW8042 and JW8043) into the *Eco*RI site of pBAD24 using In-Fusion (Clontech), simultaneously generating a new *Eco*RI site between the upstream and downstream regions of *thyA*. The *thyA* gene was then amplified by colony PCR using a universal forward primer (oligonucleotide JW8129) and each of 13 reverse primers (oligonucleotides JW8130, JW8139, JW8145, JW8169, JW8499-JW8502, and JW8675-JW8679) containing the 13 protospacer variants described in Figure 5. The resulting PCR products were cloned into the *Eco*RI site of the pBAD24 derivative using In-Fusion (Clontech) to generate plasmids pLC023-pLC035 (see Table S5 for details). Note that plasmids pLC024 and pLC025 differ from pLC023 and pLC026-pLC035 at the nucleotide position immediately adjacent to the protospacer, on the PAM-distal side. Differences at this nucleotide position are not expected to affect Cascade binding, interference or primed adaptation.

### ChIP-qPCR

Cells were grown overnight in LB, subcultured in LB supplemented with 0.2% arabinose and 100 μg/mL ampicillin at 37 °C with aeration to an OD_600_ of ∼0.6. AMD566 and LC099 with either pLC008 or pLC010 were used for ChIP-qPCR. ChIP-qPCR was performed as described previously [64], except that 2 μL anti-FLAG M2 monoclonal antibody (Sigma) and 1 μL anti-σ^54^ monoclonal antibody (NeoClone) were included simultaneously in the immunoprecipitation step. qPCR was performed using oligonucleotides JW7490-1 (amplifies the off-target site in *yggX*) and JW7922-3 (amplifies the region upstream of *hypA*). Since σ^54^ is known not to bind within *yggX* [65], we were able to normalize binding of Cse1 within *yggX* to the binding of σ^54^ upstream of *hypA*.

### ChIP-seq

Strains AMD543, LC060, LC077, AMD543/AMD554 with pCB380/pCB381, and strain LC077 with pLC008, were used for ChIP-seq of Cse1-FLAG_3_ and FLAG_3_-Cas5, except that ampicillin was only included for experiments involving a crRNA-expressing plasmid, and arabinose was only included for experiments using pLC008. Cells were grown and processed as described for ChIP-qPCR. ChIP-seq was performed in duplicate, following a previously described protocol [66] using 2 μL anti-FLAG M2 monoclonal antibody (Sigma). Sequencing was performed on an Illumina High-Seq 2000 Instrument (Next-Generation Sequencing and Expression Analysis Core, State University of New York at Buffalo) or an Illumina Next-Seq Instrument (Wadsworth Center Applied Genomic Technologies Core). ChIP-seq data analysis was performed as previously described [67], with reads mapped to the updated MG1655 *E. coli* genome (accession code U00096.3). Relative sequence coverage values were calculated by dividing the sequence read coverage at a given genomic location by (total number of sequence reads in the run/100,000). Values plotted in Figures 1A-B, 2A and 2D are the maximum values in 1 kbp regions across the genome. R^2^ values comparing ChIP-seq datasets were calculated by comparing read coverage at peak centers for all peaks identified for the analyzed datasets. Read coverage at peak centers was determined using a custom Python script. Sequence motifs were identified using MEME (version 4.12.0) [68] with default parameters.

### RNA-seq

RNA-seq was performed in duplicate with strains AMD536 and LC074, with and without pAMD191. Cells were grown overnight in LB, subcultured in LB (supplemented with 0.2% arabinose and 100 μg/mL ampicillin for experiments involving pAMD191) at 37 °C with aeration to an OD_600_ of ∼0.6. RNA was purified using a modified hot phenol method, as previously described [69]. Purified RNA was treated with 2 μL DNase (TURBO DNA-free kit; Life Technologies) for 45 minutes at 37 °C, followed by phenol extraction and ethanol precipitation. The RiboZero kit (Epicentre) was used to remove rRNA, and strand-specific cDNA libraries were created using the ScriptSeq 2.0 kit (Epicentre). Sequencing was performed using an Illumina Next-Seq Instrument (Wadsworth Center Applied Genomic Technologies Core). Differential RNA expression analysis was performed using Rockhopper (version 2.03) with default parameters [70]. Differences in RNA levels were considered statistically significant for genes with *q*-values ≤ 0.01.

### Plasmid transformation efficiency assay

LC103 was transformed with either empty pBAD33 or pAMD191 (expresses *cas3*) and these strains were then transformed with either pBAD24 (no protospacer), pLC021 (protospacer with a perfect match to sp1.8), pLC022 (protospacer with an imperfect match to sp1.8, corresponding to the off-target site in *yggX*) or pLC057 (protospacer with a perfect match to sp1.2). Cells were plated on M9 medium supplemented with 0.2% glycerol, 0.2% arabinose, 100 μg/mL ampicillin and 30 μg/mL chloramphenicol at 37 °C. After overnight growth, colonies were counted, and the relative transformation efficiency was calculated as the ratio of transformants for pAMD191-containing cells to transformants for pBAD33-containing cells for each transformed protospacer-containing plasmid.

### PCR to assess primed adaptation

Primed adaptation was assessed for AMD536 with pAMD191 and either pLC021 or pLC022 (Figure 4B), and MG1655/AMD536/AMD543/AMD544 with pAMD191 and pAMD189 (expresses a self-targeting crRNA; Figure S1). Cells were grown overnight in LB supplemented with 100 μg/mL ampicillin and 30 μg/mL chloramphenicol at 37°C with aeration, and sub-cultured the next day in LB supplemented with chloramphenicol and 0.2% arabinose at 37°C with aeration for six hours. Cells were pelleted from 1 mL of culture by centrifugation, and cell pellets were frozen at −20°C. PCRs were then performed on the cell pellets, amplifying the CRISPR-II array using oligonucleotides JW7818 and JW7819. PCR products were visualized on acrylamide gels.

### Sequence analysis of protospacers from a pooled ChIP library

LC099 with each of the 13 protospacer variant plasmids (pLC23-pLC035), was grown overnight in LB supplemented with 100 μg/mL ampicillin. 10 mL subcultures were grown in LB supplemented with 100 μg/mL ampicillin and 0.2% arabinose at 37°C with aeration to an OD_600_ of ∼0.6. 3 mL from each culture were combined. ChIP was performed on mixed cultures using 2 μL M2 anti-FLAG monoclonal antibody (Sigma), as previously described [64]. A Zymo “PCR Clean and Concentrate” kit was used to purify ChIP and input DNA. A 50 μL FailSafe (Epicentre) PCR reaction using FailSafe PCR 2X PreMix “C” and 5.48 ng of ChIP DNA was performed following the manufacturer’s instructions, using oligonucleotide JW8567 and each of oligonucleotides JW8537, JW8556, JW8557, JW8558, JW8559, JW8561, JW8562, JW8563, JW8564, and JW8565 (these incorporate different Illumina indices). PCR products were purified and concentrated using 0.8X Ampure Beads (Beckman Coulter Life Sciences) and sequenced on an Illumina Mi-Seq Instrument (Wadsworth Center Applied Genomic Technologies Core). Sequence reads were mapped to each of the 13 protospacer variants using a custom Python script. Relative protospacer abundance in input and ChIP samples for each protospacer were normalized to the total sequence reads. Values for normalized protospacer abundance were further normalized to values from the input sample. Protospacer abundance values are reported relative to those for the optimal protospacer (variant i in Figure 5).

### Measuring interference for a pooled protospacer library

Overnight cultures of LC106 strains with each of the 13 protospacer plasmids (pLC023-pLC035) were grown in LB with 100 μg/mL ampicillin and 30 μg/mL kanamycin. All 13 cultures were combined to make a single subculture (7.7 μL of each overnight culture into a single 10 mL culture). Electrocompetent cells were made and transformed with either empty pBAD33 or pAMD191 (pBAD33-*cas3*). Transformants were plated onto M9 agar supplemented with 0.2% glycerol, 0.2% arabinose, and 30 μg/mL chloramphenicol, and grown overnight at 37°C. Cells were scraped off plates, washed in LB, and protospacers were PCR amplified from cell pellets with oligonucleotide JW8567 and each of oligonucleotides JW8537, JW8558, JW8559, JW8562, JW8563, and JW8566 (these incorporate different Illumina indices). PCR products were purified and concentrated with 0.8X AMPure Beads (Beckman Coulter Life Sciences), and sequenced using a Illumina MiSeq Instrument (Wadsworth Center Applied Genomic Technologies Core). Sequence reads were mapped to each of the 13 protospacer variants using a custom Python script. “Relative Interference Efficiency” was calculated for each protospacer variant by dividing the number of sequence reads from cells transformed with empty pBAD33 by the number of sequence reads from cells transformed with pAMD191 (pBAD33-*cas3*), and normalizing to the value for the protospacer with a CCG PAM (variant ii in Figure 5).

### Measuring primed adaptation using a YFP fluorescent reporter

MLS1003 was transformed with each of plasmids LC023-LC035, and each of these strains was transformed with the *cas3*-expressing plasmid pAMD191. Cells were grown overnight at 37 °C with shaking in LB supplemented with 100 μg/ml ampicillin and 30 μg/ml chloramphenicol. Cells were subcultured 1:100 for six hours in LB supplemented with 0.2% L-arabinose and 20 μg/ml chloramphenicol at 37 °C with shaking. Cells were pelleted by centrifugation and resuspended in M9 minimal medium in twice the original volume (OD_600_ values of ∼1). Cells were transferred to 5 ml polystyrene round-bottom tubes and were analyzed by flow cytometry for single-cell detection of *yfp* expression using the BD FACSAria iiu cell sorter. 100,000 events were recorded for each sample. Experiments were performed for between three and ten independent biological replicates.

## ACKNOWLEDGEMENTS

We thank Chase Beisel for sharing strains and plasmids. We thank Lina Amlinger and Magnus Lundgren for the YFP-expressing adaptation reporter strain. We thank the Wadsworth Center Applied Genomic Technologies Core Facility and the University at Buffalo Genomics and Bioinformatics Core Facility for Sanger and MiSeq sequencing. We thank the Wadsworth Center Media and Tissue Culture and Glassware Core Facilities. We thank Todd Gray, Keith Derbyshire, and Shailab Shrestha for helpful discussions. This study was supported by NIH Grant AI126416 (to J.T.W.) and a University at Albany, SUNY, RNA Fellowship (to L.A.C.).

## SUPPORTING INFORMATION

**Figure S1. FLAG_3_-tagged Cse1 and Cas5 are fully functional for primed adaptation.** PCR-amplification of the start of each of the CRISPR-I and CRISPR-II arrays to detect primed adaptation in cells expressing *cas3* (pAMD191) and a plasmid-encoded crRNA that perfectly targets a sequence on the same plasmid (pAMD189). Adaptation was assessed for (i) MG1655 (no *cas* gene expression, except for plasmid-encoded *cas3* (pAMD191)), (ii) MG1655 with constitutive expression of *cas* genes (AMD536), (iii) MG1655 *cse1*-FLAG_3_ with constitutive expression of *cas* genes (AMD543), and (iv) MG1655 FLAG_3_-Cas5 with constitutive expression of *cas* genes (AMD554).

**Figure S2. crRNA spacers used in this study. (A)** Sequence of the crRNA spacer targeting the *lacZ* promoter (pCB380). **(B)** Sequence of the crRNA spacer targeting the *araB* promoter (pCD381). **(C)** Sequence of the CRISPR-I array. Spacers #1, #3, #4 and #8 are underlined. **(D)** Sequence of the CRISPR-II array. Spacer #2 is underlined. **(E)** Sequence of a portion of the CRISPR-I spacer #8 crRNA-expressing plasmid (pLC008). Note that the sequence downstream of the second repeat (underlined) can be used as a spacer.

**Figure S3. Spacer #2 of CRISPR-II directs Cascade binding in cells lacking CRISPR-I.** Enriched sequence motif associated with Cascade binding sites in cells expressing only endogenous crRNAs, where CRISPR-I is deleted (LC077). The motif is associated with CRISPR-II spacer #2, as indicated. The likely PAM sequence is also indicated. The number of identified motifs and the MEME E-value are shown.

**Figure S4. Sequence motifs associated with Cascade binding in cells expressing CRISPR-I spacer #8 from a plasmid. (A)** Sequence of the most strongly enriched motif, as identified by MEME, in ΔCRISPR-I cells (LC077) expressing CRISPR-I spacer #8 from a plasmid (pLC008). The motif is associated with CRISPR-I spacer #8, as indicated. The likely PAM sequence is also indicated. The number of identified motifs and the MEME E-value are shown. **(B)** The second enriched sequence motif, as identified by MEME, in ΔCRISPR-I cells (LC077) expressing1033 CRISPR-I spacer #8 from a plasmid (pLC008). The motif is associated with the sequence immediately downstream of the second repeat on the crRNA plasmid, as indicated.

**Table S1. Lists of ChIP-seq peak coordinates.**

**Table S2. Lists of regions used to search for enriched sequence motifs.**

**Table S3. Analysis of RNA-seq data.**

**Table S4. Numbers of potential off-target chromosomal binding sites for spacers in the CRISPR-I array.**

**Table S5. Strains, Plasmids, Oligonucleotides, and Chemically Synthesized dsDNA fragments used in this study.**

## REFERENCES

1. Grissa I, Vergnaud G, Pourcel C. The CRISPRdb database and tools to display CRISPRs and to generate dictionaries of spacers and repeats. BMC Bioinformatics. 2007;8: 1–10. doi:10.1186/1471-2105-8-172

2. Bolotin A, Quinquis B, Sorokin A, Ehrlich SD. Clustered regularly interspaced short palindrome repeats (CRISPRs) have spacers of extrachromosomal origin. Microbiol Read Engl. 2005;151: 2551–2561. doi:10.1099/mic.0.28048-0

3. Makarova KS, Grishin NV, Shabalina SA, Wolf YI, Koonin EV. A putative RNA-interference-based immune system in prokaryotes: computational analysis of the predicted enzymatic machinery, functional analogies with eukaryotic RNAi, and hypothetical mechanisms of action. Biol Direct. 2006;1: 7. doi:10.1186/1745-6150-1-7

4. Mojica FJM, Díez-Villaseñor C, García-Martínez J, Soria E. Intervening sequences of regularly spaced prokaryotic repeats derive from foreign genetic elements. J Mol Evol. 2005;60: 174–182. doi:10.1007/s00239-004-0046-3

5. Pourcel C, Salvignol G, Vergnaud G. CRISPR elements in Yersinia pestis acquire new repeats by preferential uptake of bacteriophage DNA, and provide additional tools for evolutionary studies. Microbiology. 2005;151: 653–663. doi:10.1099/mic.0.27437-0

6. Nuñez JK, Kranzusch PJ, Noeske J, Wright AV, Davies CW, Doudna JA. Cas1–Cas2 complex formation mediates spacer acquisition during CRISPR–Cas adaptive immunity. Nat Struct Mol Biol. 2014;21: 528–534. doi:10.1038/nsmb.2820

7. Brouns SJJ, Jore MM, Lundgren M, Westra ER, Slijkhuis RJH, Snijders APL, et al. Small CRISPR RNAs guide antiviral defense in prokaryotes. Science. 2008;321: 960–964. doi:10.1126/science.1159689

8. Carte J, Wang R, Li H, Terns RM, Terns MP. Cas6 is an endoribonuclease that generates guide RNAs for invader defense in prokaryotes. Genes Dev. 2008;22: 3489–3496. doi:10.1101/gad.1742908

9. Jore MM, Lundgren M, van Duijn E, Bultema JB, Westra ER, Waghmare SP, et al. Structural basis for CRISPR RNA-guided DNA recognition by Cascade. Nat Struct Mol Biol. 2011;18: 529–536. doi:10.1038/nsmb.2019

10. Mojica FJM, Díez-Villaseñor C, García-Martínez J, Almendros C. Short motif sequences determine the targets of the prokaryotic CRISPR defence system. Microbiol Read Engl. 2009;155: 733–740. doi:10.1099/mic.0.023960-0

11. Sashital DG, Wiedenheft B, Doudna JA. Mechanism of foreign DNA selection in a bacterial adaptive immune system. Mol Cell. 2012;46: 606–615. doi:10.1016/j.molcel.2012.03.020

12. Hayes RP, Xiao Y, Ding F, van Erp PB, Rajashankar K, Bailey S, et al. Structural basis for promiscuous PAM recognition in type I-E Cascade from E. coli. Nature. 2016;530: 499–503.

13. Hochstrasser ML, Taylor DW, Bhat P, Guegler CK, Sternberg SH, Nogales E, et al. CasA mediates Cas3-catalyzed target degradation during CRISPR RNA-guided interference. Proc Natl Acad Sci U S A. 2014;111: 6618–6623. doi:10.1073/pnas.1405079111

14. Mulepati S, Bailey S. In vitro reconstitution of an Escherichia coli RNA-guided immune system reveals unidirectional, ATP-dependent degradation of DNA target. J Biol Chem. 2013;288: 22184–22192. doi:10.1074/jbc.M113.472233

15. Mulepati S, Héroux A, Bailey S, 19;345(6203):1479-84. S 2014 S. Crystal structure of a CRISPR RNA-guided surveillance complex bound to a ssDNA target. Science. 2014;345: 1479–1484.

16. Rutkauskas M, Sinkunas T, Songailiene I, Tikhomirova MS, Siksnys V, Seidel R. Directional R-Loop Formation by the CRISPR-Cas Surveillance Complex Cascade Provides Efficient Off-Target Site Rejection. Cell Rep. 2015; doi:10.1016/j.celrep.2015.01.067

17. Westra ER, van Erp PB, Künne T, Wong SP, Staals RH, Seegers CL, et al. CRISPR immunity relies on the consecutive binding and degradation of negatively supercoiled invader DNA by Cascade and Cas3. Mol Cell. 2012;46: 595–605.

18. Szczelkun MD, Tikhomirova MS, Sinkunas T, Gasiunas G, Karvelis T, Pschera P, et al. Direct observation of R-loop formation by single RNA-guided Cas9 and Cascade effector complexes. Proc Natl Acad Sci U S A. 2014;111: 9798–9803. doi:10.1073/pnas.1402597111

19. Sinkunas T, Gasiunas G, Waghmare SP, Dickman MJ, Barrangou R, Horvath P, et al. In vitro reconstitution of Cascade-mediated CRISPR immunity in Streptococcus thermophilus. EMBO J. 2013;32: 385–394. doi:10.1038/emboj.2012.352

20. Yosef I, Goren MG, Qimron U. Proteins and DNA elements essential for the CRISPR adaptation process in Escherichia coli. Nucleic Acids Res. 2012;40: 5569–5576. doi:10.1093/nar/gks216

21. Datsenko KA, Pougach K, Tikhonov A, Wanner BL, Severinov K, Semenova E. Molecular memory of prior infections activates the CRISPR/Cas adaptive bacterial immunity system. Nat Commun. 2012;3: 945. doi:10.1038/ncomms1937

22. Swarts DC, Mosterd C, van Passel MWJ, Brouns SJJ. CRISPR Interference Directs Strand Specific Spacer Acquisition. PLoS ONE. 2012;7: e35888. doi:10.1371/journal.pone.0035888

23. Fineran PC, Gerritzen MJH, Suárez-Diez M, Künne T, Boekhorst J, Hijum SAFT van, et al. Degenerate target sites mediate rapid primed CRISPR adaptation. Proc Natl Acad Sci. 2014;111: E1629–E1638. doi:10.1073/pnas.1400071111

24. Staals RHJ, Jackson SA, Biswas A, Brouns SJJ, Brown CM, Fineran PC. Interference-driven spacer acquisition is dominant over naive and primed adaptation in a native CRISPR–Cas system. Nat Commun. 2016;7: 12853. doi:10.1038/ncomms12853

25. Savitskaya E, Semenova E, Dedkov V, Metlitskaya A, Severinov K. High-throughput analysis of type I-E CRISPR/Cas spacer acquisition in E. coli. RNA Biol. 2013;10: 716–725. doi:10.4161/rna.24325

26. Li M, Wang R, Zhao D, Xiang H. Adaptation of the Haloarcula hispanica CRISPR-Cas system to a purified virus strictly requires a priming process. Nucleic Acids Res. 2014;42: 2483–2492. doi:10.1093/nar/gkt1154

27. Richter C, Dy RL, McKenzie RE, Watson BNJ, Taylor C, Chang JT, et al. Priming in the Type I-F CRISPR-Cas system triggers strand-independent spacer acquisition, bi-directionally from the primed protospacer. Nucleic Acids Res. 2014;42: 8516–8526. doi:10.1093/nar/gku527

28. Redding S, Sternberg SH, Marshall M, Gibb B, Bhat P, Guegler CK, et al. Surveillance and Processing of Foreign DNA by the Escherichia coli CRISPR-Cas System. Cell. 2015;163: 854–865. doi:10.1016/j.cell.2015.10.003

29. Semenova E, Jore MM, Datsenko KA, Semenova A, Westra ER, Wanner B, et al. Interference by clustered regularly interspaced short palindromic repeat (CRISPR) RNA is governed by a seed sequence. Proc Natl Acad Sci USA. 2011;108: 10098–10103.

30. Wiedenheft B, van Duijn E, Bultema JB, Bultema J, Waghmare SP, Waghmare S, et al. RNA-guided complex from a bacterial immune system enhances target recognition through seed sequence interactions. Proc Natl Acad Sci U S A. 2011;108: 10092–10097. doi:10.1073/pnas.1102716108

31. Xue C, Seetharam AS, Musharova O, Severinov K, Brouns SJ, Severin AJ, et al. CRISPR interference and priming varies with individual spacer sequences. Nucleic Acids Res. 2015;43: 10831–10847.

32. Künne T, Kieper SN, Bannenberg JW, Vogel AIM, Miellet WR, Klein M, et al. Cas3-Derived Target DNA Degradation Fragments Fuel Primed CRISPR Adaptation. Mol Cell. 2016;63: 852–864. doi:10.1016/j.molcel.2016.07.011

33. Semenova E, Savitskaya E, Musharova O, Strotskaya A, Vorontsova D, Datsenko KA, et al. Highly efficient primed spacer acquisition from targets destroyed by the Escherichia coli type I-E CRISPR-Cas interfering complex. Proc Natl Acad Sci. 2016; 201602639. doi:10.1073/pnas.1602639113

34. Westra ER, Semenova E, Datsenko KA, Jackson RN, Wiedenheft B, Severinov K, et al. Type I-E CRISPR-Cas Systems Discriminate Target from Non-Target DNA through Base Pairing-Independent PAM Recognition. PLoS Genet. 2013;9. doi:10.1371/journal.pgen.1003742

35. Blosser TR, Loeff L, Westra ER, Vlot M, Künne T, Sobota M, et al. Two distinct DNA binding modes guide dual roles of a CRISPR-Cas protein complex. Mol Cell. 2015;58: 60–70. doi:10.1016/j.molcel.2015.01.028

36. Stern A, Keren L, Wurtzel O, Amitai G, Sorek R. Self-targeting by CRISPR: gene regulation or autoimmunity? Trends Genet TIG. 2010;26: 335–340. doi:10.1016/j.tig.2010.05.008

37. Vercoe RB, Chang JT, Dy RL, Taylor C, Gristwood T, Clulow JS, et al. Cytotoxic Chromosomal Targeting by CRISPR/Cas Systems Can Reshape Bacterial Genomes and Expel or Remodel Pathogenicity Islands. PLoS Genet. 2013;9: e1003454. doi:10.1371/journal.pgen.1003454

38. Luo ML, Mullis AS, Leenay RT, Beisel CL. Repurposing endogenous type I CRISPR-Cas systems for programmable gene repression. Nucleic Acids Res. 2014;43: 674–681.

39. Semenova E, Kuznedelov K, Datsenko KA, Boudry PM, Savitskaya EE, Medvedeva S, et al. The Cas6e ribonuclease is not required for interference and adaptation by the E. coli type I-E CRISPR-Cas system. Nucleic Acids Res. 2015;43: 6049–6061. doi:10.1093/nar/gkv546

40. Rath D, Amlinger L, Hoekzema M, Devulapally PR, Lundgren M. Efficient programmable gene silencing by Cascade. Nucleic Acids Res. 2015;43: 237–246.

41. Li R, Fang L, Tan S, Yu M, Li X, He S, et al. Type I CRISPR-Cas targets endogenous genes and regulates virulence to evade mammalian host immunity. Cell Res. 2016;26: 1273–1287. doi:10.1038/cr.2016.135

42. Müller-Esparza H, Randau L. Commentary: Type I CRISPR-Cas targets endogenous genes and regulates virulence to evade mammalian host immunity. Front Microbiol. 2017;8: 319. doi:10.3389/fmicb.2017.00319

43. Amlinger L, Hoekzema M, Wagner EGH, Koskiniemi S, Lundgren M. Fluorescent CRISPR Adaptation Reporter for rapid quantification of spacer acquisition. Sci Rep. 2017;7: 10392. doi:10.1038/s41598-017-10876-z

44. Wu X, Scott DA, Kriz AJ, Chiu AC, Hsu PD, Dadon DB, et al. Genome-wide binding of the CRISPR endonuclease Cas9 in mammalian cells. Nat Biotechnol. 2014;32: 670–676. doi:10.1038/nbt.2889

45. Kuscu C, Arslan S, Singh R, Thorpe J, Adli M. Genome-wide analysis reveals characteristics of off-target sites bound by the Cas9 endonuclease. Nat Biotechnol. 2014;32: 677–683. doi:10.1038/nbt.2916

46. Duan J, Lu G, Xie Z, Lou M, Luo J, Guo L, et al. Genome-wide identification of CRISPR/Cas9 off-targets in human genome. Cell Res. 2014;24: 1009–1012. doi:10.1038/cr.2014.87

47. Leenay RT, Maksimchuk KR, Slotkowski RA, Agrawal RN, Gomaa AA, Briner AE, et al. Identifying and Visualizing Functional PAM Diversity across CRISPR-Cas Systems. Mol Cell. 2016;62: 137–147. doi:10.1016/j.molcel.2016.02.031

48. Jackson RN, Golden SM, van Erp PBG, Carter J, Westra ER, Brouns SJJ, et al. Crystal structure of the CRISPR RNA-guided surveillance complex from Escherichia coli. Science. 2014;345: 1473–1479. doi:10.1126/science.1256328

49. Zhao H, Sheng G, Wang J, Wang M, Bunkoczi G, Gong W, et al. Crystal structure of the RNA-guided immune surveillance Cascade complex in Escherichia coli. Nature. 2014;515: 147–150. doi:10.1038/nature13733

50. Jung C, Hawkins JA, Jones SK, Xiao Y, Rybarski JR, Dillard KE, et al. Massively Parallel Biophysical Analysis of CRISPR-Cas Complexes on Next Generation Sequencing Chips. Cell. 2017;170: 35–47.e13. doi:10.1016/j.cell.2017.05.044

51. Xiao Y, Luo M, Hayes RP, Kim J, Ng S, Ding F, et al. Structure Basis for Directional R-loop Formation and Substrate Handover Mechanisms in Type I CRISPR-Cas System. Cell. 2017;170: 48–60.e11. doi:10.1016/j.cell.2017.06.012

52. Xue C, Whitis NR, Sashital DG. Conformational Control of Cascade Interference and Priming Activities in CRISPR Immunity. Mol Cell. 2016;64: 826–834. doi:10.1016/j.molcel.2016.09.033

53. Severinov K, Ispolatov I, Semenova E. The Influence of Copy-Number of Targeted Extrachromosomal Genetic Elements on the Outcome of CRISPR-Cas Defense. Front Mol Biosci. 2016;3: 45. doi:10.3389/fmolb.2016.00045

54. Luo ML, Jackson RN, Denny SR, Tokmina-Lukaszewska M, Maksimchuk KR, Lin W, et al. The CRISPR RNA-guided surveillance complex in Escherichia coli accommodates extended RNA spacers. Nucleic Acids Res. 2016;44: 7385–7394. doi:10.1093/nar/gkw421

55. Heussler GE, O’Toole GA. Friendly Fire: Biological Functions and Consequences of Chromosomal Targeting by CRISPR-Cas Systems. J Bacteriol. 2016;198: 1481–1486. doi:10.1128/JB.00086-16

56. Pul Ü, Wurm R, Arslan Z, Geißen R, Hofmann N, Wagner R. Identification and characterization of E. coli CRISPR-cas promoters and their silencing by H-NS. Mol Microbiol. 2010;75: 1495–1512. doi:10.1111/j.1365-2958.2010.07073.x

57. Reimann V, Alkhnbashi OS, Saunders SJ, Scholz I, Hein S, Backofen R, et al. Structural constraints and enzymatic promiscuity in the Cas6-dependent generation of crRNAs. Nucleic Acids Res. 2017;45: 915–925.

58. Smith GR, Kunes SM, Schultz DW, Taylor A, Triman KL. Structure of chi hotspots of generalized recombination. Cell. 1981;24: 429–436.

59. Blattner FR, Plunkett G, Bloch CA, Perna NT, Burland V, Riley M, et al. The complete genome sequence of Escherichia coli K-12. Science. 1997;277: 1453–1462.

60. Cherepanov PP, Wackernagel W. Gene disruption in Escherichia coli: TcR and KmR cassettes with the option of Flp-catalyzed excision of the antibiotic-resistance determinant. Gene. 1995;158: 9–14.

61. Stringer AM, Singh N, Yermakova A, Petrone BL, Amarasinghe JJ, Reyes-Diaz L, et al. FRUIT, a scar-free system for targeted chromosomal mutagenesis, epitope tagging, and promoter replacement in Escherichia coli and Salmonella enterica. PLoS One. 2012;7: e44841.

62. Baba T, Ara T, Hasegawa M, Takai Y, Okumura Y, Baba M, et al. Construction of Escherichia coli K-12 in-frame, single-gene knockout mutants: the Keio collection. Mol Syst Biol. 2006;2: 2006.0008. doi:10.1038/msb4100050

63. Guzman LM, Belin D, Carson MJ, Beckwith J. Tight regulation, modulation, and high-level expression by vectors containing the arabinose PBAD promoter. J Bacteriol. 1995;177: 4121–4130.

64. Bonocora RP, Fitzgerald DM, Stringer AM, Wade JT. Non-canonical protein-DNA interactions identified by ChIP are not artifacts. BMC Genomics. 2013;14: 254. doi:10.1186/1471-2164-14-254

65. Bonocora RP, Smith C, Lapierre P, Wade JT. Genome-Scale Mapping of Escherichia coli σ54 Reveals Widespread, Conserved Intragenic Binding. PLoS Genet. 2015;11: e1005552. doi:10.1371/journal.pgen.1005552

66. Singh SS, Singh N, Bonocora RP, Fitzgerald DM, Wade JT, Grainger DC. Widespread suppression of intragenic transcription initiation by H-NS. Genes Dev. 2014; doi:10.1101/gad.234336.113

67. Fitzgerald DM, Bonocora RP, Wade JT. Comprehensive Mapping of the Escherichia coli Flagellar Regulatory Network. PLOS Genet. 2014;10: e1004649. doi:10.1371/journal.pgen.1004649

68. Bailey TL, Elkan C. Fitting a mixture model by expectation maximization to discover motifs in biopolymers. Proc Int Conf Intell Syst Mol Biol. 1994;2: 28–36.

69. Stringer AM, Currenti SA, Bonocora RP, Petrone BL, Palumbo MJ, Reilly AE, et al. Genome-Scale Analyses of Escherichia coli and Salmonella enterica AraC Reveal Non-Canonical Targets and an Expanded Core Regulon. J Bacteriol. 2014;196: 660–671.

70. McClure R, Balasubramanian D, Sun Y, Bobrovskyy M, Sumby P, Genco CA, et al. Computational analysis of bacterial RNA-Seq data. Nucleic Acids Res. 2013;41: e140.

