## Supplementary Materials for "Determining the specificity of Cascade binding, interference, and primed adaptation *in vivo* in the *Escherichia coli* type I-E CRISPR-Cas system"

**Table S2. Lists of regions used to search for enriched sequence motifs.**

Regions used to identify motif shown in Figure 1C

```
>lacZ_Cse1_28747_28847
TGCTGCCAATTTTAGCGTTGGCGTTAACGTCATGCTTAAGCTGCTGGAGAAAGCAGCCAAAGTGATGGGT
GACTACACCGATATCGAAATTATTGAAGCAC
>lacZ_Cse1_106446_106566
AGCTGATTAAGAATTGACTGGAATTTGGGTTTCGAGGCTCTTTGTGCTAAACTGGCCCGCCGAATGTATA
GTACACTTCGGTTGGATAGGTAATTTGGCGAGATAATACGATGATCAAACA
>lacZ_Cse1_113326_113426
GATTGTAGCGTAAAAAAAGACAATTTCGCAGTCTTGCGCCGCATTGATTAGTGCGTATGATAGCGTCACT
GGAGTTGCGCTCTTACCCTTATAGCCATTAA
>lacZ_Cse1_165067_165167
TGAGCCAGACATGACCATCAGCAAGAACGAGATGGTGAAGCTGCTGGAGGCGACCCAGTATCGTCAGGTG
TCGAAAATGACCCGTCCTGGCGAATTTACCG
>lacZ_Cse1_176583_176683
ATTGCCGATGTCGTTTGGAGCAAAATATGAGTGATGACGTAGCACTGCCGCTGGAGTTTACCGACGCAGC
AGCCAACAAAGTTAAAAGCCTGATCGCTGAC
>lacZ_Cse1_201970_202088
AGCGCAAGGTTAATCAACAAGACTAACGTTCCATCTTTTGTTCGCCAAACTTTACGGCCTGTCTCATTCT
TACGATTGCGGCAGGCCGTGTTATTATTGTCGTTTCTTATATTTTGACA
>lacZ_Cse1_229798_229918
TATGGGGGAAGGTTACTCAGTAATTCCTTCTTCTACTAAACGTAAAAACCTGGAAAGTAATCTTAAGGCA
CAAAATTTACAGCTTGATGCCGAAGATAAAAAAGCGATCGCCGCACTGGAT
>lacZ_Cse1_234854_234954
GATTGCTGGGGCGATTTGCCCTGGGGAAAGCTTTATCGCAAGGCGCTGGAGCGCCAGCTCAACCCGTGGT
TCACTAAAATGTATGGTTTTCATCTGCTTAA
>lacZ_Cse1_240270_240370
ACAAATGCTGATATTGGAAATATCTGATTTGCAAATTATCGTGTTATCGCCAGGCTTTAGGAGGTTAATA
ACATGGGCAGGATAAGCTCGGGAGGAATGAT
>lacZ_Cse1_247959_248059
TTATAGGCATCATGTTGATTATATTTATATAAATCCAGTAAAGCATGGTTGGGTAAAGCAAGTGAGTGAT
TGGCCATTCTCAACGTTCCATCGCGATGTCG
>lacZ_Cse1_263064_263190
GGCCTGATAGGGCTTCGCTCACTATACATCCTTGGCTGCAGGTTTAGTTGTACACCACTCCTAAATTTAA
TGTGTTGGCAATGTGTTCAATAAAGCTCGAACAAATTAGCTCATTATGATCGGTTAA
>lacZ_Cse1_278042_278142
TATCAACGTATCCGTCGCTGAAGGTTGCCGGGAACAGGTTGGCATCGCCCATAAAGCTCGCCATAAAGCG
GGAGGCGGGCTGGCGGTAAAGATCCTGCGGT
>lacZ_Cse1_290134_290234
AGTCGAGTCTTTTTCGAAGATGAGCGCGATGTTTTTACCGGTAAGCTTCTGTACTTCCTTGCCATTTTTT
TTATCGGCTTTGAGCTGTGCGGCAAGGGTCA
>lacZ_Cse1_295123_295223
GGAGTTTGAATGCGCTCATTTTGCCACCTCAGTGATTGAGATATTGGCAAAGAAAGCTTTAAGCATTATT
GAATTAGTAGGGGTAGTAATGAAGGCAGAAG
>lacZ_Cse1_366296_366416
CATGGTCATAGCTGTTTCCTGTGTGAAATTGTTATCCGCTCACAATTCCACACAACATACGAGCCGGAAG
CATAAAGTGTAAAGCCTGGGGTGCCTAATGAGTGAGCTAACTCACATTAAT
>lacZ_Cse1_439479_439579
ATATGTTCTTCGGTGCGTTTGAGCAGCTCTTTAATTGGCGGCACGCCAGAGAAAACTTTTTTCCCGCCTT
CGCGCAGTGAAGAGTAAAGCTTACCGGAAAG
>lacZ_Cse1_516788_516888
GGTGTTATTTATCTGCTTTAATGCGGCGTGGAACGCGCAAAAACGCAGTAAATATATTGCTAAAGCTTTT
ATCTCATCGTTTATTGCCATTACGGTCGGGG
>lacZ_Cse1_557454_557559
```

```
AGAATGCCATCGATGGTGTTGTCGGCATTCAGCGTATCGATAAGCTCCAGCAGCTCCGCTTCGCTGGTGG
TTTCCGGGAGGTCATAAGAGCGGGAGACGAACCCGA
>lacZ_Cse1_582723_582823
TTGATACTCCCAAAACCCACCAGTCATAAAGCTTTAGGGTAAGTGGTGTGTAAATTCTAGCCCCATCATC
TGTGTTTTTTTATTAATTTCACCATGTTATA
>lacZ_Cse1_605994_606094
CAAAATACTCAATAGCGCCCAGCACTAAAAACCACAGACAAAACAATAAAGTGTAAAGCTGACTAAGATC
CATCAGATGGAACATGGTCACCAGTTTTTGT
>lacZ_Cse1_636764_636864
TTTAAAGAAGATTTATTATAGGAAATGTAAAGCTTTATTGAAGGTAACGGATGTTCTAGTTTTATCTCTT
TTAAGTTAAGAAAGTCACGGTAGGAATTATA
>lacZ_Cse1_655952_656063
CAATGAATGTCAGCATAATTTTTCCTGTCTCCAGGCCCCAAAGTAAATAATAAAAAATTCCTAAAGCTTA
AGGAAAAAATATGCCCAATAAATTGGCGATGAATGCTGATTA
>lacZ_Cse1_657198_657298
TTAGTAAAGTTACACTGGACAAAGCGTACCACAATTGGTGTACTGGTAACCGACACAGCATTTGTGTCTA
TTTTTCATGTAAAGGTAATTTTGATGTCTAA
>lacZ_Cse1_657405_657505
AATGGTTTTAAAACTCTTGCTGAAGGTCAGCGCGTAGAGTTCGAAATCACTAACGGTGCCAAAGGCCCTT
CTGCTGCAAACGTAATCGCTCTGTAAGATAC
>lacZ_Cse1_780942_781047
AAGCGTGTAGGATAACGTTGCGTCAGCAACGGCCCGTAGGGCGAGCGAAGCGAGTCATCCTGCACGACCC
ACCACTAATGACGGTGGGTTCGGTGGAAGTAGTTTG
>lacZ_Cse1_933339_933439
TGCCTTACTAAGCTTTAACCCTTCGGACCCCAGCTGGTCGCAAACGGCCTGGCATGAACCTATCCATAAT
TTAGGTGGGATGCCCGGTGCGTGGTTGGCAG
>lacZ_Cse1_941902_942002
CGATAAATATTGCGTTGGCTACGATGAGAAAACCCTGCCAGCCAGTGCGCCGAAAAATGGCCACTATAAA
GCTTATATTCTGGGTGAAGGGCCAGATGGCG
>lacZ_Cse1_971853_971953
GACAGCCAGGGAAACCCAATGCCGTTAATGGCAAGAAGCTTTAGTACGCCCGGAAAAGCCCGACAACTGG
CATGGAAAACCACCATGCTGGCAAGAAGTGG
>lacZ_Cse1_982571_982671
ATGACACGCCGAACCACAATCTGTTCAAGCGTGATACACGCGCATTGAGCTCAGGCTGTGTACGAGTGAA
TAAAGCTTCCGATCTGGCGAATATGCTGTTG
>lacZ_Cse1_1070347_1070447
TGTTCATAGTTTGCTCCTTACAACAGCCCGCAGGCTTCTTCAAAGGACAGACGTGGCAGGCGCGCATAAA
GCTTGCTGCTATCGCCATAGCCGATATTAAT
>lacZ_Cse1_1092822_1092922
TATATCTATTAATAATAGAAAGGGATCTACAACCTACAGATTGGTGTAGCTTTATGGAAAAAGACTATTT
GAGAATTAGTAGTACTGTATTAGTGAGCTTA
>lacZ_Cse1_1093954_1094054
CGAAGAGTTTGGCGTCTTGCTGACGGACATCGATACTGAACGTGCGAAAGCTTTAGCGGAAAGGATTCGG
GAAAATGTTGAGCGTTTAACTGGCGATAATC
>lacZ_Cse1_1182976_1183085
GGCAGAATTTCATAACTTGGTCCGGTGACGAACGAAGAAACTACCACATCGTCCATCGACAGGGTAAAGC
TTAACACCCAGCCCGCCGCCACCGCTGGCATTGCCAGTGG
>lacZ_Cse1_1195007_1195107
GCCAATTACAAATCATTAACAAAAAATTGCTCTAAAGCATCCGTATCGCAGGACGCAAACGCATATGCAA
CGTGGTGGCAGACGAGCAAACCAGTAGCGCT
>lacZ_Cse1_1261957_1262061
TGTATTAAAGAGCGCGATGCAACGTCTGGAACAAGGTGACGTTGTCACCGAAACTCAGCTTGCCCGGCTT
AAAGCATGGCTCTGTGCAATGGGGAAAGATTAGCG
>lacZ_Cse1_1267891_1267991
GATGGAAGAAAATCAGATGGAGCTGGCGTTACGCACCAGCGAAGCTTTATTACAATTCAACCCTGAAGAT
CCCTATGAAATTCGCGATCGCGGGTTGATTT
>lacZ_Cse1_1280269_1280369
```

GGGCTTCTATCATTGAAGACGCCGATAAAGCGAAAAGCTTTAAGCAGGCGCGTGGACGCGGTGGATTTGT
TCGTTCTTCCTGGCAGGAGGTGAACGAACTG
>lacZ_Cse1_1311785_1311885
ACGGCACCACCGAAATTCAGTAAGCAGAAAGTCAAAAGCCTCCGACCGGAGGCTTTTGACTATTACTCAA
CAGGTAAGGCGCGAGGTTTTCCTTCAGGATC
>lacZ_Cse1_1326656_1326756
TCGAAACAAGCTTTAACCATAAAGATAGTGTAGTATGTTGCGCCTCAAAGCCAGGCCGATAAACGTCGAG
TCGTTTACTTAAGGCCTGAAGAGTTCAAACA
>lacZ_Cse1_1435610_1435710
TCTGTTCATAAAACCTCCTGTTTTAGTATCCGCATAAAGTGTAACGCCAGATGACACTTTTTGTGTAATG
ACGGAGTTCACATTTTTAATTTAGATCAAAG
>lacZ_Cse1_1449286_1449397
GAACGGTTCGGATATTTGCCTTCCGGGAAACGCTCGCCAGGATGATAACGCGTTACCCAGAGCTGCTTGT
CCATAAAGCTTAAACGATGATAGATCCACTCGTCCGGCGCGA
>lacZ_Cse1_1450879_1450985
GAGATCTCAGTAAAACGGGTATTGGGTTTGAAGTCCGCGGAAGCTTTAACAATTTCAACGGCCTGTTTAA
TTTCGTCCGCAGTTAGCGCATTAAGTGGGTGAGGGCG
>lacZ_Cse1_1467756_1467856
ATCTGAATGGAGATACGACAATCAGCGGAGAATTCCCTCTGGGGTTTGCCGGGGTTATTCGGGTACAGGA
TAAAGCTTTGCTGGAAATTGGCAGTGGCGCT
>lacZ_Cse1_1479912_1480024
ATCACGACACGTTGGGTGAAGAGAACAGGGCGCTGGCATTTGAAAAAATGCAAAGCTTTATTAGTCGTTT
ATATGCTAACAAATCGCAAAAATTTGATTATCAGCATGAAGAC
>lacZ_Cse1_1529856_1529956
GCAACTCTTCGGGTTAGATGGTATAAAAGCAGATAATCCAATAAAAGCAAAGCAAGGGAGTTCAGATGCT
TCAAATTATTAGAGGCAAACTTGTCATTTTT
>lacZ_Cse1_1574248_1574348
CTGCAGACGTTGATTGAACAGCTGAACTAACATGCTCCATTCAGCTTGTTCGATAAAGCTTTGCTCATCG
TTCACTTGTACCATTGGCAGGCGATAATAGA
>lacZ_Cse1_1818245_1818345
AGTAACGAAATCAGCATCTTTCAATGCTTCGCGGCGATCCAGCGTTTTATAAAGCTTCATCGGGACGCCA
GCGTTATCAATCATCCGTTGGCAGAGATCGA
>lacZ_Cse1_1873422_1873522
CTCTTCCGGTATTTACGCGATGAAAGAAAACGATACGTTACATGATGCGTATAAAGCTTCCGATGAGCGT
TTATATGTCAATAAGCAGAACAAAAACAGCC
>lacZ_Cse1_1893674_1893774
GTATAAAGTTGAAATTAAGATTGTGGCTGCGGTGTAAGCTTTATCGAAGCAAAATAAGTCAGACGATAAT
TTATCGATAATACTGGTCGGTTTTACATAAA
>lacZ_Cse1_1979650_1979750
CAGACTAATATTTACCTGTTTGACCGAGTTGGGATTGCGTCGTTTCTCCATTAGGAGTAAAGCTTTAATG
TCACCTGAAGTTCACAGAATAAGGAACAGGT
>lacZ_Cse1_2019322_2019422
TTCAGTTAGGGCAAAGCAACATCAGTGGCGAAAGCTTTAGTGGTCAGCAGCAGGCCGCTTCCCAGCAACA
GCAAAGCCAACGCACAGCAAACCATGAACCT
>lacZ_Cse1_2049165_2049265
CTATATATACTACCAACCACGACGGCAATTTTTACGCAAGCTTTACCGCTACAAAAGCCGGGGTTTATCA
ATTGACGGCAACCCTCGAAAATGGCGATTCG
>lacZ_Cse1_2049729_2049829
TGACATTCACGCTACCGGCAGATGTGAGCGCAAGCTTTACTCTCGGACAAGGCGGTTCCGCCATTACTGA
TATCAACGGCAAGGCTGAAGTTACACTGAGC
>lacZ_Cse1_2050859_2050975
AACACTGATTTAAGTACCTTAAAGGCAACGGTTGAGGATGGCAGTGGTAACCTGATCGAAGGTCTCACTG
TGTACTTCGCCTTAAAAAGCGGCTCTGCCACATTAACGTCATTAACA
>lacZ_Cse1_2053236_2053336
ACAACAAAGTCTGTTTTTTCAATATTGAAATTATAAGCTTTATAATCGTAGTAACTAAATGCAATACGTC
TCAACAATGATGCTAAAAACATACCTAACCT
>lacZ_Cse1_2060508_2060608

TCGCAATTTCCTGTGGCGTCCCCTGGATTAGCTCGAGCCGAACCTCCGGGAAAAGTTCGCGAAAAGCTTT
AATGACCTCTGGCAAGCTATAACGTGCCTGA
>lacZ_Cse1_2167084_2167199
CTGCATTTATTAAGGTTATCATCCGTTTCGCTGAAAAACATAACCCATAAAATGCTAGCTGTACCAGGAA
CCACCTCCTTAGCCTGTGTAATCTCCCTTACACGGGCTTATTTTTT
>lacZ_Cse1_2177046_2177205
GTGCCAGGCGTTCCGGCGATGATGACCGGCGCATGCAGGTTGGCAGCGGTTTCTACCACCACTTGCATCG
TTTCGAGATTGTGAATATTGAATGCCGGAACCGCATAACCGCCGCGCTGTGCGTTGTTCAGCATCTGCTT
TGTCGATACCACGTACATTT
>lacZ_Cse1_2271786_2271886
TAACATTGCGCCGGACCATCTGCACAGCGAAAGCTTTAAAGCAGATATCCAGAAACTGCTCACTTCCCTG
CCCGCACACCATTTCCAGATTGTGCTGGAAA
>lacZ_Cse1_2478238_2478338
AATTGCCAAAAAAACCAATACCTCATTATTAAAGCTTGCCATTCCGCTATGTACCGCATTGATGGCAGTG
CACTGCGTGGTTCCTCCACATCCGGCTGCTT
>lacZ_Cse1_2518192_2518292
CATGCCATGCAAGCGCTCTCCCAGCTGAGCTATAGCCCCGTCACGTAAAGCTTGTCGAGTTGACGGGCGG
CATCATATGAATTCCGCCCGAATGTGTCAAC
>lacZ_Cse1_2525945_2526045
GGATCCCCTTCCCGCCCAGCCATTAGCGCATTGCCCTGGTCTCGTACCACGTCCCATAAAGCTTCAGCGG
CGGTGGAAAGTGAACGGTTCTTCCTGCGCAC
>lacZ_Cse1_2527856_2527956
GATGGTCTGTGCGGTGCTGGCGCGTCGATACAAACGCCAGACCGAACAGTTACAGGCGCAGCAGGAAAGC
AGCGCCGATAAAGCTTAAAGCGGACGCTTCA
>lacZ_Cse1_2533245_2533345
GCGGCGTTGAAACTACAAGAAGATGAAAGCTTTACCAACAAGAATATTGTGGTTATTCTACCATCATCGG
GTGAGCGTTATTTAAGCACCGCATTGTTTGC
>lacZ_Cse1_2554979_2555090
TCCGTAAATTCACCATGAAACTTGCCCGGATGCAGCTGATCGTAAATCTGCTGCCAGGCGGTAATCGTTA
AAGCATGTTCATAGACATCCGTTGTCTGTCGTTGATGAACAT
>lacZ_Cse1_2634138_2634238
TCTACCCCTTTTTGCAAAAAATGCTTGCTATCCCCGAAGGGCGGGTTACTATCGACTGAATAACCTGCTG
ATTTAGAATTTGATCTCGCTCACATGTTACC
>lacZ_Cse1_2659023_2659145
CACGGTATCCGCCGCCGCGTCCCACGTCTCGTTGTCTAACTGTTCAACCATCAACTGATGGCGGGTATCA
AACATCTTTTTCACACGTTTGATAAAGCTTTCCAGCCGCGCTTCATCTTTCGC
>lacZ_Cse1_2707698_2707798
AACCGGCGCGTGAAGCGCAGCTGCTGCATGAAGCTTTAACATCACAACTGACCAGCGCCTGCCCGTTTTG
CCAGGAGACGACGGTAGCCCACTCTTTGATC
>lacZ_Cse1_2737078_2737178
GATTTTCATCACACATTTTGACATCAGGAACGGTATGCTGAATTCACCAAGACGGGAAGACAAGAGGTAA
AATTTATGACAATGAACATTACCAGCAAACA
>lacZ_Cse1_2755568_2755668
CTGGTATACTTACCTTTACACATTGGGGCTGATTCTGGATTCGACGGGATTTGCGAAACCCAAGGTGCAT
GCCGAGGGGCGGTTGGCCTCGTAAAAAGCCG
>lacZ_Cse1_2755718_2755818
CTGCTTAGAGCCCTCTCTCCCTAGCCTCCGCTCTTAGGACGGGGATCAAGAGAGGTCAAACCCAAAAGAG
ATCGCGTGGAAGCCCTGCCTGGGGTTGAAGC
>lacZ_Cse1_2810847_2810947
ATCCTTCTGACTCGTCTTTGCATGCACATGCAAAGCAAGCTGCTGGAGAACCGCAATAAAATGCTGAAGG
CTCAGGGGATTAACGAGACGTTGTTTATGGC
>lacZ_Cse1_2870369_2870469
CAATTCCAGCTTAATTGTTGCGGATACAGCAGCATCGCTCTGCGCGCGGCTTCAACTTTTTCGCGCACCA
GTAAAGCTTGTAATTCAGTTTCTGCGGCGAC
>lacZ_Cse1_2883375_2883475
GTCCACAGCAAGAACATTTACCAATCCCAATGGGATCGCATAATTCAATATGCGCTGGTTGCCAGAATAG
ACCACGGACAAACCCAATTGACGAAGCAGGT

>lacZ_Cse1_2884220_2884320
AATACGGCGTAATTAATTCAATAATCACATTCACTGCAAAAATATATTCATTGGTTTAATACAATTAACC
TATACATATATTAAGATGTGTTGAATTGTTT
>lacZ_Cse1_2927412_2927512
ATATGGCTAATCTGAAGCTTTACCCGGATCAACCTGTTGAAGTGCTGGCTGCCGACCTGCGCCGTGCGTT
CTCCGGTATTGTGGCGGGTAACGTAAAAGAA
>lacZ_Cse1_2942730_2942830
GCGTGTTCTGGTGAACTTTTGGCTTACGGTTGTGATGTTGTGTTGTTGTGTTTGCAATTGGTCTGCGATT
CAGACCATGGTAGCAAAGCTACCTTTTTTCA
>lacZ_Cse1_2968972_2969081
ATCACCGGACTACCTCAAAATAAAGCTTTATATACGAATGATTGTTTCATACTCCAGGAAGACGGTAAAC
CACTCTCTGCAGGGCATTACACACTAATAACAATTGAATA
>lacZ_Cse1_2989810_2989910
GAACCTCGTTAATCATAAAACAATAAAATTAAATATTCTCGCAGTATATGGCAGTCTAAAGCATCAAAGA
TTTGATCAACATCTTTCATTTTAGACATCTC
>lacZ_Cse1_2998964_2999064
TATTACGGCGAAGATAAATTGGAGCGGGCGAAGGGAATCGAACCCTCGTATAGAGCTTGGGAAGCTCTCG
TTCTACCATTGAACTACGCCCGCTTCGAGAT
>lacZ_Cse1_3035337_3035437
GATTTGTTATCTTCCATCGCCTGTTTCTCGGCATTTTTCTTCTGCATCTCCAGTTCATAAAGCTTCGCTT
TCATCTGCTTCATGGCCTGATCTTTGTTCTT
>lacZ_Cse1_3045032_3045132
TCGCGTTCACGTAAGAAGAGCTTTAACTGGTACAAAGAGGTGATTGCCAGCAACGGCGAGAAGCTTTAAG
TCGATGAAGTACCGGATGCAATACTTGTTGC
>lacZ_Cse1_3073736_3073867
AATCCAGCAAATCGTCCGGGGAAACCTTACCTGTCGCAAACTGCGACTGATTGGTTAATTGTCGAACATT
TAATCGACTGAAACGCTTCAGCTAGGATAAGCGAAACGTGGAATAAAAGGAATGTTTGTCCA
>lacZ_Cse1_3073876_3073976
AGACATTTATCTGACTCACATCACACTTTTATCCCCTTTTGTGGGAAGCTTTATTCCAGGCTGGCGTAAT
AATAACCCTACAATAACTGGAATAAATTGTC
>lacZ_Cse1_3095004_3095104
CGGGCCAGGCGCTGCACAGGTGTAGATCCGAGACGTTATGCTTTACACTAAGGGCCACAATTTCTTCCAT
ATTCATACTAAGATCCTCGGAAAATGAACGA
>lacZ_Cse1_3104246_3104346
AAAAGAAACTCAACATGATGAATGCCGAGCACCGCAAGCTGCTTGAGCAGGAGATGGTCAACTTCCTGTT
CGAGGGTAAAGAGGTGCATATCGAGGGCTAT
>lacZ_Cse1_3121305_3121427
ATGCTTGATATACTTAAGGTTGTAATAAGCAAAAGAGGACTGAACTGTAAAATATAGGCGTTATACTTTA
CAGCAACAGTACGCCGCTAACGCAATTGCTACCTCTGGCATAACAAGTATATC
>lacZ_Cse1_3210870_3210970
GAAGTTCGTCGTCGTGAGTTCTATGAAAAACCGACTACCGAACGTAAGCGCGCTAAAGCTTCTGCAGTGA
AACGTCACGCGAAGAAACTGGCTCGCGAAAA
>lacZ_Cse1_3299968_3300084
ACACCCTGGTTTCTTTACTTGATCCGTACCGCCGACAATAAGCTTTATACCGGGATCACCACGGATGTCG
AACGCCGCTATCAGCAGCACCAAAGCGGCAAAGGGGCGAAAGCACTG
>lacZ_Cse1_3307901_3308014
AACTCATCATGAAATTGATCAGCAATTTTCATTGAAAAGTGTGAACCGGCTCAAAGTAGGTGTATTAACG
AACAACAACGCCCTCACCCGTTAAGGTGATGGCAATCAAAAAAG
>lacZ_Cse1_3318284_3318384
CCGCGCCTGAAACGTGGCAAATTCTACTCGTTTTGGGTAAAAAATGCAAATACTGCTGGGATTTGGTGTA
CCGAGACGGGACGTAAAATCTGCAGGCATTA
>lacZ_Cse1_3336012_3336129
ACTTTATCCATCACGATATGTGCACCTTTCAAACGACCATCGACGGAAGCTTTAACGTAACCTTCTTCCA
GTTTGATGGTCGCGCCTAATTGTTCGAGGCCAGAAATGTGTAGATCAA
>lacZ_Cse1_3350549_3350649
TCATTTAAGTTTTGCTATCTTAACTGCGTGCGGCCTGAAAAACAGTGCTGTGCCCTTGTAACTCATCATA
ATAATTTACGGCGCAGCCAAGATTTCCCTGG

```
>lacZ_Cse1_3358237_3358337
CTGGACGGTTTCACCGCCAAACAGCAGAAGCTGGCGCTGTCGAAGCTGCTGGAGACTGCCGAACCGCATC
CAGGTAAGGCACTCTACTGCACCGAAAACAA
>lacZ_Cse1_3377502_3377602
CGACTCTCTTTGTTCGAAAATCAAACAAAAAATGAGCAATACCCGACATTTGGGCAGAAAATTGGATGAT
AGTTTACCAGATTTTGCGACCATTGTGGTGA
>lacZ_Cse1_3401668_3401768
GTACTTACCAGCGTCAAACCAGCCTGGTTGACGGCTACCCGTACCCCTAAAGCATTCACTAAACGAATAA
CAGGTTGTAAACGACTGATATGTTGACCTAC
>lacZ_Cse1_3441023_3441123
CTGCATTGTGTCCTCTCTTTGGTACTAAGCTTTACTTGGAGTAAAGCTCGACGATCAGGTGTTCGTTAAT
GTCCGCAGACAGATCAGAACGCTCCGGCTTA
>lacZ_Cse1_3535819_3535919
ATAACGTACGGGCAAATGAACTTCGTGGCGAGAAGCGCAATCGCCTCATGCTTTAGAGCCGTCCGGTACA
AAGACGTAGCCCAGACCCCAGACGGTCTGAA
>lacZ_Cse1_3563530_3563630
TGATCAGGTCAGATTTCAATCTGGCCTGAGACTGATGACAAACACAAAACTGCCTGATGCGCTACGCTTA
TCAGGCCTACGTGGTTTATGCAATATATTGA
>lacZ_Cse1_3631277_3631377
TAACGATTAAAGGATCGACAAGAGATAGCCCGGCTAGTTTTAAACTTTTTGCCATTAATTATAGCATGAT
GCTTTTTATTGATAATTGCCGCTAATTCGAC
>lacZ_Cse1_3665713_3665839
CACCGCCATTAAAATAGCGATATTCACCATTAACCATGGTGAGAATATATTTATGTCTTGCATACGCAAT
TAGACAATTCCCATGTAGTGATTGCATAGTTGACTTAATATTACATAAACATATTAC
>lacZ_Cse1_3725571_3725677
CTGATCCTTGCATAACGCTTCCATATTCAACGAACGCAGAGGATCAACCTTTACCCGTTGTGCTTTCTGC
ACGGGTGACGGTATAGAACAACCCACTAGCACCATAA
>lacZ_Cse1_3793078_3793178
CCCCCAATAATATATACATGATTGCCAATCATGAAATCAAAAACCTGTTGTTGAGATGTGACAGTAAGCC
CTTTTTTTATAGTGAAAAGATGCATCTTACT
>lacZ_Cse1_3796253_3796353
CGCTCAGGGGCTGGTAAAAAGCGCGGCGCTGGTGACGCGTCTGGCGCATGGCGTAAAGCATGGCATGGAC
TGGCAAACCGCTCGCGAACCTTTAGCCAGCC
>lacZ_Cse1_3802236_3802336
ATGCCAGGGCTTGGTTGGCCCGATATAATGGATAAAAATAGTATCGTTGGTTACTGGGTTTATAAAGCTT
TCTTTGAGTTGATAATTTAAGCTGAACTGGG
>lacZ_Cse1_3890950_3891067
TGATTTATCTCTTCTGGCTCTATTGCACCATGGGGAATATTCCGCGAGAAAGCTTTAAGGCGATTATCTC
CTCAGGCGGCAACGTTGATTCGCTGGTGAAATCGTTCCTCGGCACCAA
>lacZ_Cse1_3902022_3902122
ACGATTCACCGTCGCTGGCGTATACTTTTTCTTCTCTTCATCTTTATACTTTTTGAGTTCGCTTTGCGTT
TCTCTTAACGCCTTTTCTAATAACTCAAGGC
>lacZ_Cse1_3917674_3917807
ATCATCATCTGATGCCGGTAACGGCAGCAGCTGGCTGATGGTCGGAACCTGAGACATGGTGTTAATAAAT
TTGTTGCTGACAATGTAAAGCTTGTCCAGACGGCCTTCGTCGTAGGCCTGCAACATCACTTTTA
>lacZ_Cse1_3939012_3939136
CAGGTTCCGCAGGATATCGCGGTGATTGGCTATGACGATATCGAACTGGCAAGCTTTATGACGCCACCAT
TAACCACTATCCACCAACCGAAAGATGAACTGGGGGAGCTGGCGATTGATGTACT
>lacZ_Cse1_3940616_3940716
TGTCCTGTTATCGGGTTATCACTTATCAGGTGAGCGTAGCAGCGCCTGACAAGCTTTAAATGCCGCGTCG
CCATCGCTTTGGATAATCGCATCGACAATCG
>lacZ_Cse1_3948030_3948130
GTAACTAATACTCCGCGCCATAACTAGCTCGGTCAAAGAATTAGGAGCGTGCAGGATGGCGGAAAGCTTT
ACGACGACTAATCGATATTTCGACAATAAAC
>lacZ_Cse1_3950314_3950414
AGACAAAATGACAGCCCTTCTACGAGTGATTAGCCTGGTCGTGATTAGCGTGGTGGTGATTATTATCCCA
CCGTGCGGGGCTGCACTTGGACGAGGAAAGG
```

>lacZ_Cse1_3968220_3968320
CATACAGGCCGCTGTTGGCATTGTTATGATGGTGTTCGGCAAGCTTTATCTCAGTAGCCTGGGTTATATC
TTTGGCTCCTGGGAGATGGTGCTCGGACCGT
>lacZ_Cse1_4024813_4024913
GATTGCATAATGACTCTTATCCGTTTAATCGGGGCGCAAGGATAGCAAAAGCTTTACGCTAAGTTAATTA
TATTCCCCGGTTTGCGTTATACCGTCAGAGT
>lacZ_Cse1_4045710_4045810
GCCGGTCTTTTTCCTGCCACAATGAGTTCAGCCATTGCTGAAAATGGCGCTTAAAGCTTTTGTCATTGAT
GTAGTCGCCATGTAACTCATCGGCAATAGGC
>lacZ_Cse1_4118096_4118196
ATCCTGAAGAGTTAATGTTTGTTGTATGCGTGAAAGTCACGGACCTCCACGATGCTTGTAGGCATGCTGT
AAACTTATCGTTAACGAGCAAAAACGAGAAA
>lacZ_Cse1_4154491_4154591
AACGCAAAAAAGCCTTTCATGTCGGCGGTGCCTAAGCCGTAAAGCTTGCCGTCATGCTCCGTCAGTGTAA
ACGGATCGCGCGTCCAGCGACCGTCATCAAA
>lacZ_Cse1_4220261_4220361
ATGTGGAAGATGTTTATGCGTATCGGCGCAGGCAAAGATTTAGCGTACGGTATGGGGAGATGCTTTTTTG
AGTAAAGCTTCCATATAATTTTTCTCCGCAA
>lacZ_Cse1_4323168_4323268
ACGGCCCAGCTCAGAAGCGAGAACCAGCTGCGCTTGGGTGGGGCGATGGTGATGGTTTGCATGTTTGGCT
CCGGTCTGTAGGCCGGATAAGGTGCTTGCAC
>lacZ_Cse1_4369358_4369458
TAAAGATGCCATTGGCATAAATAATAAGAGCGTCCAGATTGATCTCTAAAGCATGAATCACCAAAGTGCT
CACCACACAGCAGCCGAGCACCGCATTGAGG
>lacZ_Cse1_4392261_4392361
CTTTTTTGGGGGGTTGCAGAGGGAAAGATTTCTCGTATAATGCGCCTCCCGTAACGACGCAGAAATGCGA
AAATTACGAAAGCAAAATTAAGTAGTACGCG
>lacZ_Cse1_4427054_4427154
TCGACATCTTGCAATAAATATTTCAGTTCTGCGGTTAACCTCGCACCCGCGTAAGCTTTATTGTATTTCT
GGGCGTGAATATTATCTAATGTAGCTTCATG
>lacZ_Cse1_4438814_4438921
CAGCGGAACTTTCTTCATCCAGCTGCCAGGCACCGAGCGCGACGCCATTGCTATCGACAATAAACTGCGA
CCAGGGATAAAGCTTTTTATTACTCTCCAGACTGCTGC
>lacZ_Cse1_4464735_4464861
GCGCAATTGCGCCCCGAAGAAAATTAAACAATGTCCAGCGTGCCCAGGCGGTATTTCCGCTGATAGATAA
ACGAGGTGAGTACAATCGGGATGATGATGGCGATAGCCATTGCCAGCGCAAACACCT
>lacZ_Cse1_4464919_4465019
TCGCCATAACGCCGTTCAGGCCGCATAGCAATCCTGCCAGACCAGAACCAATCATCGCGCACAGCATCGG
GAAGCGATATTTCAGGTTGATGCCGTACATT
>lacZ_Cse1_4465442_4465552
TTCAATAACGCCCAGCGCCAGTCCGGCTAACAGTGCCGGGATCACCTGCGCCTGATAGCCTACTTTGGCG
ATGCTGAACATGCCAAAGTCCCACACTTCCGGCAGCTGCTG
>lacZ_Cse1_4466070_4466170
AGGACAAAGCGTAGGCGAGTAATACAGTGGCTCACCGTCGCAATATTGCCGCGCCCGCCGACCAGTTCAA
TCAACCGATCGATATCCGTTTGGTTTATTTT
>lacZ_Cse1_4478350_4478450
CAATTTATATGGATGATTATTCATTTGCAAGTCTAAAGCATAAATCTTTGTCACAAAGGTGGAGGCAATG
TCAGTGGTGTGTGACAATAAGAGTATCGGCA
>lacZ_Cse1_4534058_4534158
AGAACCTATTTAATCATCATGTGCAAAACGTGCAAACACACCGCGGTGTCCGCATTCGATTTCGGCGCAT
TGATAATCAGTCCGGCCTGAAAAGGTCGGGG
>lacZ_Cse1_4569408_4569521
CTGGACGAATTTGCCGACCCGACTACCGCACGGAAGCTTTATAACGCCGCGTTCCCGCTGGTGGATGTTA
CTGTCGTGCCAGACGACGAGATTGTGCAGCATCGCAGAGTCGCC
>lacZ_Cse1_4572267_4572367
TTTTATCTAAAATTGATTTATAGTATCGACCTGAAAAAATAGTTGTTGCCGCCTGAGTAACTATACAATA
TTCTGAAAGGTTTTCTTTCAAATTAGAAATG

>lacZ_Cse1_4619516_4619616
GCACACCAACTGTCTATCGCCGTATCAGCGAATAACGGTATACTGATCTGATCATTTAAATTTGAAGCAC
TGAGTACGGAGAACATATGAAACGTGCATTT
>lacZ_Cse1_4638323_4638423
CAGCGGGCCGCAAAAACTCGTTGGGCCGCTCATCGCTATTCCTGTGACCGAGCTTTATACGGTGCAGGAA
GAGGATAAAACCGTGGAGCGGAAACGAAGTT
>lacZ_Cse1_4641425_4641533
CGGCGAAAAGTGATGCAACGGCAGACCAACATCAACTGCAAGCTTTACGCGAACGAGCCATGACATTGCT
GACGACTCTGGCAGTGGCAGATGACATAAAACTGGTCGA

<u>Regions used to identify motif shown in Figure 1D</u>

>araB_Cas5_44103_44225
CGCTGACCGCAGCTTTAGCGCGTTGATCCACTCTGGCAGGGCTGCATTTTGGCCCTGCCGCTGACAGGGA
GCTCTTATGTCCGAAGATATCTTTGACGCCATCATCGTCGGTGCAGGGCTTGC
>araB_Cas5_70064_70167
AAAAAACGGGTATGGAGAAACAGTAGAGAGTTGCGATAAAAAGCGTCAGGTAGGATCCGCTAATCTTATG
GATAAAAATGCTATGGCATAGCAAAGTGTGACGC
>araB_Cas5_94590_94690
AATCAGCGTCTGGACTACTCCGATCGCGTCACGGTGGCGCGTCTGCTGGGGGTGATTGCATGATTAGCGT
AACCCTTAGCCAACTTACCGACATTCTCAAC
>araB_Cas5_131278_131378
ATCGCAATGGTCGGCGAACTGCCACCCGCAGGTGCTGTGAATCCGAGTATAAAGAGGCGGTAGTTTAAAT
TTTGACTAATCTTGGGATTCGTTGAGAAAGG
>araB_Cas5_165060_165160
TCAATCTTGAGCCAGACATGACCATCAGCAAGAACGAGATGGTGAAGCTGCTGGAGGCGACCCAGTATCG
TCAGGTGTCGAAAATGACCCGTCCTGGCGAA
>araB_Cas5_252165_252270
TGAACAAGAGGAGAAAAAGCCCATAAAGGCGCGCTTCCGTCCAAGTGCTGCAAGATTAGAGGAAATTACA
CGCCGCGCTGAACAATATCTTAATGATATGACGGAT
>araB_Cas5_256389_256489
GTGGGATATGGGCGTCGTATTCGTCCCGCCAATCTCCGGTCGCTAATCTTTTCAACGCCTGGCACTGCCG
GGCGTTGTTCTTTTTAACTTCAGGCGGGTTA
>araB_Cas5_262826_262926
ACCTTGTCCACCGTGATTCACGTTCGTGAACATGTCCTTTCAGGGCCGATATAGCTCAGTTGGTAGAGCA
GCGCATTCGTAATGCGAAGGTCGTAGGTTCG
>araB_Cas5_297157_297257
GTATGTTGTTTTGATTGATTGCTCAAGTAGTTAAAAATGCATTAACATCGCATTCGTAATGCGAAGGTCG
TAGGTTCGACTCCTATTATCGGCACCATTAA
>araB_Cas5_304479_304579
ACCGATGGATAAGTCTTAGTCCGCCGAAGGGGGCTTAGCCGGACAGGAATCGCTAATCTTAATGAATTTG
TCGTTATAGACCAGATAGTGATTCCCCGGCT
>araB_Cas5_348718_348818
CACTGACTAAAGAAAATCCATTGCAGATTGTTGGCACCATCAACGCTAATCATGCGCTGTTGGCGCAGCG
TGCCGGATATCAGGCAATTTATCTTTCTGGC
>araB_Cas5_366328_366428
TATCCGCTCACAATTCCACACAACATACGAGCCGGAAGCATAAAGTGTAAAGCCTGGGGTGCCTAATGAG
TGAGCTAACTCACATTAATTGCGTTGCGCTC
>araB_Cas5_498929_499029
CGGGAAAAAATATGAATATATTCCGGCGCTTAATGCCACGCCGGAACATATCGAAATGATGGCTAATCTT
GTTGCCGCGTATCGCTAAAGCTGAGCGGTAA
>araB_Cas5_576636_576736
GAAGCCAAGACGGGCATAAGTAGTATCACCATCATCTGCATCATTAGAGGAGAAGTAGTGCTTAGCATTA
ACTTTCCCGTACAGATCCAGCTTGTTACTGT
>araB_Cas5_593809_593909
GAAACAGCATATCGCTGACCATTTTCGCCATTCGCGTCAGCTCTTCGAGATTAGAGTAGAGCACATCTTC

CAGCTCCTTCTGGCTGCGCGACTGGCTGAGG
>araB_Cas5_653954_654054
CAGATATTACTGGCGGGAAATCTGGCGCAGGCCCGAATGATGATCGAGCGTTTTAAGCCGGGGCTAATCT
TGCTCGATAACTATCTTCCTGACGGTAGAGG
>araB_Cas5_657200_657300
AGTAAAGTTACACTGGACAAAGCGTACCACAATTGGTGTACTGGTAACCGACACAGCATTTGTGTCTATT
TTTCATGTAAAGGTAATTTTGATGTCTAAGA
>araB_Cas5_715082_715182
CAGGCACGGAAGACGCATATAAGATCTACTGCGAAAGCTTCCTCGGTGAAGAACATCGCAAGCAGATTGA
GAAAGAAGCGGTTGAGATTGTTAGCGAAGTT
>araB_Cas5_941748_941853
ATGCCCGCATGATCATCATCGATCCGCGCTATACCGACACCGGTGCCGGGCGCGAAGATGAGTGGATCCC
TATTCGTCCGGGAACAGATGCCGCACTGGTTAACGG
>araB_Cas5_972208_972319
AAGTGCTATGGACCACGCTCAATCAGTTCAATAACACCGGTCTGACGATGCAGGATAAGCGGATCCTGTT
TCGCACTCGCGAATGTCTCCCGGCATTATTTGAAGGCTTTAA
>araB_Cas5_1003470_1003570
ATTGGGATTAACGTGGGGCTGTGAGCTATTTGCCCATGATGGCACGGTCAACATTAGCGGATCGTTTCGC
CGTAATACATGCGTGCTGGCACAGGATAGCA
>araB_Cas5_1044452_1044552
TCAGCAGTGAGCTTACGTCCGGCATGTCCTTCAAGAGATACGCCATGATTAGCGGCAACATCTGCCGCTG
TCGCATCTGCAGGGTGTTTTACCAGACCATG
>araB_Cas5_1046810_1046910
TAGTCACCATCGCCATATGCTCGCGCTTTTGCGTCACGCTAATCTTCTCAACGCCTTGCGGCAGGTTATT
CACCAGAATCCGGTTGGCACGATCGACCGCT
>araB_Cas5_1067053_1067153
TCACCGACGTGATGAGCTTTCGCTTCTTGAGTTGGCATGCGTATCCTCCTGTTGAAGATTAGCCGTTAAG
TTTAACTGCCAGACCTGCGACATATTCCCCT
>araB_Cas5_1203174_1203274
GTGCAATTACGAAATCTGACTTAAGACCGGATATCTATCCGAAAGATTAGCAGAACACTTTCAATTTTTA
ACCACAGAACGATGAGGCTAATCGTGGGTAA
>araB_Cas5_1262015_1262115
CTTGCCCGGCTTAAAGCATGGCTCTGTGCAATGGGGAAAGATTAGCGCCTTTCGCCACAAAGCCATTGAG
CCATTCCGGGGCTTGCTCTAGCACCTGGCGG
>araB_Cas5_1269289_1269389
GTAATGACAACATTTTGCGGCTATTCTTGAATTGTTCTGGTTCAAGATTAGCCCCCGTTCTGTTGTCAGG
TTGTACCTCTCAACGTGCGGGGGTTTCTCT
>araB_Cas5_1441734_1441834
TTTCTTCATTGTGGTTCTCAATTACAGTTTCTGACTCAGGACTATTTTAAGAATAGAGGATGAAAGGTCA
TTGGGGATTATCTGAATCAGCTCCCCTGGAA
>araB_Cas5_1596825_1596925
CTCCGTTGCTGACAACAAGCCCCTGATTACTGATATTCATTTCGCCGTGGCCGTCATAACCAAGATTAGT
TCCGAGATTAGTGATAACGGAGTTCTTATCC
>araB_Cas5_1655257_1655357
CGCTGTGAATGGTTGACAAAAGATGAAATAGAATACCTTTTGTCAGCTGACACTTCCTCTTATCTTATTG
ATAAAATGGATTTATGTTCCTACGTGCGCCC
>araB_Cas5_1732013_1732113
AGTAAAACGAATCTTCCCAGCGATGATTAGCGGACAGGAGCGTCTTGCCTGTATGGATGATGCCGCCAGG
CTGCGTGATGCCCTCGGCGTACGACTACCAG
>araB_Cas5_1786969_1787069
CCATTCGGAACGGAAACACCCATTCCGGAAAGATTAGTAATCATTTCACCCAGGGAGGCATTTTTGCCCC
CAACCCTGTCTACATCATTCATGCCGAGTTG
>araB_Cas5_1806095_1806209
CGTACAACAAAAAAAGAGACCATCGCGGTCCCGGAAACTTTCTTAAGGATCAAAGATTAGCGTCCCTGGA
AAGGTAACGAATTATAAAAAGGCGCGAATAACTTAGCAATGTATT
>araB_Cas5_1877523_1877623
AATTTTCATTTATTGATCTCACATATTTATCCAAGATTAGAGTATCGCGGTATCGTTTTGTTTTGCAGCA

CTATTTTTATTACATTCACTCAAAACATATT
>araB_Cas5_1937486_1937586
ATAAGCAGTTACTTAATTTAAGTGACGATCGCTAAAAACGACTGTCACTGTCCTAATCTTATACGACATC
CGAATGAGATTAATTTATCGCCATCGCGGCG
>araB_Cas5_1942049_1942149
CTTCAGGTTGGCGTCAAGTTTGGCTCGACTTTGCGGCATAAGTTCCACTAATTTTCCATGGATTGCAACC
GCTGTAGCCCGCGCTATCTCTGGGGAAAGCC
>araB_Cas5_1992100_1992200
AACTGAGCTATTCCCGCATTCATCAAGCAATCAGTTAATCACTTGATTTTATTATCGTCTGGCAATCAGT
GCCGCCGTTCGATGCGTTGCATTCTACTTAC
>araB_Cas5_2010298_2010398
ATGACCGCCGCCCACTGGTACTGTCGCCAGAAGCTGCGCGGGAATGGATGCGGCAAGAGATTAGCGGTAA
AGAAGCCTCAGAAATCGCGGCCAGTGGCTGT
>araB_Cas5_2237720_2237820
AGTAAGAAAACTTTTCTTATTTAACGCACTCATGGGAAGCCCCTAATCTTAAAGGTGCAAAGACGCAAGA
CGCAGAATTTCGTTTTGCGTTGTTGTTTTTG
>araB_Cas5_2315653_2315771
CCCGATGAAAGATTAATTAGTCAAGATTATGATATCTTTTTAACGGATAATCCGTCTAATCTTACTGCCT
CTGGCTTGCTTTTAAGCGATGATGAGTCTGGCGTACGGGAAATTGGGCC
>araB_Cas5_2324759_2324859
TTGGTCGCATATCGCGTTTTATGACGCGTTTTGTCAGCCGGTGGCTTCCCGATCCACTGATCTTTGCCAT
GTTGCTGACATTGCTAACATTCGTGATCGCG
>araB_Cas5_2491567_2491679
TTTGCCGCATTCATTTGGTCCAGCTCTTCATAATCACCTTGCTGTAGGTCGACGATCGCGCGGTCGCTGG
AGCCGCTAATCATGATCATCGGAAAACCATTTACCGTTGCGTT
>araB_Cas5_2520907_2521007
TATAATGCGACTCCACACAGCGGGGGTGATTAGCTCAGCTGGGAGAGCACCTCCCTTACAAGGAGGGGGT
CGGCGGTTCGATCCCGTCATCACCCACCAAC
>araB_Cas5_2531204_2531304
ATTAATATCAGACGCAAATCCTGCATCATTATATTCTCTGTTGTTCTAACACCTTGCCACCACGGCAAAC
ATTTACTCACTAAGAGTATTTGCCGATTACC
>araB_Cas5_2544477_2544577
AGTTCGTCCGCCAGCTTTTCGATCTCAGGGGAGATATCCAGCAAGATTAGGTTCGCGCCATGACGTGCAA
AAGTTCTGGCAATTCCTTCGCCAATTCCCTG
>araB_Cas5_2615966_2616066
GATTCAGGAAACTGAAAAGTCATTTGAGTGGGCTAATCTTCGCCGTTACACTCAAAGGCGGCGCGGTGGG
AACGATATTTCACAGTATCGGTCAAATGACT
>araB_Cas5_2649102_2649210
AAACCACTGGCGGTCAGGGCAACGTTCAGTTTTTGCGGTTTATCGGTAAGATTAGTGATATCCAGCGTCA
GACGCGAGGTATCGCCACTCGCCATAAAGCGCGGCATGT
>araB_Cas5_2818910_2819010
GGAGAAATTTTGAGGGTGCGTCTCACCGATAAAGATGAGACGCGGAAAGATTAGTAACTGGACTGCTGGG
ATTTTTCAGCCTGGATACGCTGGTAGATCTC
>araB_Cas5_2884239_2884339
AATAATCACATTCACTGCAAAAATATATTCATTGGTTTAATACAATTAACCTATACATATATTAAGATGT
GTTGAATTGTTTAAAGACAATAATGCATGCA
>araB_Cas5_2903977_2904077
ATAAAAACTTGAGAAAGAGATAACGGGTTATATGGTGGTTTATCCCCGCTGGCGCGGGGAACTCGACAGA
ACGGCCTCAGTAGTCTCGTCAGGCTCCGGTT
>araB_Cas5_2925655_2925764
GCGTTTTAATAACTGGGATGAGGTGCGCCAGACGCTGGAGCGCGACTTAAGCACTTGCGCTCAGGGTAAG
ATTAGCGTGGCGTTATATCGTCTTGATGAACTGGAAGGCC
>araB_Cas5_2942699_2942799
CCTGAGCCGGAACGAAAAGTTTTATCGGAATGCGTGTTCTGGTGAACTTTTGGCTTACGGTTGTGATGTT
GTGTTGTTGTGTTTGCAATTGGTCTGCGATT
>araB_Cas5_2992772_2992872
TAGATAATATCCTTCAAACGGTACATAAACCAGAACGATCTGTAAGATTAGCTAAGCAGGACCAAGGATA

TAAGAATCATTATTTATCAGATGAAATGTTA
>araB_Cas5_2998926_2999073
TTGCGTAAACGCTTTTTATTTACAACAAAATGGGGAAGTATTACGGCGAAGATAAATTGGAGCGGGCGAA
GGGAATCGAACCCTCGTATAGAGCTTGGGAAGCTCTCGTTCTACCATTGAACTACGCCCGCTTCGAGATG
CGTAAGGC
>araB_Cas5_3056044_3056144
GTGGCGCTCCGCGGTTGGTGAGCATGCTCGGTCCGTCCGAGAAGCCTTAAAACTGCGACGACACATTCAC
CTTGAACCAAGGGTTCAAGGGTTACAGCCTG
>araB_Cas5_3104244_3104344
TGAAAAGAAACTCAACATGATGAATGCCGAGCACCGCAAGCTGCTTGAGCAGGAGATGGTCAACTTCCTG
TTCGAGGGTAAAGAGGTGCATATCGAGGGCT
>araB_Cas5_3170477_3170577
TGAGAAATGAAATTTACCCAACGTCTTAGTCTGCGCGTCAGGCTGACGCTAATCTTTTTAATTCTGGCCT
CGGTGACCTGGCTGCTTTCCAGCTTTGTCGC
>araB_Cas5_3184702_3184802
CGCCCTGAGAATAAGCGGATTCACTATAACGCTAATGATTAGCGGCAGCAACGCATAGCTTCACATAATT
CTGGTTTATGACTTACCCTTATCGCACTACA
>araB_Cas5_3185950_3186050
ATTGCTTACGAGTAATCGCATTTGGCCGGGTTCGCCCGGCCACATCATTAACGGATTAATGATAAGTGGA
TCAGATGTATAAAAAATTAAAATTAACCACA
>araB_Cas5_3223455_3223555
GCTTTGCTTTTACTGTCGAACAGCCGCAGCAATGGTCAGCAGAATCCCCTTATCTTTACCATCTGGTCAT
GACGCTGAAAGACGCCAACGGCAACGTTCTG
>araB_Cas5_3318098_3318198
TGAACCCTATAACCGCAACTGCGGTCTGGAGCACTTTCCAGAAGGATTTTTTCAAATCCCACTACGAAGG
CCGAAGTCTTCACAGTATATTTGAAAAAGGA
>araB_Cas5_3318312_3318412
CGTTTTGGGTAAAAAATGCAAATACTGCTGGGATTTGGTGTACCGAGACGGGACGTAAAATCTGCAGGCA
TTATAGTGATCCACGCCACATTTTGTCAACG
>araB_Cas5_3322069_3322169
TTTGGTACCGAGGACGGGACTTGAACCCGTAAGCCCTATTGGGCACTACCACCTCAAGGTAGCGTGTCTA
CCAATTCCACCACCTCGGCACGGATACTACT
>araB_Cas5_3350521_3350632
CACACATTTAACTGATTCATGTAACAAATCATTTAAGTTTTGCTATCTTAACTGCGTGCGGCCTGAAAAA
CAGTGCTGTGCCCTTGTAACTCATCATAATAATTTACGGCGC
>araB_Cas5_3377492_3377602
CAATAAGGAACGACTCTCTTTGTTCGAAAATCAAACAAAAAATGAGCAATACCCGACATTTGGGCAGAAA
ATTGGATGATAGTTTACCAGATTTTGCGACCATTGTGGTGA
>araB_Cas5_3519221_3519321
CCTCTGCATATCTGGTCGCTGCAAGCGCGCTGCCTTGCTACCACCGCTCTGGCGATAAATCACCGGGTAA
GATTAGCGTAAAAAAGACAGCAAAATGCCGC
>araB_Cas5_3563615_3563715
TTATGCAATATATTGAATTTGCATGGTCTTGTAGGCCAGATAAGACGTTCACGTCGCATCCGGCATGAAC
AAAGCGCACTTTGTCAAAAATCTAACCTACT
>araB_Cas5_3600798_3600898
CGATATCTTCTGGCGCTTCAGTGGTAGCAACAACTGTGCCAGAGCTTAAGAGCAACGAGGTTATCATTCA
CTGTTTTATCAGACCGTGATTTTATCCACAA
>araB_Cas5_3600933_3601033
AATGTGACATAGAGATGAAATACCGGGAAGAGACAACGGGGTCTCTTTCCCTGCTACGGAACCCATTGCA
GGGAAAGAGTATAACACGCTTTTATTATTCA
>araB_Cas5_3700098_3700198
TGTTGTAATGACAACGTTTCGCGGCTATTCTTGAGTGGTCTAGAGTCAAGATTAGCCCCCGTGGTGTTGT
CAGGTGCATACCTGCAACGTGCGGGGGTTTT
>araB_Cas5_3737213_3737317
TATATTTTTCACAAATTTGAGAGTTGAATCTCAAATCATATCAAAAATAGCTGTCAAGAGCACCCCAAGG
AATAGTCCAAATCTGAAACTATGTCACGTGTTAAC
>araB_Cas5_3811754_3811865

CAGTTCTATGTGAACTCTCGATTGCCAGGCCCAAATGCCAAACCCGAGATTCTCAAAGGTGGCGTAGTAT
ACGCTGACTCAGCGATGTGCTCAAGTCCCGAACAGACAAAGA
>araB_Cas5_3850577_3850677
ACCTGCACACCGCGCCCCGGCAGCCCCGCCAGCGCCTCGCGCCCCGAACCAAACTCGGCAACTACCTGCA
AATCAGGTTCCAGCCCCAGCAGCTGCGCAAA
>araB_Cas5_3859785_3859885
CCTCCAGAAAATCGGGGATGGTGGCAATGCCGCGCTTCAGATAGCGCGGTAAAAAGATTAGCGCGAGGAA
GATCAGCGTCACCGCTGAAGTCACTTCCCAG
>araB_Cas5_3885842_3885942
GCAACAGTATTTCGCGACGGCGTGGATCCCGCATAACGACGGTACCAACAACTTCTATACCGCTAATCTG
GGTAACGGCATCGCCGCTATCGGCTATAAAT
>araB_Cas5_3888568_3888698
TTTGCTATATCTTAATTTTGCCTTTTGCAAAGGTCATCTCTCGTTTATTTACTTGTTTTAGTAAATGATG
GTGCTTGCATATATATCTGGCGAATTAATCGGTATAGCAGATGTAATATTCACAGGGATCA
>araB_Cas5_3931432_3931541
TTTGGTTTTGGCGTTGAACGCGATGCCGTGTTTGGCTTTTTATCGCTGATCTTCTGGCTGCTAATCTTTG
TGGTTTCCATTAAATATCTCACCTTCGTGATGCGGGCAGA
>araB_Cas5_3950293_3950393
CTTCGAACAAGATGCAAGAAAAGACAAAATGACAGCCCTTCTACGAGTGATTAGCCTGGTCGTGATTAGC
GTGGTGGTGATTATTATCCCACCGTGCGGGG
>araB_Cas5_3966135_3966235
GGCGTGAGTCATGCTAACTTAGTGTTGACTTCGTATTAAACATACCTTATTAAGTTTGAATCTTGTAATT
TCCAACGCTTCCCGTTTTATCTTAAATGCGA
>araB_Cas5_3982523_3982629
GTTGGTAGAGCCCTGGATTGTGATTCCAGTTGTCGTGGGTTCGAATCCCATTAGCCACCCCATTATTAGA
AGTTGTGACAATGCGAAGGTGGCGGAATTGGTAGACG
>araB_Cas5_4049868_4049982
AACTGAACAAAAAAGAGTAAAGTTAGTCGCGTAGGGTACAGAGGTAAGATGTTCTATCTTTCAGACCTTT
TACTTCACGTAATCGGATTTGGCTGAATATTTTAGCCGCCCCAGT
>araB_Cas5_4096005_4096105
TGTAGAGTGAATCTGCGCCACATAGTGGGCAATTTCACTGTCAGGATTAGGGGTAAAGGTGGTCAGATTC
GCGGTGGTGCTGACGACCTGACGGAAATCAT
>araB_Cas5_4117619_4117719
AGCGGTAATCACCATCTCAACTGCGAAAGCCTGCACAAAATTGATATGAGGATTAGGGTAAGTAGAGAAA
GTGCCAGCCAGATCAACACTTTCAACGCTGC
>araB_Cas5_4150400_4150500
CAGAAAACCCTCGCGCAAAAGCACGAGGGTTTGCAGAAGAGGAAGATTAGCCGGTATTACGCATACCTGC
CGCAATCCCGGCAATAGTGACCATTAACGCT
>araB_Cas5_4179584_4179684
GCCTTTACGTGGGCGGTGATTTTGTCTACAATCTTACCCCCACGTATAATGCTTAATGCAGACGTATATC
CGAGATATTCGGGTTGTGGCAAGGCGGCAAC
>araB_Cas5_4274019_4274119
TTTGTTTCCCGGAACCGAGGTCACAACATAGTAAAAGCGCTATTGGTAATGGTACAATCGCGCGTTTACA
CTTATTCAGAACGATTTTTTTCAGGAGACAC
>araB_Cas5_4279782_4279882
CGCCGTGGCGCTGGTGACCTTCTATGCGGGTGGCTGGGCTATCTAATCTTTCATCGGATTCTGAAAACGG
GTGGCAATGGCTGCCCGTTTTTATTTTCTCC
>araB_Cas5_4303043_4303152
ACCACCGGTAATGGCAGGGAGTTGAGCGCGCTCATTATTGCGGCTCAAGATTAGCGACTGCCGAAGCGCC
GATGCGGAACATTTCCGGGTCAGGTTTATCGACCATGATT
>araB_Cas5_4323158_4323275
GGCGAGTACAACGGCCCAGCTCAGAAGCGAGAACCAGCTGCGCTTGGGTGGGGCGATGGTGATGGTTTGC
ATGTTTGGCTCCGGTCTGTAGGCCGGATAAGGTGCTTGCACCGCATCC
>araB_Cas5_4413822_4413922
GGGCTAATGCCGCTGCTTTGGCGCTTCTTCCCTGGTCATCCTAATCTTCTTGCGTCCTGGTTCGATGGCG
AGAAACCGCAGATTGCCGCTGGCGAAAGCTA
>araB_Cas5_4432983_4433083

CGATCAACGGCCCGACAAAAATAAGCCCCCACATCAACCCTGCTAACAGGGCGTACAGCACGCCGCTAAT
CATTACTGGCATCCATTGATCTGTCAGAAGA
>araB_Cas5_4467317_4467425
TCTGGTTTATCGTTGGTTTAGTTGTCAGCAGGTATTATATCGCCATAGATGCTACGAATATTATTGGATT
CTCCTTATTATTTGCGGCGCTTTTTTCACTTACCGGAGG
>araB_Cas5_4470473_4470573
TGCGTTTGTCATCAGACTTAATCCGGGCATGATAGCCCGGATTTCCATCAAGATTAGCGACGAACAGCGA
TCGCTTCGATCTCAATCTTCACGTCTTTCGG
>araB_Cas5_4518333_4518433
ACATGTTGTTTTTCTAAGTGTTATAAGGTAGGTATAAAATGGGATGGAGCCTCTGCTTCTGGCATGTGTC
GGTCAGAATGACTCATGATGTGGTCTGCTAT
>araB_Cas5_4540640_4540740
TTGATTTTACTTGTTACAGAACATATCACATGATATATAGATAAGATTAGTTGCATTAATGATGAGGGTT
ATTATTAGATTCGTATCCGATTGATAAATAT
>araB_Cas5_4572271_4572371
ATCTAAAATTGATTTATAGTATCGACCTGAAAAAATAGTTGTTGCCGCCTGAGTAACTATACAATATTCT
GAAAGGTTTTCTTTCAAATTAGAAATGTTGT
>araB_Cas5_4591286_4591386
CCTGGCATGTATTGATTAATAGTTGGCCGAAGCCGTTCTAGGTTTGCGTTGCGTTTGAGGAGGTAAATTG
ACCGCTATCCTGTAGTGATGTTGCCAGAGTT
>araB_Cas5_4619509_4619609
CCGGAAAGCACACCAACTGTCTATCGCCGTATCAGCGAATAACGGTATACTGATCTGATCATTTAAATTT
GAAGCACTGAGTACGGAGAACATATGAAACG

Regions used to identify motifs shown in Figure 2B

>Cse1_wt_endogenous_19061_19173
CGCAGTGGTAGAAGGCGAGCCCATTCATCTTCGCTGCTTCGAATCCACCCACGAAATGCTGCTGGAGCAA
TTAAGTCAGCATAAACTGGATATGATCATTTCTGACTGTCCGA
>Cse1_wt_endogenous_28749_28849
CTGCCAATTTTAGCGTTGGCGTTAACGTCATGCTTAAGCTGCTGGAGAAAGCAGCCAAAGTGATGGGTGA
CTACACCGATATCGAAATTATTGAAGCACAT
>Cse1_wt_endogenous_39745_39845
TCGTCAAAGGTGATTTCACAGCAGCTATCCATACGCAGACCGAGCTTTTCAAGTTTGGTCACTTTGATGC
CCGGTTTGCTCATATCAACAAACCATTCGGT
>Cse1_wt_endogenous_44116_44216
TTTAGCGCGTTGATCCACTCTGGCAGGGCTGCATTTTGGCCCTGCCGCTGACAGGGAGCTCTTATGTCCG
AAGATATCTTTGACGCCATCATCGTCGGTGC
>Cse1_wt_endogenous_94156_94256
AAACCGCCGCGCGTCTGCAACCGGTTTGCGGACGTATGGAAGTGTTCACTGCGCCAGGCAAACCGACGGT
GGTGGTGGATTACGCGCATACGCCGGATGCA
>Cse1_wt_endogenous_157898_157998
CTTCAGCTCGCAAGGCCAACAGGTCATAAGCCGCACGGAACTTAGGATGCTCCAGCAGTTTCCATGCGCG
TTTACCCTGACGACGGGACATACGCAACTGC
>Cse1_wt_endogenous_165061_165161
CAATCTTGAGCCAGACATGACCATCAGCAAGAACGAGATGGTGAAGCTGCTGGAGGCGACCCAGTATCGT
CAGGTGTCGAAAATGACCCGTCCTGGCGAAT
>Cse1_wt_endogenous_186859_186959
GTCAGCCACAGTCAGGCATACCAGATAGCGCAGACGATTTTCCGTTTGCACTTCTTCGGCAAACTGCTTG
ATGACTTCCGGGTCCTGAATATCGCGGCGTT
>Cse1_wt_endogenous_198681_198781
AAGGTATTTACGTCACGGTGAACATCACCGAAGGCGATCAGTACAAGCTTTCTGGCGTTGAAGTGAGCGG
CAACCTTGCCGGGCACTCCGCTGAAATTGAG
>Cse1_wt_endogenous_216423_216523
ACTGCGCTTTCGCCAGGCCGCTGACGCACTGTTGGCGAAAATCCTCGGAGAACGGATTCAGTGAGCACAT
CTGGCGAGTCACCATAAAGGGAATAAATTCG

>Cse1_wt_endogenous_222434_222534
TGGTCAGTTCCTGGCCATCGACCAGCACGCTACCCTCGGTTGGGCGCTCCAGCAGGTTTACACAACGTAT
AAGCGTACTCTTACCCGCGCCTGAGGCACCG
>Cse1_wt_endogenous_269908_270008
CATCAGCCGTCACTAATGCCAACCCCTGCTGTAATTCGTTAAATCCCCGCTGTATGTGCTCAAGCAGTAT
TTCACCTTCTTTCGTCAGCGTAATTTCTCGC
>Cse1_wt_endogenous_290138_290238
GAGTCTTTTTCGAAGATGAGCGCGATGTTTTTACCGGTAAGCTTCTGTACTTCCTTGCCATTTTTTTAT
CGGCTTTGAGCTGTGCGGCAAGGGTCAGCAG
>Cse1_wt_endogenous_298702_298802
TTTGTCAAAAAGCGGGTATGACATTGTTCCTCCTTAAACCACATCCGGCAGCTTATCGAGCAGCTTATCC
AGAGTGATGGGATAATCCCGTACCCGAATAC
>Cse1_wt_endogenous_303272_303372
GCCGGGCATGTCTTTAACTGGGTTGGTGTTACGCACATTATCTTTTCGATAGTGTTTGCTGTCGGTTATT
GTGTGGTCGCTGAAGTATTTCCAAAAATTAA
>Cse1_wt_endogenous_309502_309602
CAAGGTCCGTAGGCGAGGATCCGCAGCGTCGCATTTCCTGTATTTGTCAGGGAGCCGTTGGCGTACTGAA
AGTGGTAGTTCGCCTGACGAGGGGCGACGAC
>Cse1_wt_endogenous_413353_413453
GACCAGAGTTTGCGCCTGACGGCGCTGGTTTTCCAGATTCTGCTTGAGCTGTTCCAGCTGCGTTAGTGTT
TGTTCATCCATTAGCGCCGCAAGGAACGCCT
>Cse1_wt_endogenous_419967_420067
CCTGGTCTATTTCGCTATCGTTATTCTGGTTTCGCTCTATCCGGGCAAGCTGCTGGATACCGTGGGCAAC
TTCCTTGCGCCGCTGAAAATTATCGCGCTGG
>Cse1_wt_endogenous_423034_423134
CGTTCTATGGCGGCGATCTGGACGGGATAAGCGAAAAACTGCCGTATCTGAAAAAGCTTGGCGTGACAGC
GCTGTATCTCAATCCGGTGTTTAAAGCTCCC
>Cse1_wt_endogenous_436658_436758
TTGCTGTCGGATTCGGAAGTGGATTAAGCCCGATCGTTCCTGGGACGATGGGCTCGCTGGCAGCGATTCC
GTTCTGGTATCTGATGACCTTTTTGCCCTGG
>Cse1_wt_endogenous_483943_484043
TATCCTCCTGCGTCATGGTGCCATCGGTGTTGATAACGCCGACAACCATCAGGAAGCTGCTGGATGATTT
CTCAACGCTCACCCCTTGCTGCTGAACTTCT
>Cse1_wt_endogenous_491474_491574
CCAAAACCCGGCATTCTTTTCCGCGATGTCACCAGCTTACTGGAAGACCCGAAAGCTTACGCTCTCAGCA
TCGACTTGCTGGTTGAGCGTTACAAAAATGC
>Cse1_wt_endogenous_557451_557551
ACCAGAATGCCATCGATGGTGTTGTCGGCATTCAGCGTATCGATAAGCTCCAGCAGCTCCGCTTCGCTGG
TGGTTTCCGGGAGGTCATAAGAGCGGGAGAC
>Cse1_wt_endogenous_607646_607746
TAAATAAACAGCTATTTTGTTGAGGAAGGGTAAGATAACGGCGGGTGCCTGAAGCTTTCCGGTTTCAGGT
TTACTCTGAGGTCTGGAAAGATGAAGCCCCA
>Cse1_wt_endogenous_629880_629980
ACGGATAACAACTATGAACAAATCAGGGAAATACCTCGTCTGGACAGTGCTCTCTGTAATGGGAGCATTT
GCTCTGGGATACATTGCTTTAAATCGTGGGG
>Cse1_wt_endogenous_650779_650879
TCATTTTTCACCTGTTTCTCATGCGGGGGTCTTTTGACGAGCTGCCGCGTCCTGGCGGGAGTGCTCAAGC
AGGTTCTGCAAATAATGCAGCGTGACTGCAG
>Cse1_wt_endogenous_659509_659619
ACCATCAGACCAGACTTGGTCGGGATTTCCGGATGCGCTTCTTTAAAGCGTTCCAGCAGCTTCAGCGACC
AGTTGTAATCTGCACCAGGCCGTACCTGACGGTAAATACGC
>Cse1_wt_endogenous_660286_660386
TATTTTTGAATTAACGACTGGCAGTATATCATTGAAACGGACCTGAAAGCAGCCAAAGCGGTCGCGAAAA
TGTAAAATTGTTGTTGGATTGTGCCATTTTA
>Cse1_wt_endogenous_660718_660818
TTCCGGTAACAGAAGTACATGCGCAGCGTTGTCGATAGTCTCTCTGTCATCTTCTAAAGAGAGTAGTGGA
TTTCCCTCGCGGCAAATTAACACCATATTGT

>Cse1_wt_endogenous_667280_667380
GATCGTTCAACAAGGCGGAATAATCTTTGCTGGAGATACCGTCAACAAACAAGTTTGGGTCATAACTAGG
CGTGGAAACCAGCGCCAGCACCCCACCTGTA
>Cse1_wt_endogenous_684840_684940
CGGAAAGCTGCGCCGGAAACTTATTGGCGTGAGCAGAAAGCCCGACACGCTCCAGCAGTTTCAGGGCTTT
TTCACGAGCCGGCGCTTTATCGCGTTTAAGC
>Cse1_wt_endogenous_703368_703468
ATGCTCTTTCGGCAGACCGACATATTCGTCCATGTTGAAGGTGACAACGTGCTTAAAGCTGACCTGGCCT
GCTTTATGCATTTCGACTAACGCTTTATAGG
>Cse1_wt_endogenous_708948_709048
ATGGATGAGTTTTATCAGAACGATAAAACCACCAAAGTTGATTATCTGCACTCCTTTGGGGCGAAATACG
ACTTCAAAAATAACTTCGTACTGGAAGCGGC
>Cse1_wt_endogenous_715065_715165
GTTCGCCGCGCGTCCGTCAGGCACGGAAGACGCATATAAGATCTACTGCGAAAGCTTCCTCGGTGAAGAA
CATCGCAAGCAGATTGAGAAAGAAGCGGTTG
>Cse1_wt_endogenous_765813_765913
CCACGCGCAATTTGTTCACGAATGCGATCGGCAATCTGCCGGTATAAGGGCTTGTGTCCCATTTTTAGTA
TCTCATTAATACGAATTTAACCATTATGCCC
>Cse1_wt_endogenous_791589_791689
CCATCAGCTTGCTTTTGCCGTAAGGGCTTTGCGGTGTGCCGGTCGGGAAGCTTTCAACGTATGGAATTTT
GGGCTGATCGCCATAAACGGTGGCGGAGGAG
>Cse1_wt_endogenous_807613_807713
CCAAAGTCGGTACGCGTCTGCAAAACGCCGTCTGGCATTGCTACCAGACCAGAGCCAAACCCTTGCGGTA
ATACGCGGGTATCAGCGGGTAAGTTAACAAC
>Cse1_wt_endogenous_838687_838787
ACATAAATGATGCCACTCGTACGCCGCGGGTTACGGGTGTCACGCAATGCAGGCTGCTGGAGGGATACAA
CACGAGATCGCCTGCCGGGAGTTTTACCCGA
>Cse1_wt_endogenous_873139_873239
TGGCTATTTCTTTAAAGATATTGAACGTTCGGCCCTGACATTAAAGGACTCACTGTATTTATTAAAAAAT
ACAGAGGAGATTCAACGCGCCGTGATTCTTA
>Cse1_wt_endogenous_906234_906334
TCCGAACAGACTTTGTGGGCGCAGAATAGTAAAGCGCGTTTGTGGATTCGCCTGCGAAAGCATATTGATC
ACTTCTTCGCTGGCTGCTTTGCTGCGGGCAA
>Cse1_wt_endogenous_917062_917162
GCAAAATCTCTCTGATGGGCAGATTCGTCAGGGACTTTCCGCAATGGTACAGCTGCTTGAGCATTATTTC
TCTGAGCAGGGGGCCGGACAGGCGCGATATC
>Cse1_wt_endogenous_919083_919183
TGAAATCAATGGTTATACATGGCGTCGATTTCACCATTGCGTATCTTAACCAAACATCAATAGTGTGATT
ACTAACGTAAATTTTAGGGTTTTGTTGATAT
>Cse1_wt_endogenous_924030_924130
CACAAAATATCGCGGCGACTTTGAAAAACGTTTTAAAGCGTTGCTCAAGCAGCTGGAGCAGGACACTAAC
AGCATCCTGTTTATTGATGAGATCCACACCA
>Cse1_wt_endogenous_927771_927871
AGCTGGCGTCCGCCTTCACCTAACCAACTGTTGAGACCTGCATCCTCGAGCAGCTTTTCCAGGCCAACGC
GACGCAAGATCTCCGACAGAGCCTCATCACT
>Cse1_wt_endogenous_980139_980239
GCGTGCGTCTCAACGTGAACGTGCGTGAAACGCACGCCATGCTACTGGATGTGCTCTCTGAACAGCACGA
GCAGCATCAGGATCTGTTTAACAGCAACCGT
>Cse1_wt_endogenous_1049672_1049772
TTCAGGGGCAGCAAACGCAGCTGGCGTCGTCAACATGAAGGCCATAAGGATTGCAGAAAGCTTGTGTTTC
ATAACTTTTCCTTTATTCATCGCATGGACAA
>Cse1_wt_endogenous_1093946_1094046
GTCGGCGGCGAAGAGTTTGGCGTCTTGCTGACGGACATCGATACTGAACGTGCGAAAGCTTTAGCGGAAA
GGATTCGGGAAAATGTTGAGCGTTTAACTGG
>Cse1_wt_endogenous_1096826_1096926
TGAGGAAGTGATAGGAAGTGACCAGATAATACATATATGTTCTGTACTCTCTTGCGCATTTTGATTGTTG
ACTGAGTAACCAGACAGTTGATGTGCACGAT

>Cse1_wt_endogenous_1099516_1099616
TTTTCCGCCAGAGCGCATTTTGAATGTTTCTCCGCGCCGCTTACTGAACTTCCTTGAATCTCGCGGTATG
GCACCGATTGCGGAATTTGCAGACCTTTAAT
>Cse1_wt_endogenous_1273992_1274092
TATGAATGATAACGGGCTGGATACTGGCGAGCAGGCAAAAGCTTTCGCATTGGGAAAAGTCCGCGACGCG
CTTAGTCAACAGGTTAATCAGCACGTAGAGT
>Cse1_wt_endogenous_1281291_1281404
CTGACGCTGGCAAACTACGGTCTGGAACGTGGCCTGAACGACGTTAACTGTGCAACCAGCTATGACGATG
TGAAAGCTTATACCCCGGCCTGGGCCGAGCAGATTACCGGCGTT
>Cse1_wt_endogenous_1290401_1290501
TGATGATATGTGATATCGCGATGCCACGAATGAACGGGCTTAAACTGCTGGAGCATATACGTAACAGAGG
CGACCAGACCCCAGTTCTGGTGATATCTGCC
>Cse1_wt_endogenous_1328066_1328166
GTTTTGTGTATCCCAGGTAAAGCCCGTTGCCATCACCTGAAACTCAACGCCATGTGGCTCCAGCAGATTG
CGTTCGCCATTAGGCCAGGTGCCTTTAATAA
>Cse1_wt_endogenous_1340783_1340883
GAAACCTGTTCCGTTCGCGTGGCGAAGTTGATCGCGCTATTCGCATCCATCAGACCCTAATGGAAAGCGC
CTCGCTGACCTATGAACAGCGTCTGTTGGCG
>Cse1_wt_endogenous_1428341_1428441
GCCAGCGATTGATGGTCTTGAACAATTGACGGGCCAGTTGATGCTGCTCGAGCAGGTGGCGGAAATTCAT
GATGGTGGTGCGATCCGGCAGGGCGCTATCC
>Cse1_wt_endogenous_1451764_1451864
CACCTGGATGGTATCGCCTGGCTTTACGGGTTCGATAAAACGCAAGCTTTCCAGCCCGTAGTTAGCAATG
ACCGGACCGACACCGGCATCGACAAACAGAC
>Cse1_wt_endogenous_1500369_1500469
CGTAATAACACATTTTTTTAGTGCCATCTTCCTGATAACTAAAGGCTCGCTGTTCTTCACTTAGACGAT
ATTTGCGCATAGCGTTTTCCCACAGGTGACT
>Cse1_wt_endogenous_1529865_1529965
CGGGTTAGATGGTATAAAAGCAGATAATCCAATAAAAGCAAAGCAAGGGAGTTCAGATGCTTCAAATTAT
TAGAGGCAAACTTGTCATTTTTTTAATTACC
>Cse1_wt_endogenous_1579238_1579338
ATTAACAACACTGAAGTTTTATTATTCTTACGCAGCCAAAAGGGCTTCAACAGACAGAGATACTTTGCTA
TCAACATACGAAGCGTAATGGGAATGGTTAT
>Cse1_wt_endogenous_1626888_1626988
TTCTGTATTCAGTACTTTTAATTTTGCTTTATCAGCTTGCGCAAGTTTGGCTCCGGCAAGGACAAAACGT
TGATGAATCACCTCCACCAGGCGGATGGATT
>Cse1_wt_endogenous_1719387_1719487
AGTTGCCCGAGTTGAGTATCAAGCTGTCCAAACTGCGAAAGTGTAAGCTGCTGGCCCTGGCAAATATCCA
GTACAGCCTGTTTCAGAGATACCGGGATCGG
>Cse1_wt_endogenous_1802003_1802103
GTTCGGTACGTGCGGACCGCGGCACATATCGACATATTCTTCATGGAAGTACAGACCTGGCTTGTCATCA
TGGGCGATGTTTTCGTCAAGAATGGAGACTT
>Cse1_wt_endogenous_1821855_1821955
GGCCAAAAAAAACCGGCGCAATGGCCGGTTTCCGTTGTTACTCAAGCTTTCAGACGAATTGATTACTTCG
CAGCCTGTGGATCAGTGTCGTATTCAGCACA
>Cse1_wt_endogenous_1898912_1899012
GTCATGGAGATGATTAACATCGACTTACTGACGGCAATGCTGCTTGAGCCACAACTGCCGCAAATCAGTA
GCGCCAGCCTGACGGTGGACAAACGGCATTT
>Cse1_wt_endogenous_1942013_1942113
CTGCGTTTCGGTTGAGGCTAATTGTGCCTCAAAATCCTTCAGGTTGGCGTCAAGTTTGGCTCGACTTTGC
GGCATAAGTTCCACTAATTTTCCATGGATTG
>Cse1_wt_endogenous_1952341_1952441
TGCAGTACCGCGTTGCCTATGGCGACGGCGGTTTTAGCGAACTGCAAAGCGCTATTCGCATTCATGGTAA
CGCGGTGGAATATATTCCCATCGCGATTGTG
>Cse1_wt_endogenous_1977267_1977367
AAACAGCCTGTACTCTCTGTTCATCCAGCAGTTGTGGGATAATATCGGCAGGATTCTGGGAAAGTTTACG
TCTTTTTACTGCCCGGGATGGCGGTTGACAT

>Cse1_wt_endogenous_1993531_1993640
ACTTCCGGCAATTTCAACACGCTGGCAAGCGCGGTCAGTCGCTGGTGAACGGTAGATTGCTGCGAAAGTT
TGCTGGTTAAGGCCGTCGCCGCATTGGTGCGCGCGAGTTT
>Cse1_wt_endogenous_2030915_2031015
CATGTTGTGCCTCTTGCTGACGCAACGCCACCGCCTGTTTGATTTTTGGCTCAAGCAGTTTTGCCACCGC
GGCAAAGACCGGCACGACCACCGAGTTACCG
>Cse1_wt_endogenous_2050876_2050976
CTTAAAGGCAACGGTTGAGGATGGCAGTGGTAACCTGATCGAAGGTCTCACTGTGTACTTCGCCTTAAAA
AGCGGCTCTGCCACATTAACGTCATTAACAG
>Cse1_wt_endogenous_2183414_2183514
GATCCGCTCCTGCCAGCGCGAAACATCTATTCCGTGAATGGTGTAACTGGCAGGAATGCGAATAGCGAAA
GATTTAACCGGGCGATAACCGTAGAAGTGGA
>Cse1_wt_endogenous_2336117_2336217
CTGATTTCCGTTTTGCTTAACTGGCTGTCGCTGGTAGCGGCAAACAACAATGGCTCCAGCAGTTTGCTTA
AACCACTGCGCTGGATCAGCACCATATAATG
>Cse1_wt_endogenous_2339692_2339792
CTGCGTCTGGGCTATATTGCCGAGCGTGCTGGCGGTTTATTTGGCAAAAAGGTGCTCGATGTCGGTTGTG
GCGGCGGCATTCTGGCCGAGAGTATGGCGCG
>Cse1_wt_endogenous_2354451_2354551
GACTGGAATCTTTGCGTCAGCAGGCACCAGCCCATCCTTACTCCCTTCTCGAGCCACAACGCGTGCTCGA
TCTCGCTTGCCAGGCGCAGGCATTAATCGCT
>Cse1_wt_endogenous_2367078_2367178
AATCCACCCTGTTAAGAAAGTCTCGGTGGTTATTCCCGTTTATAACGAGCAGGAAAGCTTACCGGAATTA
ATCAGGCGCACCACCACAGCCTGTGAATCGT
>Cse1_wt_endogenous_2406861_2406961
TACTCACTCTTTTCTATTTATGACATGCGCGTGTTTGTATAAATGTAAATGTGAGTCCTTGTTCCACTCT
CGTGCAGCATCGCTGGTCATACGCGAACACG
>Cse1_wt_endogenous_2416250_2416350
AACAGGATGAAGTTGATGGTCTGGTTTCCGGTGCTGTTCACACTACCGCAAACACCATCCGTCCGCCGCT
GCAGCTGATCAAAACTGCACCGGGCAGCTCC
>Cse1_wt_endogenous_2425796_2425896
CGAGAGCTTACCGCCAGCGCCAATTAAACCGATGATTACAGCGAGCACTACAGAGCTGATAGCCAGCTCC
AGCGTGACGAGCGCACCCTGTAAAATAACAC
>Cse1_wt_endogenous_2488203_2488303
CTTAATAACGGCGCGACGGGTACACCAACTTCATGTATTCTGGCTAACCAAACTTCAGCTGCCTGCGTTT
TTAACGTCCGCTCAATATATTGTTTAAGAAT
>Cse1_wt_endogenous_2531202_2531302
TGATTAATATCAGACGCAAATCCTGCATCATTATATTCTCTGTTGTTCTAACACCTTGCCACCACGGCAA
ACATTTACTCACTAAGAGTATTTGCCGATTA
>Cse1_wt_endogenous_2535666_2535766
GAAGATTATCCGTAACACGAACTTCGAAGATGCGAAGGTGTTAGCAGAGCAGGCTCTTGCTCAACCGACA
ACGGACGAGTTAATGACGCTGGTTAACAAGT
>Cse1_wt_endogenous_2560797_2560897
ACTTCAAACTTGTTGAGCGTACTGAAGGCTATATACCAGGATCGGCAGAAAGGGAGTGCAACGAATACCA
GTTATTATTCAAGCCGTACAACATGAAATAA
>Cse1_wt_endogenous_2601858_2601958
TCTGTTTGCTAGTCTGGCTTCGCTGTTTTTACTCGGTCTGGACAGACTGGCTATTAAGCTTTCCGGCATC
ACATCGCTGGTGGGCGGCGTGATTGGCATCA
>Cse1_wt_endogenous_2622891_2622991
CCAGCTCTGCCGCTGGCGTTTTTCAATTCACCTGTAAATCGCAAGCTCCAGCAGTTTTTTTCCCCCTTTT
CTGGCATAGTTGGACATCTGCCAATATTGCT
>Cse1_wt_endogenous_2639791_2639891
CCAGACCACGCACCAGACGCTGGTTTACGGTGTAAGCGATCCCCGCGCTCTCCAGCAGTTTGCACAGACC
GGCAAAATGCTCACGAGATTCCTCGTCCAGA
>Cse1_wt_endogenous_2709255_2709355
ACCTCGGGAGTATCACCCCGCATTGAGTCACGGATTAAGTGATAGCTTTCCCAGGTTTTCTGCATTTCTG
GGTTATGAGCCAGTTCGTTAAGCAGCTCACT

>Cse1_wt_endogenous_2720942_2721042
CCAGAACATGTGGCAATATCTAACCCGCTCGATCTACGCGATGACGCCAGCAGTGAGCACTATATTAAAA
CGCTGGATATTCTGCTCCACAGCCAGGATTT
>Cse1_wt_endogenous_2740707_2740807
GGATCTAACGCTTCCGTCGCCAGTGGCAGTCCCATATTCACCAGCTCAAGCAGCAATTTACGCGCGATCT
GCAGCCCGGCTTCTACATCAAAAGAGCCATC
>Cse1_wt_endogenous_2752485_2752585
TGGCATTAATGGCCGACGGTGAAGACGCAAACCTGCAAAGTCAGCTTTACACGGCTAAACAACTGGTGAG
CGAATTGATTGGCATGGACAGCAAACTGTCC
>Cse1_wt_endogenous_2796975_2797082
AGCACGCCGAGCGGCGGCAGAATGATGGTGATGACGATTCTCCAGAAACCCATATGTACTCCCTATAAGA
AAATTACTCATTGTTTAAAAAGAGATTTTATCTCTTAA
>Cse1_wt_endogenous_2800933_2801033
GTCTGCATCCAGCGCCACACGCGGCCAGTGCATGAGCCAGCTCGTCTACTTCTCCAGCAGCTCCGAAAAC
ACGCAGCGTTTTATCGAACGTTTAGGTCTGC
>Cse1_wt_endogenous_2810840_2810940
TCAGGAGATCCTTCTGACTCGTCTTTGCATGCACATGCAAAGCAAGCTGCTGGAGAACCGCAATAAAATG
CTGAAGGCTCAGGGGATTAACGAGACGTTGT
>Cse1_wt_endogenous_2910936_2911036
GTACAAGGAACACACAGACACATACCTGTTTTGTTGTTGTACATGAAAGGACTCAGGACAACAGCTGGAC
GATGTCCAGCTTGCTCGCTACCTTTTGTCGG
>Cse1_wt_endogenous_2928545_2928645
CTTCGCTATCTACCCGATCCTGCTGGTTTATAGCGTGGCAATCACCAATACCGTTGAAAGCTTCATGTCT
CACCAGCTGGGTATGACGCCACCGCCGCGTG
>Cse1_wt_endogenous_2932087_2932187
TTCACGCCCATAACGCGCGCGATATCGCGGTACTTCTCACCGGTAAAGTCAGCGTTATAACGCATGACAT
GCGGTAACAGGATGGCGTTCGCAACACCGTG
>Cse1_wt_endogenous_2975995_2976095
GAAAAAAGCTAAAAAGCATTCCAACTCCCTTTGCTCTGATTCAGTAAAAGCGAATGGAGGGAGATTACAC
GAGATAAAGAACGCGAGCGACAGTAAATTAG
>Cse1_wt_endogenous_2983007_2983107
CCAAACTCCGCTACCGCGCGATCCAGCAGTGCTGGAATACCATCAATCTTTCGCAGATCGGCGGTCAGGC
TTAAAAAACGACGCCCCAGCGCTGTGACCTG
>Cse1_wt_endogenous_3013282_3013382
TTCGCCATCGAACACGGCCTGGGCAAACGCCACGGTACAACGAATCACTGTCGGTTTTTCCACTTCCAGC
ATGATCACTTTAAAACCCGCATGATACAGAC
>Cse1_wt_endogenous_3016611_3016711
ACGCTGAATAAATTACTCCAGGATGACGCATTTTTAGCTCGCCACGGATTGCAGGAAAAACGCGAAAGCT
TGCAAGCCTTACCCGCTCGCATCCCCACCAG
>Cse1_wt_endogenous_3026095_3026195
CGTCGTTATGAGGATTTAGAGTACGCCCGCGAAATGTCGGCGTTCTTCATCAAGCAGCTTTTACGTAACG
GAACCACCACGGCGCTGGTGTTTGGCACTGT
>Cse1_wt_endogenous_3071915_3072015
GCTTCGTACAGGGATTTACCCACATCGTGGCCTTGTGCCGCGATAAAGGTGTTAGCGATACCACCACCAA
CAATCAGCTGGTCAGCGATTTTAGACAGGGA
>Cse1_wt_endogenous_3104238_3104338
GATTAATGAAAAGAAACTCAACATGATGAATGCCGAGCACCGCAAGCTGCTTGAGCAGGAGATGGTCAAC
TTCCTGTTCGAGGGTAAAGAGGTGCATATCG
>Cse1_wt_endogenous_3123335_3123439
CCGCTGACCATCGCCCCTTCCTGCGGGATGATGTCGCTGCCGTATAACGCATCGTACAGTGAGCCCCAGC
GAGCGTTCGCCGCGTTCAGCGCGTAGCGGGCGTTC
>Cse1_wt_endogenous_3189538_3189638
GATATCGAAAACTCGGTTATCACCACCACCTTAACCGAGGGTGCTATTGGCTACGCCAAACTTGATACCC
GCAAGGGCTACCAAATCATAGGGGTTATTCG
>Cse1_wt_endogenous_3194979_3195079
CTCAGGACGTTGCAGCGTTTTGCGTGACCGCTCGGGGAAGGCAAAATTGCCTCTGGGAAAGCATTGCGCG
GGGTCCGGCGCTCATCAACAATCGGGGGGCA

>Cse1_wt_endogenous_3208603_3208703
TGTGGATGGGGATTGTTGCCGACTGCGTGACCAGCGCCATTTTCTTGACGGCGATGGCACCAAACTTGCT
GTTAATTGGACTGATGAAAAGCGCATCTCAC
>Cse1_wt_endogenous_3209968_3210068
GCAGTACCCTGAGCCGCCATTTTCGACAGTAACGGCCCGCCAGGATAATCCAGCCCCAGCAGCTTCGCGG
TTTTATCAAACGCTTCCCCGGCGGCATCATC
>Cse1_wt_endogenous_3213546_3213646
CCCGAACGCAGAAGAAGATCTGGCACCTACCGCCACTCACGTCGGTTCTGAGCTTTCCCAGGAAGATCTG
GACGATGACGAAGATGAAGACGAAGAAGATG
>Cse1_wt_endogenous_3253206_3253306
CACAGAAGGTACTTCTGGCAGCAATCGTTACGGAAACGATCCGAAGTTTGGTTCAAATTAATCTTAGAAT
TGGGGCGATATTTCGCCCCTTTTTATTAACA
>Cse1_wt_endogenous_3257377_3257477
TAGCGAACCGACGTTATACAGATGCCACATTCCCACCATCGACACGCCCAGCGCCGCGACCACCAGCAGC
TTGGTCAGCACCATGCCGGTCGAAATTTTGA
>Cse1_wt_endogenous_3263911_3264011
ATCATGCTGATAGTGTTCAGTTTACCCAGCGACACTTTAGTTTTGTCGCCTTTATAACCAAACTTCAGGA
CCAGGCCATTCAGACCTTCCAGCGTTCCCAG
>Cse1_wt_endogenous_3270241_3270341
TCTGTCGTGGACAGTCATTCATCTAGGCCAGCAATCGCTCACTGGCTCAAGCAGCCTACCCGGGTTCAGT
ACGGGCCGTACCTTATGAACCCCTATTTGGC
>Cse1_wt_endogenous_3274610_3274710
CAGGAAGGTGAATAGCGACCAGAAAAAGAGGCTGTAGGTGTAAACTTTTTTCGAGCCAAACTTATCAAGC
AGCCAGCCGCCGGGGATTTGCATCAGCAAGT
>Cse1_wt_endogenous_3346187_3346287
ACCGGAAATAACGTCGAGATCACCGAGGCACTGCGCGAATTTGTTACAGCCAAATTTGCCAAACTTGAGC
AATATTTTGACCGAATCAACCAGGTCTATGT
>Cse1_wt_endogenous_3352532_3352632
GTGAAGCATCAAGGAAGGAACGTAAGAATGAGGATTGCTGCTCGAGCTGAATTTGTGTCTCTTCGCGCTC
TTTGATTTCAATTTTCAGTTGGCCGAAGGTT
>Cse1_wt_endogenous_3354482_3354582
GTATCAGCTTTCCCGATAAGTTGGAAATCCGCTGGAAGCTTTCTGGATGAGCAGCCTGCTCATCATATTT
ATGCAGTAATTGAGATCCCCTCTTCACCGTA
>Cse1_wt_endogenous_3358240_3358340
GACGGTTTCACCGCCAAACAGCAGAAGCTGGCGCTGTCGAAGCTGCTGGAGACTGCCGAACCGCATCCAG
GTAAGGCACTCTACTGCACCGAAAACAACCC
>Cse1_wt_endogenous_3363731_3363831
AAACAGACGGTAGTTCGCACAACTATATTGTTCCCTCCACCCTGTTTAATCAGCATGTAGAAAGCTTCCA
GGGTTATCGCTTCGCGATAGGGCGGGTAGAC
>Cse1_wt_endogenous_3373608_3373708
CGATATACCTTTATACCTGTTATACCAGATCAATTAAGCAACACCCCATACAGAAAGCTTATAATGCGAT
CTGCTTCACTAAAGTGGCATTATTTCTTTTT
>Cse1_wt_endogenous_3380385_3380485
ATGGCGCACGACTATCGCCAGCTGTATCAGCACATGGCAAAAAGCTCCAGCAGCCTGCTGCCGGAACTGT
CTGCTGAAGCAAACCCGTTCCGTAATCGTCT
>Cse1_wt_endogenous_3384822_3384922
ATTAATCAGTCTAAAGTCTCGCGGATGTTGACCAAGTTTGGTGCTGTACGTACACGCAATGCCAAAATGG
AAATGGTTTACTGCCTGCCAGCTGAACTGGG
>Cse1_wt_endogenous_3387952_3388052
ATACAGCCAATAAATGTGCCGATGATGCGCAAAAAGCCACGATAGCGAATAGCGCCAGAATACGGTTCAC
CTCCCGCAGCAAAGGCCGTACCGGCGGCAAC
>Cse1_wt_endogenous_3443883_3443983
GAGCACCACCAGAGAACATGTTAAACATCTCAATGATGGTGCCTCGCTGTTGCTCAAGCAGTTTGGCAAG
TACAGCGGCATCAATACCAGGGATCGGAATA
>Cse1_wt_endogenous_3480316_3480416
TAACAGCACCTGTGCCGCGCCTACGCCAAAAATCGAACGCCGCAGTTGCCAAAGTTTGGAGGGATTCAAC
TCAAGGCCGATGATAAACATCAGGAATACCA

>Cse1_wt_endogenous_3483115_3483215
AAATCCGGCCTGGAAGAGTGCGAAATGGCATGGCTGGAAGCCCAGGAGCAGCTTGAGCAGATGCTGCTGG
AAGGCCAAAGCAACTGATGGCGCAGATAACG
>Cse1_wt_endogenous_3535698_3535798
AACATACGCACTTCGTACGCGAGGACTTTATTAAACTGCTGGAGGCTCGGCAAAATCGCGAAGTTCAGCA
CCACCAGATAAGTCGTCACCAGGCTGGCGAA
>Cse1_wt_endogenous_3546346_3546455
TAAATTTACCCCGCTCTGGTGATTCTCAAACGCCAGATGTTACCCGTATCATTCACATGGGTACCAAACA
TACTCCTGACATCTGACTACAATAATTAGTTTTAGTGGGT
>Cse1_wt_endogenous_3548024_3548124
ATCCTTCAGCAACTTGTTCACGCCATCATCGGCAAACATCGACTCAAGCGTTGCGGAAAGCTTGCGTCGC
CAGTTTTTATACTGGTAACTGGTGCCAGGAA
>Cse1_wt_endogenous_3566229_3566329
CATTTTCTTCATCAATCAGCTCTTCGAGATTTAACCCCATCGCTTCCAGTGCGCCCTGTACATCTTCGTA
AATTCCTAGCGACAACATGGCGTTGGAGAGC
>Cse1_wt_endogenous_3585153_3585256
ATAATGCTTTGTGTACTGCCCATCGCCTCTTTCAGCGCCACTTTCTGACCTTTTGCTTCCAGCAGCTTGA
GCGTATCCGGGCTAAACCCTTTTTCGACACGCAG
>Cse1_wt_endogenous_3613457_3613557
AGCCATTGTCAGCTTGAAAATTTAAGCCTTGCGATCTGCACCCCTACTCCCTTTACGCTCACCGCCGACA
GTGTGCAATGGATCGATTCAGTTAACTGATC
>Cse1_wt_endogenous_3629588_3629688
CCGCACCCACGCTACGCCCGGCAAACCGGTTTTGACATATTCCAGATGCTGCTGGAGTAATTCCGGTGGG
ATACGCGCTTTGACGCGGAACATCAGTTTCA
>Cse1_wt_endogenous_3641882_3641982
AAGACCTCTCGATGCCGATGAAATTTACTCTCGGCATGTTTATGTGCTCACTGGGCTTTTTGACGGCGGC
AGCTGCGGGAATGTGGTTTGCGGATGCACAA
>Cse1_wt_endogenous_3654506_3654606
ACACCTGAGCTGGTCAAATAACACCACCGAAAGATGCAGATGCTTTCTCAGCATCTGCATCATGCATTAC
ATCAAATTAATACACAGTAAGCTAACTATTA
>Cse1_wt_endogenous_3659493_3659593
CGCTGTATCAGATTGATCCTGCACCTTTACAGGCCGAGCTAAACTCCGCCAAAGGCTCGCTGGCGAAAGC
GCTCTCTACCGCCAGCAATGCCCGCATCACC
>Cse1_wt_endogenous_3682075_3682175
ACCCAGGTGGTAAGTTCAGGCACAGGGGTCAATTATGCGCAAACACCCGCACTCGGGGAAGGGAGTGCGG
GCATAAGTGATGAGATTAAGAGGATAATTCG
>Cse1_wt_endogenous_3734537_3734637
CAATCTGGAAGCAGCACGTCTCTCCGGGATTAACGTTGAACGCACCAAACTTGCCGTGTTCGCGATTAAC
GGATTAATGGTAGCCATCGCCGGATTAATCC
>Cse1_wt_endogenous_3755325_3755425
GAATCGTCGGTTCGAGGTAGTAGCCGTCTTTCAGTTCACCTTCCAGCAGCTTGCGCCGCCCGCCTGTGAG
CACGTCAGCGCCCTCTTTTTTACCGATATCA
>Cse1_wt_endogenous_3781024_3781124
CGCCCCCTCACCCTAACCCTCTCCCTCAGGGAGAGGGGACCGTTCGGCGCTGTATGTACTCCCTCACTCT
GAAACGACACCGCACTCTTTTTTTCTCCCTC
>Cse1_wt_endogenous_3788742_3788842
GTACGCGTGTTGCAACAGATGGTTTATAACCTGCCGCCAGACATTACGCTGGTGAAAGCCAGCAGCTTGC
TGAATGAACCGCAGGTTGATACTTCTACACC
>Cse1_wt_endogenous_3792545_3792645
ACGCATTGCGATTCTCCAGACAGGGCAAATTCCAGCACATATTACCCAAACTTATAGGTCGCGACGAGAT
AACGCGTTAACACTTCTGCAAAATTCAGGAT
>Cse1_wt_endogenous_3793084_3793184
ATAATATATACATGATTGCCAATCATGAAATCAAAAACCTGTTGTTGAGATGTGACAGTAAGCCCTTTTT
TTATAGTGAAAAGATGCATCTTACTTTCATC
>Cse1_wt_endogenous_3812449_3812549
CGCGTTCCGGTACGCAGAAATAGCGCCAGCAGCTCGACATCCGTTAAGGCGCTAATACCAAACTTCAGCA
TTTTTTCGCGCGGCATCAACAGCTGTGAATT

>Cse1_wt_endogenous_3821807_3821925
CAAAATTGAACTGGACCGCCGTCTACGCGGTCGCGGTCAGGACAGCGAAGAGGTCATTGCAAAGCGTATG
GCGCAAGCTGTTGCAGAAATGAGCCATTACGCCGAATATGATTATCTGA
>Cse1_wt_endogenous_3850566_3850671
CGCAAATACACACCTGCACACCGCGCCCCGGCAGCCCCGCCAGCGCCTCGCGCCCCGAACCAAACTCGGC
AACTACCTGCAAATCAGGTTCCAGCCCCAGCAGCTG
>Cse1_wt_endogenous_3866903_3867003
TTCTCCTTCTAAGAAGCGAGTAAGTACCTGCAAATCCGAAGATTCGCATATGCTCCCTGACGGCGAGCAT
GGAGATGTCAGGCCGCGCCAGGCGGCCTTAG
>Cse1_wt_endogenous_3875105_3875205
GAACGATCTGTTTACCCAGCGTAATGACAATGCGATCGGTTTTATTGAGAGTCATGGAGAGTCCTTGTGC
TCTGTATGTTCTTCTCTACTTTACCCCGATC
>Cse1_wt_endogenous_3880735_3880835
ATTGCGCTGCTTGAGCAATCGCTTGAGATTGCTCCAGGCGGTGAAAAATCCGGGTTCGTTGTGAAAGCAT
CCCCAGTCGAGGAATGCTCTTCTGTATTTGG
>Cse1_wt_endogenous_3884646_3884746
CCAAGAAAAACGTTCGACGCGCCCATGAACGCAATCGGATTAAACGTCTGACGCGTGAAAGCTTCCGTCT
GCGCCAACATGAACTCCCGGCTATGGATTTC
>Cse1_wt_endogenous_3918544_3918644
GGCGCATACTGTTTCTGTTTCAGCAGTTCGGTCACTTTCTGACCGTGGTCAAGCTGCTTACGTGTTGCAT
CGTCAAGGTCGGATGCAAACTGAGAGAACGC
>Cse1_wt_endogenous_3922150_3922250
AAAACCGCCAACGCCACCACCAGTAACACCAACATCGCCAGAACTTTGAAAGCTTCGCCAAATGCGAATG
TCCAGGCCACCCGGCCTTTCGCTGGTGTATG
>Cse1_wt_endogenous_3948029_3948129
CGTAACTAATACTCCGCGCCATAACTAGCTCGGTCAAAGAATTAGGAGCGTGCAGGATGGCGGAAAGCTT
TACGACGACTAATCGATATTTCGACAATAAA
>Cse1_wt_endogenous_3967879_3967979
TACTTCTGCTAATAATTTTCTCTGAGAGCATGCATTGTGAATTTACTGACAGTGAGTACTGATCTCATCA
GTATTTTTTTATTCACGACACTGTTTCTGTT
>Cse1_wt_endogenous_3979559_3979659
TACCTGGGAAGTGCTGAATAACCACTCCGGACTGGCGATCTCGCCTACGCTTATAGGCTCACTGGTGGTG
ATGGGCGGCGCGTTGTTCATCCCGCTCGGGG
>Cse1_wt_endogenous_3980803_3980903
TTACGAAAACATTTGCAACACTCGATGTACCCATAACGATAACCGGTAACACCGGAAAGCATGCAAACAC
AACACGAGGATTTATGGCAGATAACAAACCA
>Cse1_wt_endogenous_3987290_3987390
TATACGCCTGTTCCGCCAGACGTAATACTTCCGGATGGCGTGGCGTAACTTCCAGCAGCTTATCCACGCC
GTGGCGTGCAGCATGGTTTTCATTACGGGCC
>Cse1_wt_endogenous_4010240_4010340
TTCTCCCGACCATACGTTATCCCCTTACGCCAAAGAAACTCTGAAGCTGCTCACCGCGCGCGGCATCAAC
TTTGTGTTTGCGACCGGTCGTCACCACGTTG
>Cse1_wt_endogenous_4024626_4024726
ATGAATCACTTCTGGGTCAAATTCCACAAACAGGTAGTTGGGGAACAATGGCTCACTGACTGCAGTACGT
TTTCCACGCACGATTTTTTCCAGGGTGATCA
>Cse1_wt_endogenous_4029825_4029925
GTGGATTGTGGAGATCACGCCAGCCAGTTTCAGACCATCGATCTTGCCGCGCTCAAGCTGCTTGTTCAGC
AGTTTCGCGGCTTCGGTCATGCCGAGGGTTA
>Cse1_wt_endogenous_4053720_4053820
CGCCACTCGATACCAGATTAATTATTGATTCCAGCATAATGACTCTCCCCGTTTTCCGGGCAAGATCATA
CTGAACTTATCGGAACAGTAAAGCGTAAAAT
>Cse1_wt_endogenous_4055031_4055131
CGGTCATAATGATGACCGGAAGCATTGGATGGCGCTGTTTAATCTGCTTGAGCAGCGCCAGCCCGTCCAT
TCCCGGCATACGGATATCTGAAAGCAGCACA
>Cse1_wt_endogenous_4071169_4071269
CGATCCAGCCATTTTTTGTCGTGAGTGACGTCATAAACAATCAAGAAAGCTTCCACCGCGTGCATATTGG
CATTGCCGCCGCGGTACTCTTCGGTTTTGCT

>Cse1_wt_endogenous_4071304_4071404
TCTTCTTCGCTCCAGAAATATTTCTCGATAATTTCAATGGTGTAATCGAGCAGCTTGCGCGCTTCCGGGT
GACCCGTTGTGACGGCGCTGGCGGCACCCAG
>Cse1_wt_endogenous_4117590_4117690
GTTAACGCCAGGATCAGCCCCATCAGAATAGCGGTAATCACCATCTCAACTGCGAAAGCCTGCACAAAAT
TGATATGAGGATTAGGGTAAGTAGAGAAAGT
>Cse1_wt_endogenous_4119584_4119693
CAGCCAGCCCCACAGGCTATGCAGCGAAAAGAGATTAAACAGCGCCAGACACACCAGCGAGCCCATCAGC
AGGCAGGCATGATAACGACGCGCGTTCACTTCACCTAAGC
>Cse1_wt_endogenous_4131427_4131527
TTCCTGTGGATGCGCGCCCATCCGTATGATGATTTAGTGGTGCTGGACGTTACCGCCAGCCAGCAGCTTG
CTGATCAGTATCTTGATTTCGCCAGCCACGG
>Cse1_wt_endogenous_4135499_4135604
CGCGTTGATGCGCGTCAGGATCAGACTGACATTGAGATGTTTGAGCTGCTGGAGCCAATTGCTGACGGTT
TCCGTAACTATCGCGCTCGTCTGGACGTTTCCACCA
>Cse1_wt_endogenous_4220338_4220438
CTTCCATATAATTTTTCTCCGCAATGTATCGAGGGTTATCCGTAAAGCCAAAGCTTTCAGCCATCTTATT
TAATGTATTAAGGATTAATTCAGCAATAACC
>Cse1_wt_endogenous_4226928_4227028
AAAAACAGGCTTCGCTAACTACTGTCTCGCTGACTTCGTTGCGCCGAAGCTTTCTGGTAAAGCAGATTAC
ATCGGCGCATTTGCCGTGACTGGCGGGCTGG
>Cse1_wt_endogenous_4260060_4260160
CACCGAGAACCCTTATTGTTGCCGTAATGTTGATTTTCTGTTTTGTAGGTAAGGTGTTATGTTGCCTTGT
CGTACCATTATCAACACGATAATAATTAATA
>Cse1_wt_endogenous_4285393_4285493
CGCATCGGGCAATTGTGGGTTACGATGGCATCGCGATAGCCTGCTTCTCTTCAAGCAGCTTCTCGACTAC
GCCAGGATCGGCAAGCGTCGAGGTATCGCCC
>Cse1_wt_endogenous_4323191_4323291
CCAGCTGCGCTTGGGTGGGGCGATGGTGATGGTTTGCATGTTTGGCTCCGGTCTGTAGGCCGGATAAGGT
GCTTGCACCGCATCCGGCATCAACGCCTGCA
>Cse1_wt_endogenous_4331124_4331224
GCCGTTAGGGATTATCGGGCTTTACCTGCGCCATGCGCTGGAAGAGACTCCGGCGTTCCAGCAGCATGTC
GATAAACTGGAACAGGGCGACCGTGAAGGTT
>Cse1_wt_endogenous_4340153_4340253
ATAAAGTCGGCGGTATCTTCCAGAATCCAGGCGAATTCATCGACAAGCTCCAGCAGGTCGCGATCCATTG
CGGCGAGGGCTTTTTCCCGATCGCCCAACAG
>Cse1_wt_endogenous_4368400_4368500
GACCGACGATTATCCCCTGCATCGACCGAATACCCGAGATCATATGCTGCTTGAGGATTTCTACCGTAAT
CTGGATCACTTTAAGTGTCGGTTTTTACCCC
>Cse1_wt_endogenous_4383067_4383167
GGCAGCAGAAAGCCTTTCTTATCAACAAGCTTTCTTGCGTTATCTGGAAATTGACCCGCTCTCTGCCGAC
AAAACGCAACTGCGGGAAGTCGCAGCGAAAC
>Cse1_wt_endogenous_4384102_4384202
TTTATGTGGTTTTCCTCTTATATCATCTGGATGGTGAGTACCTCCGCGAAAGTTTGGGTACCGTTCTCAA
CATTCCTCTATGGTAGCGACATGACCCAGCA
>Cse1_wt_endogenous_4395337_4395437
TTAGGCATTATTGATGCCGGAAATGCAGGCATGGCGAGCGGCGGCATGGGCGATGTGCTCTCTGGTATTA
TTGGCGCATTGCTTGGGCAAAAACTGTCGCC
>Cse1_wt_endogenous_4406380_4406480
CATTCGCCTGGGTAAACCGGCGAGTGCGATACGTATTGGTGATGTGGTGCGCGAGCTGGAGCCCTTATCG
CTGGTGAATTGCAGCAGTGAGTTTTGCCACA
>Cse1_wt_endogenous_4407613_4407713
ACGTCGGTGTACTTCCCTTCGCAGGTTATCCCGATGCTGCCGGAAGTGCTCTCTAACGGCCTGTGTTCGC
TCAACCCGCAGGTAGACCGCCTGTGTATGGT
>Cse1_wt_endogenous_4413315_4413415
TCGCTGTATGGACGGATGGATTTTGCCTGGTGTGGCAATGCGCCGGTGAAGCTGCTGGAGTACAACGCCG
ATACGCCAACTTCATTGTACGAGTCGGCTTA

>Cse1_wt_endogenous_4456691_4456791
ACGCGCTGGGTTCGTTTATTAATGCTCGTCGCCGTCTGGAGTTGCGTGGTGAAGCGAATGGCGTCACGGT
ATATGACGATTTTGCCCATCACCCGACGGCG
>Cse1_wt_endogenous_4457874_4457974
CTTAAAAAAAGAGGCTAATGTTACCAGTTAAGATGCGCACTGAAAAACGGTTCTCTGTTAGACTTCAGAG
AAACTCTCTACATTATGGCACTTGCAATGAA
>Cse1_wt_endogenous_4461302_4461413
TTCACGCCTTCGAGACGAGCGATACGGGTCATCAGCGCCTTACGTGCCAGCACCAGACGTTCATCCAGCA
GCTTCCAGAAGGTGGCTTCATCGCCTTTTGCTTCCAGAGCAA
>Cse1_wt_endogenous_4484054_4484154
CGCGCGGTCCTTCTCCCGCTAAATTATGCGGAACAAAGCTTTCTGCCGGACGCGCCCACAGGGCTTCATC
CAGCCGGTAAGCCTGCTTTTCATCTTCACAG
>Cse1_wt_endogenous_4485959_4486059
CTCCTGAATCTTAAAGACAACGGCGGTGGCTACAGATAGAATTGCAAGCTTTCGTAACTCATGTCCGCTG
TTGCGATGACTTCGTGTTAATCTTAACGTTA
>Cse1_wt_endogenous_4519731_4519831
TGTAACTCCATGGCCAGTCTGGCGCTTGATACCACGGCGGGCACGTGCAGCGAGGCGACACGGCCTGAAA
GCTTGTAGGCGATTTGCGATGAGGATTCAAT
>Cse1_wt_endogenous_4539058_4539158
TCACGAGACATATCACCGGATAAGTCTTGTTGTCGGTAAGCTTTCCAGTCGTAACGATAGCGAATGCCAA
AATTAAGATCTTTTGTCGCGTCCCAGGACAG

Regions used to identify motif shown in Figure S3

>Cse1_deltaCR1_endogenous_113333_113433
GCGTAAAAAAAGACAATTTCGCAGTCTTGCGCCGCATTGATTAGTGCGTATGATAGCGTCACTGGAGTTG
CGCTCTTACCCTTATAGCCATTAACCCCAGG
>Cse1_deltaCR1_endogenous_657201_657320
GTAAAGTTACACTGGACAAAGCGTACCACAATTGGTGTACTGGTAACCGACACAGCATTTGTGTCTATTT
TTCATGTAAAGGTAATTTTGATGTCTAAGATTAAAGGTAACGTTAAGTGG
>Cse1_deltaCR1_endogenous_1641603_1641703
AAACGACTTCGTACTTAATTGGAGAGACTCAAAGAAGGAATAAGTGAATAACACCTGAAATGAGAACTGC
TTTAGTAAACTACTTCGTATATCGTCTGTTC
>Cse1_deltaCR1_endogenous_2416201_2416350
GCCCGCGAACAGCTGGAAGACAACGTGGTGCTCGGTACGCTGATGCTGGAACAGGATGAAGTTGATGGTC
TGGTTTCCGGTGCTGTTCACACTACCGCAAACACCATCCGTCCGCCGCTGCAGCTGATCAAAACTGCACC
GGGCAGCTCC
>Cse1_deltaCR1_endogenous_2531167_2531314
TACCAGTAAAGCGATTATGGCGATCGCGCCAACAATGATTAATATCAGACGCAAATCCTGCATCATTATA
TTCTCTGTTGTTCTAACACCTTGCCACCACGGCAAACATTTACTCACTAAGAGTATTTGCCGATTACCTC
AAGTGCAA
>Cse1_deltaCR1_endogenous_2535622_2535757
CTGGACGAATTCTCTATGAGCGCCATTTCTATCCCGCGCATTAAGAAGATTATCCGTAACACGAACTTCG
AAGATGCGAAGGTGTTAGCAGAGCAGGCTCTTGCTCAACCGACAACGGACGAGTTAATGACGCTGG
>Cse1_deltaCR1_endogenous_3071923_3072023
CAGGGATTTACCCACATCGTGGCCTTGTGCCGCGATAAAGGTGTTAGCGATACCACCACCAACAATCAGC
TGGTCAGCGATTTTAGACAGGGAGTCCAGAA
>Cse1_deltaCR1_endogenous_3089483_3089583
GTTGGCGCAACGTTCCTGACCATGCTCAACACGCTGGGTAACGCCAACACCTTCTGGGTGTATGCGGCTC
TGAACGTACTGTTTATCCTGCTGACATTGTG
>Cse1_deltaCR1_endogenous_3308451_3308551
TGATAAAACGCCAGCAGATCATCTTGCGCTAACTTGTCACGACCGCCGTAATATAATGCGATCCCGCGAT
TCAAGTGCGCGTAGTTGTAAGTTGGATCAAG
>Cse1_deltaCR1_endogenous_3308602_3308702
GCGTTAAATATATGCCTAAGTAATTGAATACTTCAGGCATATCCGGTCGGATTGCCAGCGCTTGCGAAAA
ATCGTTACGCGCTAATGCCCTCAGACCGAGA

>Cse1_deltaCR1_endogenous_3384656_3384756
TCTGTATGCACAATAATGTTGTATCAACCACCATATCGGGTGACTTATGCGAAGCTCGGCTAAGCAAGAA
GAACTAGTTAAAGCATTTAAAGCATTACTTA
>Cse1_deltaCR1_endogenous_3445383_3445483
CAGCCGCGTCTTTGTTACCGGTGTACTTCAGTTGTTCAGCGATAGCTTTTTCTACAGTAGAAGCAGCTAC
CAGAACTTCAGAACCGTTCGGTGCAATTACC
>Cse1_deltaCR1_endogenous_3719887_3719987
CAACGGTTTGACGTACAGACCATTAAAGCAGTGTAGTAAGGCAAGTCCCTTCAAGAGTTATCGTTGATAC
CCCTCGTAGTGCACATTCCTTTAACGCTTCA
>Cse1_deltaCR1_endogenous_3737195_3737306
TTAATCATTCCAACACCTTATATTTTCACAAATTTGAGAGTTGAATCTCAAATCATATCAAAAATAGCT
GTCAAGAGCACCCCAAGGAATAGTCCAAATCTGAAACTATGT
>Cse1_deltaCR1_endogenous_3888394_3888494
GTAATAATGTAGCCTCGTGTCTTGCGAGGATAAGTGCATTATGAATATCTTACATATATGTGTGACCTCA
AAATGGTTCAATATTGACAACAAAATTGTCG
>Cse1_deltaCR1_endogenous_3888592_3888707
TTGCAAAGGTCATCTCTCGTTTATTTACTTGTTTTAGTAAATGATGGTGCTTGCATATATATCTGGCGAA
TTAATCGGTATAGCAGATGTAATATTCACAGGGATCACTGTAATTA
>Cse1_deltaCR1_endogenous_3946955_3947055
TAGGGGCGTAGTTCAATTGGTAGAGCACCGGTCTCCAAAACCGGGTGTTGGGAGTTCGAGTCTCTCCGCC
CCTGCCAGAAATCATCCTTAGCGAAAGCTAA
>Cse1_deltaCR1_endogenous_3966191_3966316
TGAATCTTGTAATTTCCAACGCTTCCCGTTTTATCTTAAATGCGAAGTGAACAGATTTCTGGCTCGTCAC
TCAATCCGTCTTGTCGTTTCAGTTCTGCGTACTCTCCTGTGACCAGGCAGCGAAAA
>Cse1_deltaCR1_endogenous_4260020_4260157
GGGTTAAGAATGTCCTTACTTTACCATGTTCCAGGAAAAACACCGAGAACCCTTATTGTTGCCGTAATGT
TGATTTTCTGTTTTGTAGGTAAGGTGTTATGTTGCCTTGTCGTACCATTATCAACACGATAATAATTA
>Cse1_deltaCR1_endogenous_4383941_4384062
GTTATAGTGCGATTCCCTTTTATATATTTTCTGCATTGTTATTCTTTATTCCATTCGCCTTAATGATGGC
TGAAATGGGAGCTGCTTATCGCAAAGAAGAAGGCGGTATCTATTCCTGGATG
>Cse1_deltaCR1_endogenous_4465785_4465885
CGAAAACCGAGGATCAAACCGCCGCTAATCAACGCGGGCAGCAACGGGAAGAAGATCACCGCGAAATGAG
AGATCAACTGCTCATGCCATTTCATATTATG
>Cse1_deltaCR1_endogenous_4572138_4572238
AATTGTTTTATACCCTGGAAAGTTAAATGTCAGCTACTGAATACTTTTTGATTGTTTGAGATTTATTTTC
ATTTGAAATTATAAAATCAGGTGATAAATGA

Regions used to identify motifs shown in Figure S4

>Cse1_deltaCR1_plasmidspacer8_5_117
TTCATTCTGACTGCAACGGGCAATATGTCTCTGTGTGGATTAAAAAAAGAGTGTCTGATAGCAGCTTCTG
AACTGGTTACCTGCCGTGAGTAAATTAAAATTTTATTGACTTA
>Cse1_deltaCR1_plasmidspacer8_19062_19179
GCAGTGGTAGAAGGCGAGCCCATTCATCTTCGCTGCTTCGAATCCACCCACGAAATGCTGCTGGAGCAAT
TAAGTCAGCATAAACTGGATATGATCATTTCTGACTGTCCGATAGACT
>Cse1_deltaCR1_plasmidspacer8_28724_28853
GCTGCCGATATTGCGATTGTCTTTGCTGCCAATTTTAGCGTTGGCGTTAACGTCATGCTTAAGCTGCTGG
AGAAAGCAGCCAAAGTGATGGGTGACTACACCGATATCGAAATTATTGAAGCACATCATA
>Cse1_deltaCR1_plasmidspacer8_70800_70909
AACGCCGGGCAAGGGGAAGGGCGCTATTCGGAGCTGCTGGCGATAAATCTGCTTGAGCAATTGTTACTGC
GGCGCATGGAAGCGATTAACGAGTCGCTCCATCCACCGAT
>Cse1_deltaCR1_plasmidspacer8_157875_158002
CTGCAGTTCAGCGTTACGCTCAACTTCAGCTCGCAAGGCCAACAGGTCATAAGCCGCACGGAACTTAGGA
TGCTCCAGCAGTTTCCATGCGCGTTTACCCTGACGACGGGACATACGCAACTGCAACT
>Cse1_deltaCR1_plasmidspacer8_165045_165163
TTTATGGCCGAATGGTCAATCTTGAGCCAGACATGACCATCAGCAAGAACGAGATGGTGAAGCTGCTGGA

GGCGACCCAGTATCGTCAGGTGTCGAAAATGACCCGTCCTGGCGAATTT
>Cse1_deltaCR1_plasmidspacer8_222432_222532
CGTGGTCAGTTCCTGGCCATCGACCAGCACGCTACCCTCGGTTGGGCGCTCCAGCAGGTTTACACAACGT
ATAAGCGTACTCTTACCCGCGCCTGAGGCAC
>Cse1_deltaCR1_plasmidspacer8_261194_261321
TTCGGTGCGCCGCCGGCGGGTGAAATCACGGTAGATGAAGGGGCAACTGCCGCCATTCTGGAACGCGGCA
GCTCCCTGTTGCCGAAAGGCATTAAAAGCGTGACTGGCAATTTCTCGCGTGGTGAAGT
>Cse1_deltaCR1_plasmidspacer8_269916_270016
GTCACTAATGCCAACCCCTGCTGTAATTCGTTAAATCCCCGCTGTATGTGCTCAAGCAGTATTTCACCTT
CTTTCGTCAGCGTAATTTCTCGCGTACTGCG
>Cse1_deltaCR1_plasmidspacer8_298713_298813
GCGGGTATGACATTGTTCCTCCTTAAACCACATCCGGCAGCTTATCGAGCAGCTTATCCAGAGTGATGGG
ATAATCCCGTACCCGAATACCGGTGGCGTTA
>Cse1_deltaCR1_plasmidspacer8_332420_332526
TCATTTGTCATTGCCTGGTAGCGTATCCGAGAAAGAACGACTGCTACTCAAGCTGCTGATGCAGGGAATG
TCTGTAACAGAAATATCACAGTACAGAAATCGCAGTG
>Cse1_deltaCR1_plasmidspacer8_413347_413447
CTGAGTGACCAGAGTTTGCGCCTGACGGCGCTGGTTTTCCAGATTCTGCTTGAGCTGTTCCAGCTGCGTT
AGTGTTTGTTCATCCATTAGCGCCGCAAGGA
>Cse1_deltaCR1_plasmidspacer8_419968_420068
CTGGTCTATTTCGCTATCGTTATTCTGGTTTCGCTCTATCCGGGCAAGCTGCTGGATACCGTGGGCAACT
TCCTTGCGCCGCTGAAAATTATCGCGCTGGT
>Cse1_deltaCR1_plasmidspacer8_483953_484062
CGTCATGGTGCCATCGGTGTTGATAACGCCGACAACCATCAGGAAGCTGCTGGATGATTTCTCAACGCTC
ACCCCTTGCTGCTGAACTTCTTGCGGCAGCAACGGCATCG
>Cse1_deltaCR1_plasmidspacer8_557431_557562
CCGCCGGTAACGGCAGTTGAACCAGAATGCCATCGATGGTGTTGTCGGCATTCAGCGTATCGATAAGCTC
CAGCAGCTCCGCTTCGCTGGTGGTTTCCGGGAGGTCATAAGAGCGGGAGACGAACCCGACTT
>Cse1_deltaCR1_plasmidspacer8_650790_650890
CTGTTTCTCATGCGGGGGTCTTTTGACGAGCTGCCGCGTCCTGGCGGGAGTGCTCAAGCAGGTTCTGCAA
ATAATGCAGCGTGACTGCAGGGACCAGCGGC
>Cse1_deltaCR1_plasmidspacer8_659505_659620
TCCCACCATCAGACCAGACTTGGTCGGGATTTCCGGATGCGCTTCTTTAAAGCGTTCCAGCAGCTTCAGC
GACCAGTTGTAATCTGCACCAGGCCGTACCTGACGGTAAATACGCG
>Cse1_deltaCR1_plasmidspacer8_684859_684959
CTTATTGGCGTGAGCAGAAAGCCCGACACGCTCCAGCAGTTTCAGGGCTTTTTCACGAGCCGGCGCTTTA
TCGCGTTTAAGCACTTTCACCTGCGCCAGGG
>Cse1_deltaCR1_plasmidspacer8_757714_757814
CTATAACCCGGATGTTGATGATGCTCCGCGTATGCAGGATTACACCCTGGAAGCGGATGAAGGTCGCGAC
ATGATGCTGCTGGATGCGCTTATCCAGCTAA
>Cse1_deltaCR1_plasmidspacer8_838689_838789
ATAAATGATGCCACTCGTACGCCGCGGGTTACGGGTGTCACGCAATGCAGGCTGCTGGAGGGATACAACA
CGAGATCGCCTGCCGGGAGTTTTACCCGATG
>Cse1_deltaCR1_plasmidspacer8_847166_847278
AATGGCCCCGGAGCGGGTTTGCGCCCCTCACCCTAATCCTCTCCCCATAGGGGAGAGGGAACTGCCAGTG
CGTTTTACAGGTGTAGCGTTATTATTTCGGTTCAGTACCGAAC
>Cse1_deltaCR1_plasmidspacer8_917055_917155
CACATCCGCAAAATCTCTCTGATGGGCAGATTCGTCAGGGACTTTCCGCAATGGTACAGCTGCTTGAGCA
TTATTTCTCTGAGCAGGGGGCCGGACAGGCG
>Cse1_deltaCR1_plasmidspacer8_927773_927882
CTGGCGTCCGCCTTCACCTAACCAACTGTTGAGACCTGCATCCTCGAGCAGCTTTTCCAGGCCAACGCGA
CGCAAGATCTCCGACAGAGCCTCATCACTACTGCCAGGCG
>Cse1_deltaCR1_plasmidspacer8_953234_953334
TTCAGCGGAGCTTCAGTCTGCAGACCAACGATTTTCTCAAGCTGCTTGTTGATGTAGCCAGCGTCGTGAG
AGGTGATGGTGGAAGCAACAGCGGTGTCAAA
>Cse1_deltaCR1_plasmidspacer8_1290395_1290516
CAGACCTGATGATATGTGATATCGCGATGCCACGAATGAACGGGCTTAAACTGCTGGAGCATATACGTAA

CAGAGGCGACCAGACCCCAGTTCTGGTGATATCTGCCACTGAAAATATGGCA
>Cse1_deltaCR1_plasmidspacer8_1428358_1428458
TTGAACAATTGACGGGCCAGTTGATGCTGCTCGAGCAGGTGGCGGAAATTCATGATGGTGGTGCGATCCG
GCAGGGCGCTATCCAGGGATAATCGGGCAAA
>Cse1_deltaCR1_plasmidspacer8_1523921_1524021
GGCGAATTGATTACCATCGACACCGCCGACAATAAAATCCTCAGCCGTAAAAAGCTGCTGGATGACGGCA
AAGAGCACTTCTTTATCAACATTAGCCTTGA
>Cse1_deltaCR1_plasmidspacer8_1584536_1584636
GTGGCTGCGTACTGTCGCCATGACCAGCTTACTGTTAAACGCTGAAGAGGGATCTTCTAACAGCCCTTTG
CTGGTAATGAAATTAGCCTTACCTGACGGCG
>Cse1_deltaCR1_plasmidspacer8_1799327_1799433
TAAAAGTCAATGTTTTGATGGCGTTGAAACGAAAAGAGGGAGACTAGCTCCCTCTTTCAACTGGCTTATG
CCAGAGCTGCTTTCGCTTTTTCAACCAGAGCGGTGAA
>Cse1_deltaCR1_plasmidspacer8_1898886_1899018
CTGCAGCGGGCAGTAGCAATGCTGGGGTCATGGAGATGATTAACATCGACTTACTGACGGCAATGCTGCT
TGAGCCACAACTGCCGCAAATCAGTAGCGCCAGCCTGACGGTGGACAAACGGCATTTGCTCTA
>Cse1_deltaCR1_plasmidspacer8_2030922_2031030
TGCCTCTTGCTGACGCAACGCCACCGCCTGTTTGATTTTTGGCTCAAGCAGTTTTGCCACCGCGGCAAAG
ACCGGCACGACCACCGAGTTACCGAACTGGCGATAGGCC
>Cse1_deltaCR1_plasmidspacer8_2336118_2336223
TGATTTCCGTTTTGCTTAACTGGCTGTCGCTGGTAGCGGCAAACAACAATGGCTCCAGCAGTTTGCTTAA
ACCACTGCGCTGGATCAGCACCATATAATGTGAAAG
>Cse1_deltaCR1_plasmidspacer8_2406037_2406137
CCATATCGCGAAACGGGCTAGGATCATCCAGCAATACAAGAGGGATCGGCTCGCCTTTTTGCAATATGTA
TTCCGCTGCGCAGTACCAGTGTGTTGGCGAG
>Cse1_deltaCR1_plasmidspacer8_2513980_2514080
GCGACCAAACTGGTTTCCAACGAGTTCGTTGCGATGATGGATCTGCAGAAAATTGCTTCCACGCTCTCTC
CGCGTGCTGAAGGCATCATCTCTGTGTTCCT
>Cse1_deltaCR1_plasmidspacer8_2532659_2532779
CGGTTACAAACTCACCCTGACCATGCCAGAAACCATGAGTATTGAACGCCGCAAGCTGCTGAAAGCGTTA
GGTGCAAACCTGGTGCTGACGGAAGGTGCTAAAGGCATGAAAGGCGCAATC
>Cse1_deltaCR1_plasmidspacer8_2535662_2535762
TTAAGAAGATTATCCGTAACACGAACTTCGAAGATGCGAAGGTGTTAGCAGAGCAGGCTCTTGCTCAACC
GACAACGGACGAGTTAATGACGCTGGTTAAC
>Cse1_deltaCR1_plasmidspacer8_2559031_2559131
GGAGTTTACCCGCTAATCACACTCAAAGAAGCACGCAGGCGCGCCACGGAGAGCAGATCCCTTATTGCCA
ATGGAATTAACCCAGTGGAACAAGCCCGCAA
>Cse1_deltaCR1_plasmidspacer8_2622895_2622995
CTCTGCCGCTGGCGTTTTTCAATTCACCTGTAAATCGCAAGCTCCAGCAGTTTTTTTCCCCCTTTTCTGG
CATAGTTGGACATCTGCCAATATTGCTCGCC
>Cse1_deltaCR1_plasmidspacer8_2740715_2740821
CGCTTCCGTCGCCAGTGGCAGTCCCATATTCACCAGCTCAAGCAGCAATTTACGCGCGATCTGCAGCCCG
GCTTCTACATCAAAAGAGCCATCCATATGGGGATCGT
>Cse1_deltaCR1_plasmidspacer8_2800931_2801042
CCGTCTGCATCCAGCGCCACACGCGGCCAGTGCATGAGCCAGCTCGTCTACTTCTCCAGCAGCTCCGAAA
ACACGCAGCGTTTTATCGAACGTTTAGGTCTGCCCGCGGTGC
>Cse1_deltaCR1_plasmidspacer8_2810808_2810943
TTTCGCGCCAGCCGCCACGAAGATTTTCCTTATCAGGAGATCCTTCTGACTCGTCTTTGCATGCACATGC
AAAGCAAGCTGCTGGAGAACCGCAATAAAATGCTGAAGGCTCAGGGGATTAACGAGACGTTGTTTA
>Cse1_deltaCR1_plasmidspacer8_2917282_2917382
CCGGTTATCGCGGTAACGGCGCATGCAATGGCCGGGCAAAAAGAGAAGCTGCTTGGCGCAGGGATGAGCG
ATTATCTGGCGAAACCGATTGAAGAAGAGCG
>Cse1_deltaCR1_plasmidspacer8_3026117_3026217
ACGCCCGCGAAATGTCGGCGTTCTTCATCAAGCAGCTTTTACGTAACGGAACCACCACGGCGCTGGTGTT
TGGCACTGTTCATCCGCAATCTGTTGATGCG
>Cse1_deltaCR1_plasmidspacer8_3071924_3072024
AGGGATTTACCCACATCGTGGCCTTGTGCCGCGATAAAGGTGTTAGCGATACCACCACCAACAATCAGCT

GGTCAGCGATTTTAGACAGGGAGTCCAGAAC
>Cse1_deltaCR1_plasmidspacer8_3104247_3104347
AAAGAAACTCAACATGATGAATGCCGAGCACCGCAAGCTGCTTGAGCAGGAGATGGTCAACTTCCTGTTC
GAGGGTAAAGAGGTGCATATCGAGGGCTATA
>Cse1_deltaCR1_plasmidspacer8_3120162_3120283
TAAACACCGCATCCATTGGCGTTGGTTGGGCGACAATGGGTGCCGCTTTCAACACTTGTTGATGCAAATG
AGGGATCTGGAAATTAATCACCAGTGAATAAAACGCGCCGCCCGGAGCAAAT
>Cse1_deltaCR1_plasmidspacer8_3209975_3210075
CCTGAGCCGCCATTTTCGACAGTAACGGCCCGCCAGGATAATCCAGCCCCAGCAGCTTCGCGGTTTTATC
AAACGCTTCCCCGGCGGCATCATCGATAGAC
>Cse1_deltaCR1_plasmidspacer8_3257387_3257487
ACGTTATACAGATGCCACATTCCCACCATCGACACGCCCAGCGCCGCGACCACCAGCAGCTTGGTCAGCA
CCATGCCGGTCGAAATTTTGAATAACAATTT
>Cse1_deltaCR1_plasmidspacer8_3270232_3270332
AAGCCGGGTTCTGTCGTGGACAGTCATTCATCTAGGCCAGCAATCGCTCACTGGCTCAAGCAGCCTACCC
GGGTTCAGTACGGGCCGTACCTTATGAACCC
>Cse1_deltaCR1_plasmidspacer8_3330632_3330739
GTCATCCCAGTCTTCATCATCCTCTTCAGCAATCTCTTCAAGCTGCTGGCGATGATAATCATCCCACATG
AATTCGACTTTCTCTGGCTGTTTCGCTTCTTCAGCCTG
>Cse1_deltaCR1_plasmidspacer8_3330833_3330933
TTCCCAGCCCAGCGCCTCAGCGATCGCTTTCGCTTTCTCTTCGGCTTCTACCTTATCCAGCAGATCGATC
TTGTTGAACACTAACCAACGCGGTTTCGTCG
>Cse1_deltaCR1_plasmidspacer8_3336268_3336368
CGCGGGCATCAATATGCACAGAACCATTACGTTCTACTTTCGCACCCAGCTGGCTTAGCAGCTTCATTGA
TGTATCGACGTCTTTCAGTTTCGGGACGTTC
>Cse1_deltaCR1_plasmidspacer8_3352511_3352611
TACGATAAAAAACCAGGTCGGGTGAAGCATCAAGGAAGGAACGTAAGAATGAGGATTGCTGCTCGAGCTG
AATTTGTGTCTCTTCGCGCTCTTTGATTTCA
>Cse1_deltaCR1_plasmidspacer8_3357651_3357751
TATCGAGGACTTAGCGCAGCTCATTTTCGACCTCAAGCAGGTTAACCCGAAAGCGATGATCTCCGTGAAG
CTGGTTTCCGAACCGGGAGTAGGCACCATCG
>Cse1_deltaCR1_plasmidspacer8_3358234_3358345
GAGCTGGACGGTTTCACCGCCAAACAGCAGAAGCTGGCGCTGTCGAAGCTGCTGGAGACTGCCGAACCGC
ATCCAGGTAAGGCACTCTACTGCACCGAAAACAACCCGCCGT
>Cse1_deltaCR1_plasmidspacer8_3380370_3380487
GAATTACTGGATACCATGGCGCACGACTATCGCCAGCTGTATCAGCACATGGCAAAAAGCTCCAGCAGCC
TGCTGCCGGAACTGTCTGCTGAAGCAAACCCGTTCCGTAATCGTCTGG
>Cse1_deltaCR1_plasmidspacer8_3384672_3384772
TGTTGTATCAACCACCATATCGGGTGACTTATGCGAAGCTCGGCTAAGCAAGAAGAACTAGTTAAAGCAT
TTAAAGCATTACTTAAAGAAGAGAAATTTAG
>Cse1_deltaCR1_plasmidspacer8_3443850_3443994
TGATCCCCAGAGCAAAGATAGAAGCACGGCTGAGAGCACCACCAGAGAACATGTTAAACATCTCAATGAT
GGTGCCTCGCTGTTGCTCAAGCAGTTTGGCAAGTACAGCGGCATCAATACCAGGGATCGGAATAAAAGAG
CCAAT
>Cse1_deltaCR1_plasmidspacer8_3535712_3535812
GTACGCGAGGACTTTATTAAACTGCTGGAGGCTCGGCAAAATCGCGAAGTTCAGCACCACCAGATAAGTC
GTCACCAGGCTGGCGAACAGCAAGGTGACGA
>Cse1_deltaCR1_plasmidspacer8_3543270_3543370
CATTCACGCGCTTTGTGATATGAAAACCCTCAAGCTGGAGTCGAGCAGCTTTATTGATGACGATCTGCGT
GAAAGCTATTCCGATGTGCTGTGGTCGGTGA
>Cse1_deltaCR1_plasmidspacer8_3585164_3585264
TGTACTGCCCATCGCCTCTTTCAGCGCCACTTTCTGACCTTTTGCTTCCAGCAGCTTGAGCGTATCCGGG
CTAAACCCTTTTTCGACACGCAGCTCGTCCG
>Cse1_deltaCR1_plasmidspacer8_3629593_3629693
CCCACGCTACGCCCGGCAAACCGGTTTTGACATATTCCAGATGCTGCTGGAGTAATTCCGGTGGGATACG
CGCTTTGACGCGGAACATCAGTTTCAGCCGT
>Cse1_deltaCR1_plasmidspacer8_3729890_3729990

AACAGGCCAAGCGCAATGGCGGTGGCTATTTCATGATGGAAAGAGTGACCTGCCAGCGTCGCGTGGTTAG
CTTCAATGTTCAGTTTAATCTCTTTTTCCAG
>Cse1_deltaCR1_plasmidspacer8_3755301_3755427
CCCGCATATTGTTCTGACCAAACAGAATCGTCGGTTCGAGGTAGTAGCCGTCTTTCAGTTCACCTTCCAG
CAGCTTGCGCCGCCCGCCTGTGAGCACGTCAGCGCCCTCTTTTTTACCGATATCAAT
>Cse1_deltaCR1_plasmidspacer8_3788726_3788826
TCCTCACCCTGCAACGGTACGCGTGTTGCAACAGATGGTTTATAACCTGCCGCCAGACATTACGCTGGTG
AAAGCCAGCAGCTTGCTGAATGAACCGCAGG
>Cse1_deltaCR1_plasmidspacer8_3918555_3918655
TTTCTGTTTCAGCAGTTCGGTCACTTTCTGACCGTGGTCAAGCTGCTTACGTGTTGCATCGTCAAGGTCG
GATGCAAACTGAGAGAACGCTGCCAGTTCAC
>Cse1_deltaCR1_plasmidspacer8_3920387_3920508
TAATAAATTCAGACATCAGCCCCTCCCTCCTTACAGTTCAGCGACAAGTTTATCCACGATGTCGCTGTTA
GCAGCTTCATCCACGGAACGTTCGATGATCTTCTCGGCGCCAGCAACAGCCA
>Cse1_deltaCR1_plasmidspacer8_3948946_3949066
AACAAGCTGGAAACGGGCTGGATAGGTTATTTTTGCTCGTGTGCGTGAAAGATGGATCTGCCCGGATTCA
ATCGGCTCTCGCAAGGCATCCAGTGTACGCCGTTCAAATTCAGGTAGCTCA
>Cse1_deltaCR1_plasmidspacer8_4040583_4040683
CGCCGATGGTAGTGTGGGGTCTCCTCATGCGAGAGTAGGGAACTGCCAGGCATCAAATAAAACGAAAGGC
TCAGTCGGAAGACTGGGCCTTTCGTTTTATC
>Cse1_deltaCR1_plasmidspacer8_4053231_4053331
CGCTGATTACTTTGCGGAAGATCTCAAGCTGCTCGCCCCGTTAGCAAAAGATGGGCTGGTGGATGTGGAT
GAGAAGGGAATACAGGTGACGGCGAAAGGTC
>Cse1_deltaCR1_plasmidspacer8_4055038_4055138
AATGATGACCGGAAGCATTGGATGGCGCTGTTTAATCTGCTTGAGCAGCGCCAGCCCGTCCATTCCCGGC
ATACGGATATCTGAAAGCAGCACATCCGGCG
>Cse1_deltaCR1_plasmidspacer8_4071318_4071418
GAAATATTTCTCGATAATTTCAATGGTGTAATCGAGCAGCTTGCGCGCTTCCGGGTGACCCGTTGTGACG
GCGCTGGCGGCACCCAGCAGAGCAAAGAAGT
>Cse1_deltaCR1_plasmidspacer8_4105550_4105663
AAGATCAAGCGCCTGTTCCCCATCGTGGGCAACAATCACGTTGAAGCCTTCCATCTCGAGCAGCTCCTTT
AATAGGGAAGTCAGCTCTCGGTCATCATCAACTAACAGGATTTT
>Cse1_deltaCR1_plasmidspacer8_4131432_4131532
GTGGATGCGCGCCCATCCGTATGATGATTTAGTGGTGCTGGACGTTACCGCCAGCCAGCAGCTTGCTGAT
CAGTATCTTGATTTCGCCAGCCACGGTTTCC
>Cse1_deltaCR1_plasmidspacer8_4135480_4135580
TGTACCGTTTGCGCCGGGTCGCGTTGATGCGCGTCAGGATCAGACTGACATTGAGATGTTTGAGCTGCTG
GAGCCAATTGCTGACGGTTTCCGTAACTATC
>Cse1_deltaCR1_plasmidspacer8_4260069_4260174
CCCTTATTGTTGCCGTAATGTTGATTTTCTGTTTTGTAGGTAAGGTGTTATGTTGCCTTGTCGTACCATT
ATCAACACGATAATAATTAATAATTCATTTTTAAAT
>Cse1_deltaCR1_plasmidspacer8_4285398_4285502
CGGGCAATTGTGGGTTACGATGGCATCGCGATAGCCTGCTTCTCTTCAAGCAGCTTCTCGACTACGCCAG
GATCGGCAAGCGTCGAGGTATCGCCCAGGTTGCTG
>Cse1_deltaCR1_plasmidspacer8_4340177_4340277
ATCCAGGCGAATTCATCGACAAGCTCCAGCAGGTCGCGATCCATTGCGGCGAGGGCTTTTTCCCGATCGC
CCAACAGGAAGACCGGCACGTTTTGTTGGCG
>Cse1_deltaCR1_plasmidspacer8_4368394_4368505
TTTTTCGACCGACGATTATCCCCTGCATCGACCGAATACCCGAGATCATATGCTGCTTGAGGATTTCTAC
CGTAATCTGGATCACTTTAAGTGTCGGTTTTTACCCCTTAAT
>Cse1_deltaCR1_plasmidspacer8_4413318_4413422
CTGTATGGACGGATGGATTTTGCCTGGTGTGGCAATGCGCCGGTGAAGCTGCTGGAGTACAACGCCGATA
CGCCAACTTCATTGTACGAGTCGGCTTATTTCCAG
>Cse1_deltaCR1_plasmidspacer8_4453605_4453738
GATGATCACCATCGGCGTCTTTGTGTTGGGTTATCTTTACTGCCTGACCCAGTTTCCCGGTTTTGCTTCC
ACAAGAGTGATCTGCAATATCCTGACCGATAATGCCTTTCTTGGGATCATTGCCGTTGGCATGA
>Cse1_deltaCR1_plasmidspacer8_4461299_4461418

GCTTTCACGCCTTCGAGACGAGCGATACGGGTCATCAGCGCCTTACGTGCCAGCACCAGACGTTCATCCA
GCAGCTTCCAGAAGGTGGCTTCATCGCCTTTTGCTTCCAGAGCAATACGC
>Cse1_deltaCR1_plasmidspacer8_4478280_4478380
TCAGGAAATGCTTATGATAAAACCCGGACATAGATCCCTCCTGTGGCTAACGCCTCAATGAATTAAAATT
CAATTTATATGGATGATTATTCATTTGCAAG
>Cse1_deltaCR1_plasmidspacer8_4530360_4530460
CGCATGGAGGTTACAATAAGGTCCATACCGTTACAGTTAAGAGGGATCTCATTCAGACAGCATTGGGCGG
TAGTGGCTGAAATGCCCTGTTCAGTCAGGAA
