## Supplementary Materials for "Determining the specificity of Cascade binding, interference, and primed adaptation *in vivo* in the *Escherichia coli* type I-E CRISPR-Cas system"

**Table S4. Numbers of potential off-target chromosomal binding sites for spacers in the CRISPR-I array.**

| | 5 bp[a] | 6 bp[a] | 7 bp[a] | 8 bp[a] | 9 bp[a] | 10 bp[a] |
|---|---|---|---|---|---|---|
| **Spacer 1** | 112 | 35 | 18 | 6 | 1 | 0 |
| **Spacer 2** | 157 | 41 | 7 | 3 | 1 | 0 |
| **Spacer 3** | 16 | 3 | 0 | 0 | 0 | 0 |
| **Spacer 4** | 87 | 7 | 2 | 0 | 0 | 0 |
| **Spacer 5** | 181 | 53 | 13 | 3 | 0 | 0 |
| **Spacer 6** | 183 | 27 | 13 | 4 | 2 | 0 |
| **Spacer 7** | 36 | 13 | 5 | 0 | 0 | 0 |
| **Spacer 8** | 232 | 133 | 46 | 14 | 5 | 0 |
| **Spacer 9** | 33 | 2 | 0 | 0 | 0 | 0 |
| **Spacer 10** | 57 | 18 | 2 | 1 | 1 | 1 |
| **Spacer 11** | 207 | 65 | 15 | 4 | 0 | 0 |
| **Spacer 12** | 152 | 54 | 10 | 1 | 1 | 1 |
| **Spacer 13** | 123 | 32 | 3 | 1 | 1 | 0 |

| | 7 bp[a,b] | 8 bp[a,b] | 9 bp[a,b] | 10 bp[a,b] |
|---|---|---|---|---|
| **Spacer 1** | 25 | 7 | 1 | 0 |
| **Spacer 2** | 28 | 8 | 5 | 1 |
| **Spacer 3** | 4 | 0 | 0 | 0 |
| **Spacer 4** | 23 | 6 | 3 | 0 |
| **Spacer 5** | 35 | 11 | 2 | 0 |
| **Spacer 6** | 43 | 11 | 4 | 1 |
| **Spacer 7** | 8 | 2 | 0 | 0 |
| **Spacer 8** | 74 | 23 | 7 | 1 |
| **Spacer 9** | 5 | 0 | 0 | 0 |
| **Spacer 10** | 5 | 2 | 1 | 1 |
| **Spacer 11** | 70 | 25 | 10 | 1 |
| **Spacer 12** | 46 | 14 | 7 | 1 |
| **Spacer 13** | 16 | 5 | 3 | 0 |

[a] Number of chromosomal loci with an AAG PAM flanked by an identical sequence match to the specific length from the start of the indicated spacer.
[b] Number of chromosomal loci with an AAG PAM flanked by an identical sequence match to the specific length from the start of the indicated spacer, allowing for a mismatch at position 6.
