## Supplementary Materials for "Determining the specificity of Cascade binding, interference, and primed adaptation *in vivo* in the *Escherichia coli* type I-E CRISPR-Cas system"

**Table S5. Strains, Plasmids, Oligonucleotides, and Chemically Synthesized dsDNA fragments used in this study.**

Strains

| **Name** | **Description** | **Source** |
|---|---|---|
| MG1655 | MG1655 (F− λ− Δ*ilvG rfb*-50 *rph*-1) | (1) |
| CB386 | MG1655 [Δ*cas3* P*cse1*]::[*cat* P$_{J23119}$] | (2) |
| MLS1003 | MG1655 *araB*::T7pol_*tetA* Δ*araA* Δ*cas3* ΔCRISPR-I ΔCRISPR-II *galK*::J23101_L-II_RII_link_SYFP_op | (3) |
| AMD688 | MLS1003 [Δ*cas3* P*cse1*]::[ P$_{J23119}$] | This Study |
| AMD536 | MG1655 [Δ*cas3* P*cse1*]::[ P$_{J23119}$] | This Study |
| AMD543 | CB386 Cse1-FLAG$_3$ | This Study |
| AMD554 | CB386 FLAG$_3$-Cas5 | This Study |
| LC060 | AMD536 Cse1-FLAG$_3$ ΔCRISPR-II | This Study |
| LC074 | AMD536 ΔCRISPR-I | This Study |
| LC077 | LC074 Cse1-FLAG$_3$ | This Study |
| AMD566 | AMD536 Cse1-FLAG$_3$ | This Study |
| LC099 | AMD566 *yggX** | This study |
| LC103 | AMD536 Δ*yggX*::*kan*$^R$ | This Study |
| LC106 | LC103 Δ*cas1* | This Study |

Plasmids

| **Name** | **Description** | **Source** |
|---|---|---|
| pBAD24 amp | Empty pBAD24 | (4) |
| pCB380 | pcrRNA.con-*lacZ* | (2) |
| pCB381 | pcrRNA.con-*araB* | (2) |
| pAMD172 | Plasmid with synthesized DNA Fragment | This Study |
| pAMD179 | Parent vector for cloning crRNAs | This Study |

| | | |
|---|---|---|
| pLC008 | pAMD179 expressing wild-type sp1.8 | This Study |
| pLC010 | pAMD179 expressing mutant sp1.8 | This Study |
| pAMD189 | pAMD179 expressing a self-targeting crRNA | This Study |
| pLC021 | pBAD24 with protospacer matching the off-target site from *yggX* (includes AAG PAM) | This Study |
| pLC022 | pBAD24 with protospacer that is a perfect match to sp1.8, with an AAG PAM | This Study |
| pBAD33 Cam | pBAD33 Cam | (4) |
| pAMD191 | pBAD33-*cas3* | This Study |
| pLC020 | "Pre-protospacer" Plasmid | This Study |
| pLC023 | Derivative of pLC020 with optimal protospacer matching sp1.8 (variant i) | This Study |
| pLC024 | Derivative of pLC020 with protospacer matching sp1.8 with mismatches across positions 25-32 (variant ix) | This Study |
| pLC025 | Derivative of pLC020 with protospacer matching sp1.8 with mismatches across positions 19-32 (variant x) | This Study |
| pLC026 | Derivative of pLC020 with protospacer matching sp1.8 with mismatches across positions 1-6 (variant xi) | This Study |
| pLC027 | Derivative of pLC020 with protospacer matching sp1.8 with a CCG PAM (variant ii) | This Study |
| pLC028 | Derivative of pLC020 with protospacer with mismatches across positions 1-6 and 25-32 (variant xii) | This Study |
| pLC029 | Derivative of pLC020 with protospacer matching sp1.8 with an ATT PAM (variant iii) | This Study |
| pLC030 | Derivative of pLC020 with protospacer with mismatches across positions 7-24 (variant xiii) | This Study |
| pLC031 | Derivative of pLC020 with protospacer matching sp1.8 with seed mismatches (GGT; variant iv) | This Study |

| | | |
|---|---|---|
| pLC032 | Derivative of pLC020 with protospacer matching sp1.8 with seed mismatches (CGC; variant vii) | This Study |
| pLC033 | Derivative of pLC020 with protospacer matching sp1.8 with seed mismatches (GTC; variant v) | This Study |
| pLC034 | Derivative of pLC020 with protospacer matching sp1.8 with seed mismatches (CCT; variant viii) | This Study |
| pLC035 | Derivative of pLC020 with protospacer matching sp1.8 with seed mismatches (TTT; variant vi) | This Study |
| pCP20 | pPC20 | (5) |
| pLC057 | pBAD24 with protospacer that is a perfect match to CRISPR-I sp1.2, with an AAG PAM | This Study |

<u>Oligonucleotides and Chemically Synthesized dsDNA fragments</u>

| **Name** | **Sequence** |
|---|---|
| JW6272 | GTTAAAATCCTGGGTTCGTAATAATGGCGAGGCGTGAACATGAGAGGCGGTGGCGACTAC |
| JW6273 | CCCCCAGGCTTGCATTGGCCCAGCAAGCCGCAAGATCAAATAAGACGCCGCCTTGTCATC |
| JW6364 | GCGCTTGCCCGCGCCACGCTATACAAACATTTACGGGAGTTAAAAGGCGGTGGCGACTAC |
| JW6365 | GCATCAATTTCATCAGCCATTTGATGGCCCTCCTTGCGGGTTGGGAGCTCACTACTTGTC |
| JW6421 | GTTTTTTTGGGCTAGCGAGTTC |
| JW6513 | CCCGTTTTTTTTGGGCTAGGGAGTTC |
| JW6518 | CAGCGGGGATAAACC |
| JW7490 | AAACCATGCTGATTAATGAAA |
| JW7491 | TCGATATGCACCTCTTTACC |

| | |
|---|---|
| JW7529 | TTTATGGGAAAAAATGCTTTAAGAACAAATGTATACTTTTAGATAGACAGCTGCATGCAT |
| JW7530 | GCGGGGAACACCAGCGTCAGGCGTGAAATCTCACCGTCGTTGCGTGTAGGCTGGAGCTG |
| JW7537 | ACTGGCTTAAAAAATCATTAATTAATAATAGGTTATGTTTAGATAGACAGCTGCATGCAT |
| JW7538 | TGCTAATATAAAAACTTGAGAAAGAGATAACGGGTTATATGGTGTGTAGGCTGGAGCTG |
| JW7539 | ATCATTAATTAATAATAGGTTATGTTTAGAACCATATAACCCGTTATCTCTTTCTCAAGT |
| JW7540 | ACTTGAGAAAGAGATAACGGGTTATATGGTTCTAAACATAACCTATTATTAATTAATGAT |
| JW7598 | GTTTTTTTGGGCTAGGGAGTTCCCCGCGCCAGCGGGGATAAACCG |
| JW7635 | GATTAATGAAAAGAAACTCAACATGATGAATGCCGAGCACCGCTAGACAGCTGCATGCAT |
| JW7636 | ATCTTCCGGCGTATAGCCCTCGATATGCACCTCTTTACCCTCGGTGTAGGCTGGAGCTG |
| JW7637 | GATGAATGCCGAGCACCGCAAGTGACTTGAGCAGGAGATGGTCAAC |
| JW7638 | GTTGACCATCTCCTGCTCAAGTCACTTGCGGTGCTCGGCATTCATC |
| JW7693 | GCGCGGGGAACTCGACTGGTGAGTACTCAACCAAGTCATTCTGAGCGGTTTATCCCCGC |
| JW7736 | GGGCTAGCGAATTCGAAAAACAGGGAGGCTATTAAT |
| JW7738 | GATCCCCGGGTACCGTTATTTGGGATTTGCAGGGA |
| JW7818 | TCGTCGGCAGCGTCAGATGTGTATAAGAGACAGAAAGTTGGTAGATTGTGACTGGC |
| JW7819 | GTCTCGTGGGCTCGGAGATGTGTATAAGAGACAGCAACAGCAGCACCCATGAC |
| JW7898 | TGGCTTGCTATCTTTGGCTCCACTGTGATTGAGGTGTAATAAATAGACAGCTGCATGCAT |
| JW7899 | GAGGTACATTTTCAGTGACCACGACCAACATACTCATTGTGTAGGCTGGAGCTG |
| JW7900 | TCCACTGTGATTGAGGTGTAATAAAAATGAGTATGTTGGTCGTGGTCG |
| JW7901 | CGACCACGACCAACATACTCATTTTTATTACACCTCAATCACAGTGGA |
| JW7911 | GCGCGGGGAACTCGAGGCGGCTTGCCTTGCAGCCAGCTCCAGCAGCGGTTTATCCCCGC |
| JW7912 | GCGCGGGGAACTCGAGGCGGCTTGCCTTGCAGCCAGCTCCAGTCACGGTTTATCCCCGC |
| JW7938 | TGCGACGCTGGCGATATCTGAATGCCGAGCACCGCA |
| JW7939 | AAACAGCCAAGCTTGCATGCTGCACCTCTTTACCCTCG |
| JW7922 | ATGCACGAAATAACCCTCT |
| JW7923 | TCACATACTGTTGGCATGTT |
| JW7940 | TGCGACGCTGGCGATATCCGAGCACCGCAAGCTGCTGGAGCTGGCTGCAAGGCAAGCCGC |
| JW7941 | AAACAGCCAAGCTTGCATGTGCACCTCTTTACCCTCGTGGGCGGCTTGCCTTGCAGCCAG |

| | |
|---|---|
| JW8040 | CTAGCAGGAGGAATTCTGATTTTATCGCACCACTC |
| JW8042 | CGTCGGTTTTTTTACCCTC |
| JW8043 | GTACCATGGTGAATTTCGCTCATTTCGACTTCGG |
| JW8128 | GTAAAAAAACCGACGGAATTCGAAACGTGTTGCTGTGGG |
| JW8129 | CGTTTCCTGAGAATTNNNNNGTCTCGACTGAGGAACCATGAAACAGTATTTAGAACTG |
| JW8130 | AAAACCGACGGAATTGAAGCTGCTGGAGCTGGCTGCAAGGCAAGCCGCCCAACACACTGGCGTCGGCTCT |
| JW8139 | AAAACCGACGGAATTGAAGCTGCTGGAGCTGGCTGCAAGGCAAAGTATATGCCACACTGGCGTCGGCTCT |
| JW8145 | AAAACCGACGGAATTGAAGCTGCTGGAGCTGGCTGCAGACTTCAGTATATGCCACACTGGCGTCGGCTCT |
| JW8169 | AAAACCGACGGAATTGAAGTGATGCGAGCTGGCTGCAAGGCAAGCCGCCCAACACACTGGCGTCGGCTCT |
| JW8499 | AAAACCGACGGAATTGCCGCTGCTGGAGCTGGCTGCAAGGCAAGCCGCCCAACACACTGGCGTCGGCTCT |
| JW8500 | AAAACCGACGGAATTGAAGTGATGCGAGCTGGCTGCAAGGCAAAGTATATGACACACTGGCGTCGGCTCT |
| JW8501 | AAAACCGACGGAATTGATTCTGCTGGAGCTGGCTGCAAGGCAAGCCGCCCAACACACTGGCGTCGGCTCT |
| JW8502 | AAAACCGACGGAATTGAAGCTGCTGACTGATATCATTGACTTCGCCGCCCAACACACTGGCGTCGGCTCT |
| JW8537 | CAAGCAGAAGACGGCATACGAGATGGATTCACGTCTCGTGGGCTCGGAGATGTGTATAAGAGACAGCGGAGGGTAAAAAAACCGACGGAATT |
| JW8556 | CAAGCAGAAGACGGCATACGAGATGCTAGACTGTCTCGTGGGCTCGGAGATGTGTATAAGAGACAGCGGAGGGTAAAAAAACCGACGGAATT |
| JW8557 | CAAGCAGAAGACGGCATACGAGATCTCAGGTAGTCTCGTGGGCTCGGAGATGTGTATAAGAGACAGCGGAGGGTAAAAAAACCGACGGAATT |
| JW8558 | CAAGCAGAAGACGGCATACGAGATCGCATTAGGTCTCGTGGGCTCGGAGATGTGTATAAGAGACAGCGGAGGGTAAAAAAACCGACGGAATT |
| JW8559 | CAAGCAGAAGACGGCATACGAGATTGTCCAAGGTCTCGTGGGCTCGGAGATGTGTATAAGAGACAGCGGAGGGTAAAAAAACCGACGGAATT |
| JW8561 | CAAGCAGAAGACGGCATACGAGATGGCTATCAGTCTCGTGGGCTCGGAGATGTGTATAAGAGACAGCGGAGGGTAAAAAAACCGACGGAATT |

| | |
|---|---|
| JW8562 | CAAGCAGAAGACGGCATACGAGATAACTCGTGGTCTCGTGGGCTCGGAGATGTGTATAAGAGACAGCGGAGGGTAAAAAAACCGACGGAATT |
| JW8563 | CAAGCAGAAGACGGCATACGAGATATGGACTCGTCTCGTGGGCTCGGAGATGTGTATAAGAGACAGCGGAGGGTAAAAAAACCGACGGAATT |
| JW8564 | CAAGCAGAAGACGGCATACGAGATCATATGGCGTCTCGTGGGCTCGGAGATGTGTATAAGAGACAGCGGAGGGTAAAAAAACCGACGGAATT |
| JW8565 | CAAGCAGAAGACGGCATACGAGATTAAGGCGAGTCTCGTGGGCTCGGAGATGTGTATAAGAGACAGCGGAGGGTAAAAAAACCGACGGAATT |
| JW8566 | CAAGCAGAAGACGGCATACGAGATCGTACTAGGTCTCGTGGGCTCGGAGATGTGTATAAGAGACAGCGGAGGGTAAAAAAACCGACGGAATT |
| JW8567 | AATGATACGGCGACCACCGAGATCTACACTCCAGGTATCGTCGGCAGCGTCAGATGTGTATAAGAGACAGGACTTTGAGATTGAAGGCTACGATCCG |
| JW8675 | AAAACCGACGGAATTGAAGGGTCTGGAGCTGGCTGCAAGGCAAGCCGCCCAACACACTGGCGTCGGCTCT |
| JW8676 | AAAACCGACGGAATTGAAGCGCCTGGAGCTGGCTGCAAGGCAAGCCGCCCAACACACTGGCGTCGGCTCT |
| JW8677 | AAAACCGACGGAATTGAAGGTCCTGGAGCTGGCTGCAAGGCAAGCCGCCCAACACACTGGCGTCGGCTCT |
| JW8678 | AAAACCGACGGAATTGAAGCCTCTGGAGCTGGCTGCAAGGCAAGCCGCCCAACACACTGGCGTCGGCTCT |
| JW8679 | AAAACCGACGGAATTGAAGTTTCTGGAGCTGGCTGCAAGGCAAGCCGCCCAACACACTGGCGTCGGCTCT |
| 144148263 (synthesized dsDNA fragment) | CAGACATTTGGGTTAAACAGGCGTACCCCGGTAGATTTGGATGGTTTAAGGTTGGTGTCTTTTTTACCTGTTTGAAAACAAAGAATTAGCTGATCTTTAATAATAAGGAAATGTTACATTAAGGTTGGTGGGTTGTTTTTATGGGAAAAAATGCTTTAAGAACAAATGTATACTTTTAGAGAGTTCCCCGCGCCAGCGGGGATAAACCGCAGCTCCCATTTTCAAACCCATCAAGACGCCTTCGCCAACTCCTTCACCAGAGGTAGCATTATCCGCATAACGTCACGGCAGCGACGTTCTATTCTTCCAGGAAGAGCCTTATCAATATGTTGGTGATTATCCAGTCTTACGTCATGCCAGCTATTTCCCGCCGGGAAGGCAGGTGTTTTTGCGCG |
| Chemically synthesized insert from pAMD172 | TCGTAATACGACTC_CTATAGACAACGGTCAGGAGAAGGAGGACGGCATGTACCCATACGACGTCCCAGACTACGCTAGTACTGACTACAAGGATCACGACTACAAGGACCACGACTATAAAGACCACGACGCAATGTAGGTTTTTTTGGGCTAGCGAGTTCCCCGCGCCAGCGGGGATAAACCGCGGAATAATAATAATCTCGAGTTCCCCGCGCCAGCGGGGATAAACCGAGGGAACTGCCAGGCATCAAATAAAACGAAAGGCTCAGTCGAAAGACTGGGCCTTTCGTTTTATCTGTTGTTTGTCGGTGAACGCTCTCCTGAGTAGGACAAGCTTGGCTGTTTT |

| | |
|---|---|
| | GGCATAATACGACTCACTATAGTGAACGGTCTCCCCGTCCTC_ACGGCGACTACCCATACGACGTCCCAGACTACGCTAGTACTGACTACAAGGATCACGACTACAAGGACCACGACTATAAAGACCACGACGCAATGTA_TCACACTGG |
| JW9131 | TGCGACGCTGGCGATATCCGAGCACCGCAAGCAGCCGAAGCCAAAGGTGATGCCGAACAC |
| JW9132 | AAACAGCCAAGCTTGCATGTGCACCTCTTTACCCTCGAGCGTGTTCGGCATCACCTTTG |

### REFERENCES


1. Blattner FR, Plunkett G, Bloch CA, Perna NT, Burland V, Riley M, Collado-Vides J, Glasner JD, Rode CK, Mayhew GF, Gregor J, Davis NW, Kirkpatrick HA, Goeden MA, Rose DJ, Mau B, Shao Y. 1997. The complete genome sequence of Escherichia coli K-12. Science 277:1453–1462.

2. Luo ML, Mullis AS, Leenay RT, Beisel CL. 2014. Repurposing endogenous type I CRISPR-Cas systems for programmable gene repression. Nucleic Acids Res gku971.

3. Amlinger L, Hoekzema M, Wagner EGH, Koskiniemi S, Lundgren M. 2017. Fluorescent CRISPR Adaptation Reporter for rapid quantification of spacer acquisition. Sci Rep 7:10392.

4. Guzman LM, Belin D, Carson MJ, Beckwith J. 1995. Tight regulation, modulation, and high-level expression by vectors containing the arabinose PBAD promoter. J Bacteriol 177:4121–4130.

5. Cherepanov PP, Wackernagel W. 1995. Gene disruption in Escherichia coli: TcR and KmR cassettes with the option of Flp-catalyzed excision of the antibiotic-resistance determinant. Gene 158:9–14.
