## Supplementary figures and images for "Determining the specificity of Cascade binding, interference, and primed adaptation *in vivo* in the *Escherichia coli* type I-E CRISPR-Cas system"

### Supplementary Materials

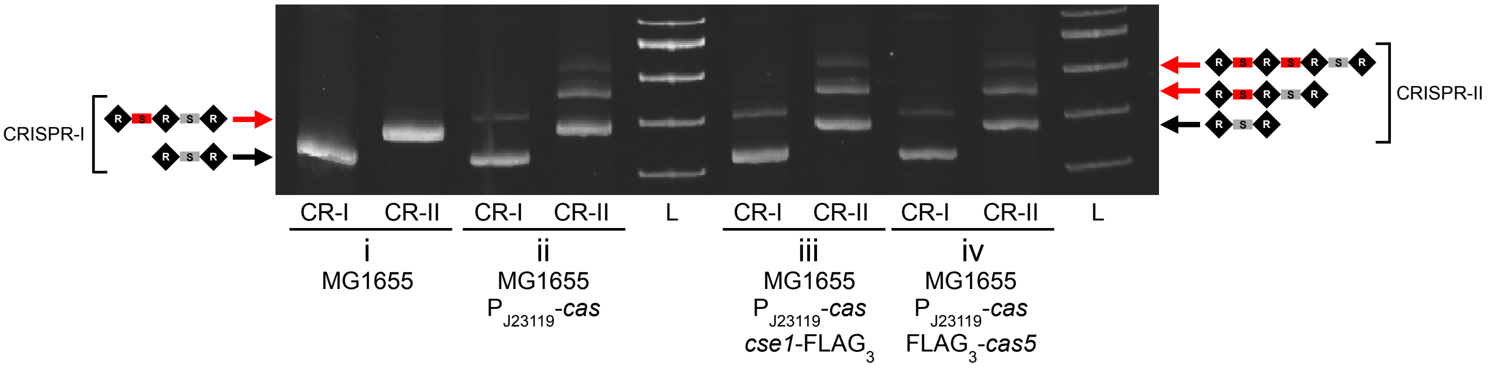

### Supplementary Materials

#### Figure S3

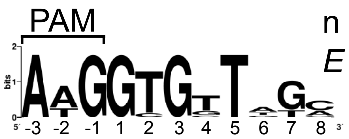


n = 10

*E* = 5.3e-1

**GTGTTTGCG... CRISPR-II Spacer #2**

### Supplementary Materials

#### Figure S4

**A**

n = 64
*E* = 9.4e$^{-136}$

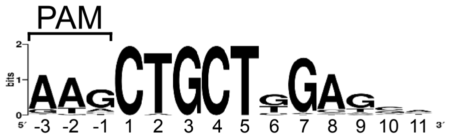


**sp1.8 CTGCTGGAGCT...**

**B**

n = 14
*E* = 6.8e$^{-3}$

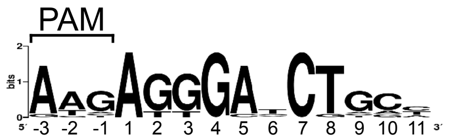


**AGGGAACTGCC...**
