## Supplementary Materials for "Determining the specificity of Cascade binding, interference, and primed adaptation *in vivo* in the *Escherichia coli* type I-E CRISPR-Cas system"

### Figure S2

A

*lacZ*-targeting crRNA spacer: 5′ CTTTACACTTTATGCTTCCGGCTCGTATGT 3′

B

*araB*-targeting crRNA spacer: 5′ ATTAGCGGATCCTACCTGACGCTTTTTATC 3′

C

CRISPR-I

**GAGTTCCCCGCGCCAGCGGGGATAAACCG**CTTTCGCAGACGCGCGGCGATACGCTCACGCA**GAG**
**TTCCCCGCGCCAGCGGGGATAAACCG**CAGCCGAAGCCAAAGGTGATGCCGAACACGCT**GAGTTC**
**CCCGCGCCAGCGGGGATAAACCG**GGCTCCCTGTCGGTTGTAATTGATAATGTTGA**GAGTTCCCC**
**GCGCCAGCGGGGATAAACCG**TTTGGATCGGGTCTGGAATTTCTGAGCGGTCGC**GAGTTCCCCGC**
**GCCAGCGGGGATAAACCG**CGAATCGCGCATACCCTGCGCGTCGCCGCCTGC**GAGTTCCCCGCGC**
**CAGCGGGGATAAACCG**TCAGCTTTATAAATCCGGAGATACGGAAACTA**GAGTTCCCCGCGCCAG**
**CGGGGATAAACCG**GACTCACCCCGAAAGAGATTGCCAGCCAGCTT**GAGTTCCCCGCGCCAGCGG**
**GGATAAACCG**CTGCTGGAGCTGGCTGCAAGGCAAGCCGCCCA**GAGTTCCCCGCGCCAGCGGGGA**
**TAAACCG**GGGGGCGCATGACCGTAAACATTATCCCCCGG**GAGTTCCCCGCGCCAGCGGGGATAA**
**ACCG**GGAGTTCAGACATAGGTGGAATGATGGACTAC**GAGTTCCCCGCGTTAGCGGGATAAACC**
**G**CCCGGTAGCCAGGTTTGCAACGCCTGAACCGA**GAGTTCCCCGCGCCAGCAGGGATAAACCG**GC
AACGACGGTGAGATTTCACGCCTGACGCTG**GTGTTCCCCGCATCAGCGGGGATAAACCG**GGCGC
ACTGGATGCGATGATGGATATCACTTG**GAGTTCCCCCGCCTCTGCGGTAGAACTCC**

D

CRISPR-II

**GTGTTCCCCGCGCCAGCGGGGATAAACCG**GCAAAAACCGGGCAATCGCAAAAAGGCGTAAT**GTG**
**TTCCCCGCGCCAGCGGGGATAAACCT**GTGTTTGCGGCATTAACGCTCACCAGCATTTC**GTGTTC**
**CCCGCGCCAGCGGGGATAAACCG**ACGTGGTCATGGGTGCTGCTGTTGCAGAGCCA**GTGTTCCCC**
**GCGCCAGCGGGGATAAACCG**AGCAGATACACGGCTTTGTATTCCGTGCGCCC**GTGTTCCCCGCG**
**CCAGCGGGGATAAACCG**AATAGCAATAGTCCATAGATTTGCGAAAACAG**GTGTTCCCCGCGCCA**
**GCGGGGATAAACCG**GAGCCTGACGAGACTACTGAGGCCGTTCTGTC**GAGTTCCCCGCGCCAGCG**
**GGGATAAACCA**

E

Portion of CRISPR-I spacer #8-expressing crRNA plasmid

**GAGTTCCCCGCGCCAGCGGGGATAAACCG**CTGCTGGAGCTGGCTGCAAGGCAAGCCGCCTC**GAG**
**TTCCCCGCGCCAGCGGGGATAAACCG**AGGGAACTGCCAGGCATCAAATAAAACGAAAGGCTCAG
TCGAAAGACTGGGCCTTTCGTTTT
